# A genetic program boosts mitochondrial function to power macrophage tissue invasion

**DOI:** 10.1101/2021.02.18.431643

**Authors:** Shamsi Emtenani, Elliott T. Martin, Attila Gyoergy, Julia Bicher, Jakob-Wendelin Genger, Thomas R. Hurd, Thomas Köcher, Andreas Bergthaler, Prashanth Rangan, Daria E. Siekhaus

## Abstract

Metabolic adaptation to changing demands underlies homeostasis. During inflammation or metastasis, cells leading migration into challenging environments require an energy boost, however what controls this capacity is unknown. We identify a previously unstudied nuclear protein, Atossa, as changing metabolism in *Drosophila melanogaster* immune cells to promote tissue invasion. Atossa’s vertebrate orthologs, FAM214A-B, can fully substitute for Atossa, indicating functional conservation from flies to mammals. Atossa increases mRNA levels of Porthos, an unstudied RNA helicase and two metabolic enzymes, LKR/SDH and GR/HPR. Porthos increases translation of a gene subset, including those affecting mitochondrial functions, the electron transport chain, and metabolism. Respiration measurements and metabolomics indicate that Atossa and Porthos powers up mitochondrial oxidative phosphorylation to produce sufficient energy for leading macrophages to forge a path into tissues. As increasing oxidative phosphorylation enables many crucial physiological responses, this unique genetic program may modulate a wide range of cellular behaviors beyond migration.

## INTRODUCTION

Charged with protecting the organism against continuously changing threats, the immune system must constantly adapt, altering the location, number, and differentiation status of its different immune cell subtypes (Nicholson, 2016). Such continuous adjustment comes at a cost, as it requires high levels of energy. However, how immune cells adjust their metabolic capacities to achieve these increased metabolic requirements is just beginning to be understood (Guak et al., 2020; O’Neill et al., 2016). The main energy currency in the cell is ATP. The conversion of carbohydrates into ATP is mediated mostly by cytoplasmic glycolysis and the mitochondrial TCA cycle that feeds electron donors into oxidative phosphorylation (OxPhos) complexes I through IV. Anaerobic glycolysis is quick and does not require oxygen, but respiratory OxPhos extracts considerably more ATP from a single molecule of glucose, albeit more slowly (Berg et al., 2002). Amino acids and fatty acids also feed into the TCA cycle and fuel OxPhos (O’Neill et al., 2016). OxPhos is most directly regulated by the activity and the amount of complexes I through V that carry it out (Hüttemann et al., 2007), but can also be affected by mitochondrial fusion (Rambold et al., 2015) and biogenesis (Le Bleu et al., 2014). Upregulation of OxPhos is known to be required for many important immune cell functions, such as B cell antibody production (Price et al., 2018), pathogenic T cell differentiation during autoimmunity (Shin et al., 2020), and CD8+ memory T cell development and expansion (van der Windt et al., 2012; van der Windt et al., 2013), T reg suppressive function (Angelin et al., 2017; Weinberg et al., 2019; Beir et al., 2015) and the maturation of anti-inflammatory macrophages (Vats et al., 2006). However, what genetic programs immune cells utilize to upregulate OxPhos remains unclear and how such shifts in metabolism could influence immune cell migration is unexplored.

Immune cells move within the organism to enable their distribution and maturation (Kierdorf et al., 2015; Masopust and Schenkel, 2013), as well as to detect and respond to homeostatic challenges, injuries, tumors or infections (Woodcock et al., 2015; Luster et al., 2005; Ratheesh et al, 2015). To migrate across unimpeded environments cells expend energy to restructure their own actin cytoskeleton, activate myosin ATPase, and reorganize their cell membrane (Bernstein and Bamburg, 2003; Cuvelier et al., 2007; Rottner and Schaks, 2019; Li et. al, 2019). Even greater energy requirements exist when cells must also remodel their surroundings as they move ahead against the resistance of flanking cells or extracellular matrix (Zanotelli et al., 2018; Zanotelli et al., 2019, Cunniff et al., 2016; Kelley et al., 2019). Most studies on the metabolism that enables immune cell migration *in vitro* or *in vivo* have highlighted the importance of glycolysis, in macrophages, dendritic cells and regulatory T cells (Guak et al., 2018; Kishore et al., 2018; Semba et al., 2016; Liu et al., 2019). To our knowledge, only one study has demonstrated a need for a functional electron transport chain (ETC) to speed neutrophil migration *in vivo (*Zhou et al., 2018) potentially through polarized secretion of ATP to amplify guidance cues (Bao et al., 2015). In cancer cells an increase in the transcription of mitochondrial genes, mitochondrial biogenesis and thus OxPhos by PGC-1 appears to underlie enhanced invasion and metastasis (Le Bleu et al., 2014). OxPhos is particularly required in the first cancer cell leading coordinated chains into challenging environments *in vitro* (Khalil et al., 2010; Commander et al., 2020); these leader cells have been shown to need higher ATP levels to create a path (Zhang et al., 2019). Although the ability of immune cells to invade tissues or tumors also depends on movement against surrounding resistance, it is not known if immune cells similarly require OxPhos for such infiltration.

To identify new mechanisms governing *in vivo* migration, we study *Drosophila* macrophages, also called plasmatocytes. Macrophages are the primary phagocytic and innate immune cell in *Drosophila* and share remarkable similarities with vertebrate macrophages in ontogeny, functional properties, and migratory behavior (Brückner et al., 2004; Nourshargh and Alon, 2014; Ratheesh et al., 2015; Weavers et al., 2016; Wood and Martin, 2017; Weavers et al., 2020). Phagocytic macrophages not only resolve infections, but also influence development and homeostasis (Caputa et al., 2019; Riera-Domigo et al., 2020; Buck et al., 2016; Bunt et al., 2010). Embryonic *Drosophila* macrophages follow guidance cues to disseminate along predetermined routes (Cho et al., 2002; Brückner et al., 2004; Wood et al., 2006) from their initial site of specification. During embryogenesis a dynamic chain of macrophages invades into the extended germband between the closely apposed ectoderm and mesodermal tissues, moving against the resistance of surrounding tissues (Siekhaus et al., 2010; Ratheesh et al., 2018; Valoskova et al., 2019). Importantly, the rate limiting step for this tissue invasion is the infiltration of the pioneer macrophage, a process affected both by the properties of the surrounding tissues (Ratheesh et al., 2018) as well as macrophages themselves (Valoskova et al., 2019; Belyaeva et al., 2021).

Here we identify a program that powers the invasive capability of these pioneer macrophages *in vivo*. We characterize a metabolic shift orchestrated in these immune cells by a single previously unexamined nuclear factor that we name Atossa. We show that Atossa induces higher mRNA levels of two metabolic enzymes and a previously unstudied helicase. This helicase which we name Porthos enhances translation of a diverse set of proteins, including those affecting mitochondrial and metabolic function, to increase OxPhos and ATP. Our work thus reveals a detailed cellular mechanism that induces concerted metabolic and mitochondrial reprogramming to support higher energy levels. Given that we find that Atossa’s mammalian orthologs maintain its function, our data lay the foundation for mammalian studies on diverse pathological conditions, from autoimmunity to cancer, as well as those independent of invasion.

## RESULTS

### *CG9005* is required in macrophages for their early invasion into the extended germband

To identify unknown molecular pathways mediating germband invasion, we searched for previously uncharacterized genes enriched in macrophages prior to and during germband tissue entry. Examining the BDGP *in situ* project (https://insitu.fruitfly.org/cgi-bin/ex/report.pl?ftype=1&ftext=FBgn0033638) we identified CG9005 as a gene fitting these requirements. CG9005 is enriched in macrophages from their birth through their invasion of the germband. CG9005 is maternally deposited and expressed in all mesodermal cells during stage 4-6 when macrophages are specified in the head mesoderm. CG9005 is further upregulated in macrophages starting at Stage 7 while its expression decreases in the remaining mesoderm. CG9005 continues to be expressed during Stage 9-12 in macrophages, during their ingression, dissemination, and movement towards and into the germband. After invasion, CG9005 is downregulated in macrophages to match the lower expression levels found ubiquitously in the embryo.

We examined a P element insertion allele, *CG9005*^*BG02278*^, henceforth abbreviated to *CG9005*^*PBG*^, visualizing macrophages through expression of a nuclear fluorescent marker. Quantification of the number of macrophages within the germband in fixed embryos at Stage 12 revealed a 36% decrease in *CG9005*^*PBG*^ mutant embryos compared to the control (Figs. 1A-B and 1D), similar to that seen when *CG9005*^*PBG*^ was placed over either *Df(2R)ED2222* or *Df(2R)BSC259* that remove the gene entirely (Fig. 1D), demonstrating that *CG9005*^*PBG*^ is a genetic null for macrophage germband invasion. Expressing CG9005 in macrophages in the mutant completely restored their capacity to invade the germband (Figs. 1C-D). Depleting *CG9005* by driving any one of three independent RNA interference (RNAi) lines in macrophages caused a 37-40% decrease in macrophages within the germband compared to controls (Fig. 1E). We also observed 24-27% more macrophages sitting on the yolk near the entry site that have not yet invaded the germband in *CG9005*^*PBG*^ (Fig. S1A) and in the RNAi lines (Fig. S1B) compared to their controls. This finding supports the conclusion that macrophages in these backgrounds migrate normally up to the germband but are less able to enter. We counted macrophages migrating along the ventral nerve cord (vnc) in late Stage 12 embryos, a route guided by the same factors that lead into the germband (Brückner et al., 2004; Cho et al., 200; Wood et al., 2006) but that does not require tissue invasion (Siekhaus et al., 2010; Weavers et al., 2016). There was no significant difference in both the *CG9005*^*PBG*^ mutant (Fig. S1C) and the *CG9005* RNAi-expressing macrophages (Figs. S1D-F) compared to their controls, arguing that basic migratory processes and recognition of chemotactic signals are unperturbed. Moreover, we detected no significant change in the total number of macrophages for any of these genotypes (Figs. S1G-H). Taken together, these results from fixed embryos clearly suggest that CG9005 is specifically required in macrophages for the early steps of germband invasion.

**Figure 1.**
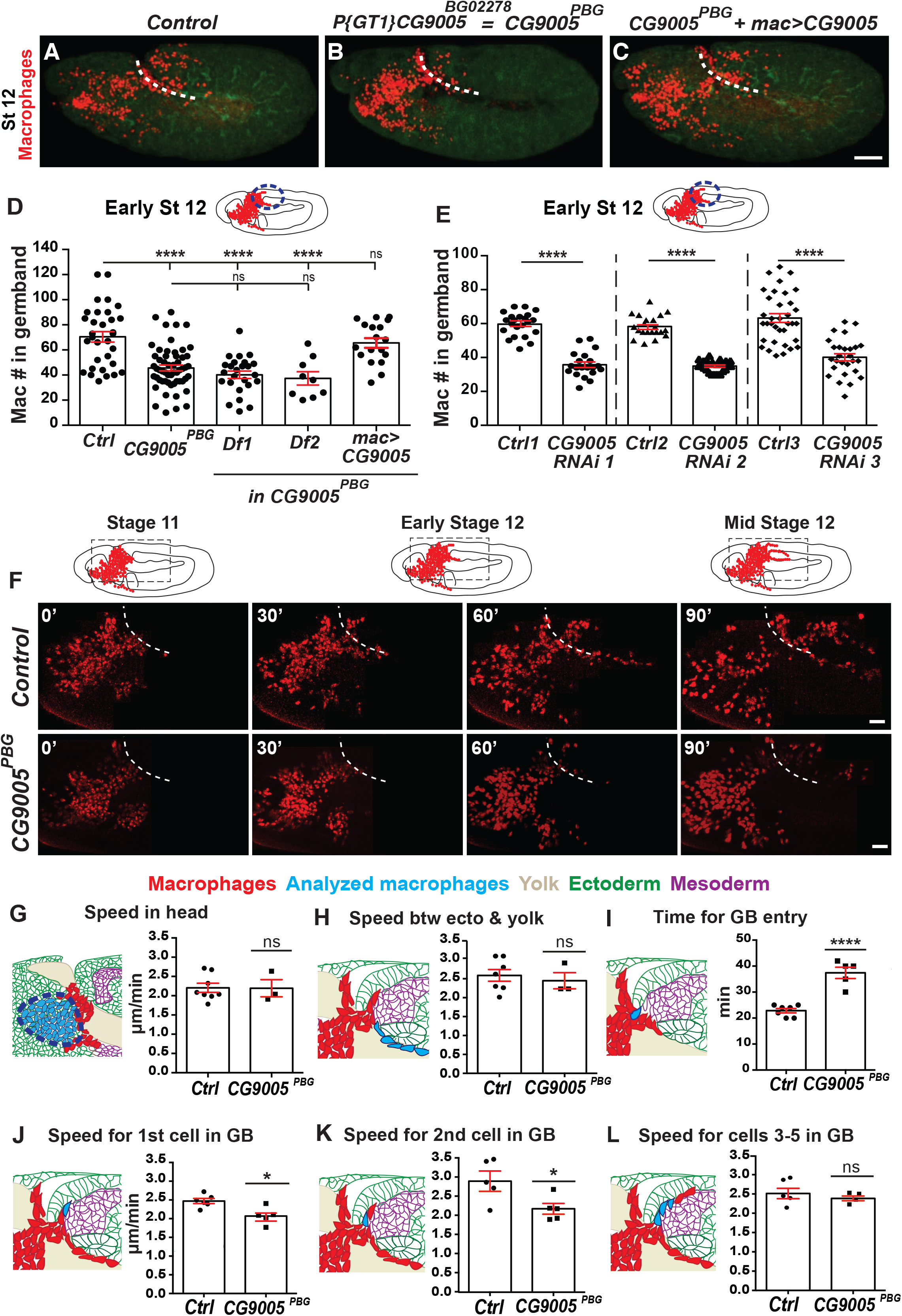
CG9005 acts in macrophages to spur pioneer cell infiltration into the germband tissue. **Fig 1A-C**. Representative confocal images of Stage 12 embryos from the control, the *P{GT1}CG9005*^*BG02278*^ P element mutant (henceforth called *CG9005*^*PBG*^), and *CG9005*^*PBG*^ with *CG9005* expression restored in macrophages. Macrophages (red) and phalloidin to visualize embryo (green). “*mac*” represents the *srpHemo-Gal4* driver. Germband edge indicated by dotted white line. **Fig 1D**. Quantification reveals a significant decrease in the number of macrophages that have penetrated the germband in Stage 12 embryos from *CG9005*^*BG*^ (n=56), and from *CG9005*^*BG*^ over two deficiencies (Df) that completely remove the gene (*CG9005*^*PBG*^*/Df1(2R)* n=25 and *CG9005*^*PBG*^*/Df2(2R)* n=9), compared to the control (n=35). Macrophage expression of *CG9005* rescues the mutant phenotype arguing that *CG9005* is required only in macrophages for germband penetration (n=18 for rescue, p<0.0001 for control vs mutant, p=0.98 for control vs rescue, p=0.001 for mutant vs rescue). *Df1(2R)=Df(2R)ED2222. Df2(2R)=Df(2R)BSC259*. **Fig 1E**. Macrophage specific knockdown of *CG9005* by *UAS* RNAi lines under the control of *srpHemo-GAL4* can recapitulate the mutant phenotype (control 1 n=22, *CG9005 RNAi 1* (VDRC 106589) n=20; p<0.0001; control 2 n=21, *CG9005 RNAi 2* (VDRC 36080) n=23; p<0.0001; control 3 n=35, *CG9005 RNAi 3* (BL33362) n=28, p<0.0001). **Fig 1F**. Stills from two-photon movies of control and *CG9005*^*PBG*^ mutant embryos showing macrophages (nuclei, red) migrating starting at Stage 10 from the head towards the germband and invading into the germband tissue. Elapsed time indicated in minutes. The germband edge (white dotted line) was detected by yolk autofluorescence. **Fig 1G-H**. Quantification shows no change in macrophage migration speed (**G**) in the head or (**H**) between the yolk sac and the germband edge in the *CG9005*^*PBG*^ mutant compared to the control. Head speed: control and mutant=2.2 µm/min; movie #: control=8, mutant=3; track #: control=360, mutant=450, p=0.65. Between yolk sac and germband speed: control=2.6 and mutant=2.4 µm/min; # movies: control=7, mutant =3; # tracks: control=46, mutant=19, p=0.62. **Fig 1I**. The time required for the first macrophage nucleus to enter into the extended germband is increased by 65% in the *CG9005*^*PBG*^ mutant compared to the control (control=22.8 min, n=7, mutant=37.4 min, n=5, p<0.0001). **Fig 1J-K**. The migration speed of the first and second macrophage into the germband between the mesoderm and ectoderm is significantly slower in the *CG9005*^*PBG*^ mutant compared to the control. First macrophage speed: control=2.5 and mutant =2.1 µm/min, movie #: control=6, mutant=5, p=0.012. Second macrophage speed: control=2.9 and mutant=2.2 µm/min, movie #: control=5, mutant=5, p=0.03. **Fig 1L**. The migration speed of the third to fifth macrophage nuclei along the first 25-30 um of the path between the germband mesoderm and ectoderm is similar in the *CG9005*^*PBG*^ mutant and the control (speed: control=2.5 and mutant=2.4 µm/min, movie #: control=5, mutant=4, p=0.17). In schematics, macrophages are shown in red and analyzed macrophages in light blue, the ectoderm in green, the mesoderm in purple, and the yolk in beige. Macrophage nuclei visualized by *srpHemo-H2A::3xmCherry* expression. See also **Fig. S1** and **Videos 1** and **2**. Throughout this work, embryos were staged based on germband retraction away from the anterior of less than 29% for stage 10, 29%-31% for stage 11, and 35%–40% for stage 12. In all figures and histograms show mean±SEM and ns=p>0.05, *p<0.05, **p<0.01, ***p<0.001, ****p<0.0001. One-way ANOVA with Tukey for (**D**-**E**), and unpaired t-test for (**G**-**L**). Scale bars: 50 µm in (**A**-**C**), 30 µm in (**F**). See also **Figure S1** and **Videos 1** and **2**: representative movies of macrophage migration into the germband in the control (**Video 1**) and the *CG9005*^*PBG*^ (*atos*) mutant (**Video 2**). Macrophages (red) are labeled with *srpHemo-H2A::3xmCherry*. Arrow indicates first macrophage moving into the germband. The time interval between each acquisition is 40 s and the display rate is 15 frames/s. Scale bar: 20 μm.

### Atossa (CG9005) promotes efficient invasion of pioneer macrophages into the germband tissue

To directly assess CG9005’s role in germband invasion, we conducted two-photon live imaging. We labeled macrophages with the nuclear marker *srpHemo-H2A::3xmCherry* in control and *CG9005*^*PBG*^ embryos (Figs. 1F and S1I, Videos 1 and 2). We observed no significant change in speed or directionality during macrophage migration from their initial position at Stage 9 in the head mesoderm up to the yolk neighboring the germband entry point in *CG9005*^*PBG*^ (Fig. 1G, Figs. S1J-L) (speed in the head and yolk: 2.2 µm/min for both the control and *CG9005*^*PBG*^; p=0.65, p=0.78 respectively), nor in their directionality within these regions (directionality: 0.39 in control and 0.37 in mutant in both regions, p=0.74 for head, p=0.86 for yolk). We also observed no significant change in migration speed for macrophages moving between the yolk and ectoderm (control=2.6 and *CG9005*^*PBG*^=2.5 µm/min, p=0.62) (Fig. 1H). However, the first macrophage in *CG9005*^*PBG*^ required 65% more time than the control to enter into the germband tissue (time to entry: control=23 min and *CG9005*^*PBG*^=38 min, p<0.0001) (Fig. 1I). The speed of the first two pioneering macrophages is also significantly slower as they invade along the path between the mesoderm and ectoderm in *CG9005*^*PBG*^ mutant embryos compared to the control (1^st^ cell: control=2.5 and *CG9005*^*PBG*^*=*2 µm/min, p=0.012; 2^nd^ cell: control=2.9 and *CG9005*^*PBG*^=2.1 µm/min, p=0.03) (Figs. 1J-K). However, the speed of the next few cells migrating along this path was not affected (3^rd^-5^th^ cells: control=2.5 and *CG9005*^*PBG*^=2.4 µm/min, p=0.17) (Fig.1L). We conclude that CG9005 specifically regulates tissue invasion, facilitating the initial entry into and subsequent movement within the germband tissue of the first two pioneer macrophages. Since the macrophage stream invading the germband becomes a trickle in *CG9005*^*PBG*^ we named the gene *atossa* (*atos*), for the powerful Persian queen whose name means trickling.

### Atossa (CG9005) is a nuclear protein whose conserved motifs and TADs are important for macrophage tissue invasion, a function conserved by its vertebrate orthologs

Atossa (Atos) contains a conserved domain of unknown function (DUF4210) and a Chromosome segregation domain (Chr_Seg) (Fig. 2A). Atos also displays two trans-activating domains (TADs) common among transcription factors as well as three nuclear localization signals (NLS) and a nuclear export signal (NES). We first tested the subcellular distribution of the Atos protein, transfecting the macrophage-like S2R+ cell line with a *FLAG::HA* tagged form of *atos* under the control of the macrophage promoter *srpHemo*. We found Atos mainly in the nucleus colocalized with DAPI, and also partially in the cytoplasm (Fig. S2A). When expressed *in vivo* in macrophages, Atos is also predominantly a nuclear factor (Fig. 2B). To assess the importance of the conserved domains and TADs, we made versions of Atos lacking these regions. All mutant forms were present in the nucleus similarly to wild-type Atos (Fig. S2A). While expression of wild-type Atos in the macrophages of *atos* embryos completely rescues germband invasion (Figs. 2C-D), Atos lacking either the conserved DUF2140, the Chr_Seg domain, or either or both of the two TAD motifs failed to do so (Figs. 2D, S2B-C). Consistent with a germband invasion defect, expression of these Atos mutants led to a higher number of macrophages sitting on the yolk at the germband entry site prior to invasion than in the rescue with wild-type Atos (Fig. S2D). These data clearly show that the conserved domains and TADs are critical for the primarily nuclear protein, Atos, to facilitate macrophage invasion.

**Figure 2.**
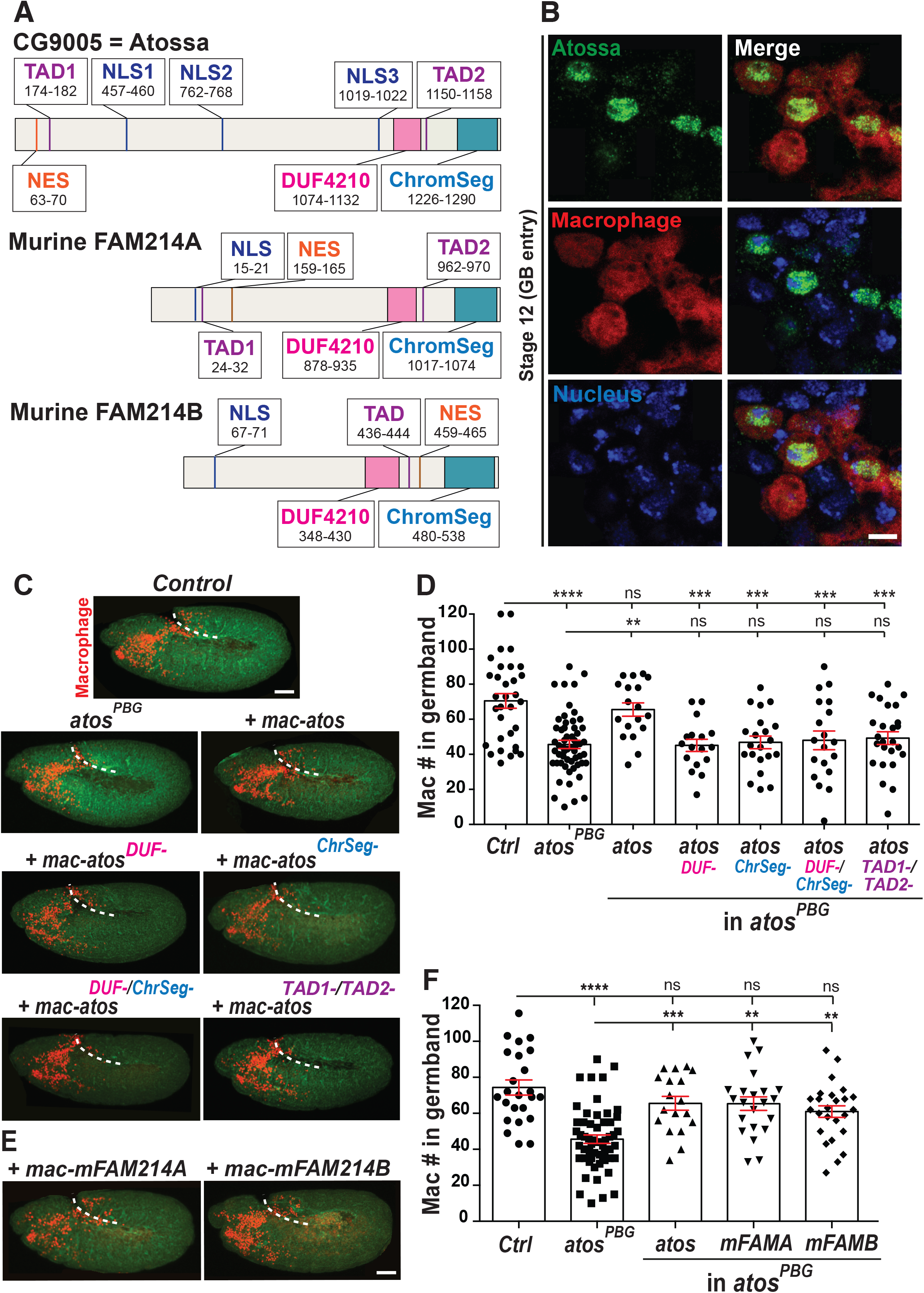
CG9005/Atossa requires conserved domains linked to transcriptional activation to enhance tissue invasion, a function maintained by its mammalian orthologs. **Fig 2A**. Deduced protein structure of *Drosophila* CG9005/Atossa (Atos) and its murine orthologs, mFAM214A and B. These proteins all contain the same conserved motifs: a domain of unknown function (DUF4210), a domain associated with Chromosome segregation (ChromSeg), at least one transcriptional activation domain (TAD), nuclear localization signal (NLS) and nuclear export signal (NES). FAM214A and B are 44-45% identical to Atos. **Fig 2B**. Macrophages (red) near the germband in Stage 11/12 embryos display colocalization of Atos tagged with HA (HA antibody, green) with the nucleus stained by DAPI (blue). *srpHemo-atos::H2A* line utilized. **Fig 2C**. Representative confocal images of Stage 12 embryos from the control, *atos*^*PBG*^, and *atos*^*PBG*^ expressing Atos itself or variants lacking particular domains in macrophages. **Fig 2D**. Germband macrophage quantification in control, *atos*^*PBG*^, and *atos*^*PBG*^ expressing Atos or its altered forms in macrophages. The tissue invasion defect in *atos*^*PBG*^ can be fully rescued by Atos expression in macrophages unless Atos lacks the conserved DUF4210, ChrSeg, or TADs. Control n=32, mutant n=56, WT rescue n=18, DUF4210^−^ rescue n=17, ChrSeg^-^ rescue n=21, DUF4210^−^/ChrSeg^-^ rescue n=19, TAD1^-^/TAD2^-^ rescue n=25. For control vs mutant p<0.0001, for control vs rescue p=0.99, for mutant vs rescue p=0.0014. **Fig 2E**. Representative confocal images of *atos*^*PBG*^ rescued with a murine ortholog, *mFAM214A* or *mFAM214B*, expressed in macrophages. **Fig 2F**. Quantification of macrophages in the germband in Stage 12 embryos from the control, *atos*^*PBG*^, and *atos*^*PBG*^ embryos expressing *mFAM214A* or *mFAM214B* in macrophages shows that Atos’s mammalian orthologs can rescue *atos*’s macrophage tissue invasion defect. Control n=25, *atos*^*PBG*^ n=56, rescue with *atos* n=18, with *mFAM214A* n=22, with *mFAM214B* n=25. For control vs *mFAM214A* and *mFAM214B* rescues p>0.05, for *atos*^*PBG*^ vs *mFAM214A* and *mFAM214B* rescues p<0.005. *mFAM214A* or *B* are expressed under the direct control of the *srpHemo* promoter. Throughout paper > indicates *GAL4 UAS* regulation. In (**C**) and (**E**) macrophages (red) are visualized by *srpHemo-H2A::3xmCherry* expression and actin by Phalloidin staining (green). One-way ANOVA with Tukey for (**D**) and (**F**). Scale bars are 5 µm in (**B**) and 50 µm in (**C**) and (**E**). See also **Figure S2**.

Atos’s uncharacterized murine orthologs, mFAM214A and mFAM214B, maintain these domains, displaying 40% identity to their *Drosophila* counterpart (Fig. 2A). Expression in macrophages of either mFAM214A or B in *atos*^*PBG*^ rescued the germband invasion defect as efficiently as the *Drosophila* protein itself (Figs. 2E-F) and restored the normal number of macrophages on the yolk next to the extended germband (Fig. S2E). Therefore, we conclude that the molecular functions that enable Atos to promote macrophage tissue invasion are maintained in vertebrates.

### Atos raises mRNA levels of an RNA helicase and metabolic enzymes, which are each required for germband invasion

Given Atos’s nuclear localization and requirement for TADs, we hypothesized that Atos might modulate transcription in macrophages to aid their initiation of germband invasion. To identify potential targets, we performed RNA-sequencing analysis on FACS isolated macrophages from wild type and *atos*^*PBG*^ embryos during germband invasion in early Stages 11-12 (Fig. S3A) (Supp. Data 1). Transcriptome analysis revealed 25 genes with reduced mRNA levels and 39 genes with higher ones in the absence of Atos, requiring a P value<0.05 (Fig. S3B). Gene ontology analysis (GO term) indicates that the significantly downregulated genes are involved in oxidation-reduction (redox) processes, stress responses as well as the nervous system (Fig. S3C). We therefore conclude that the presence of Atos in macrophages controls the mRNA levels of a small set of proteins.

We tested the hypothesis that the *atos*^*PBG*^ macrophage germband invasion defect is caused by the lower levels of the downregulated genes. We focused only on the 5 genes that had at least a >5-fold change in expression and were enriched in embryonic macrophages or had an identified molecular function (Fig. 3A). We expressed *RNAi* constructs against them in macrophages and observed a significant reduction in germband macrophage numbers for three of these 5 candidates (Figs. 3B-G, S3D-E). For all three we also observed an increase in the number of macrophages sitting on the yolk next to the germband before invasion, consistent with a specific defect in germband invasion (Figs. S3F-H). These were a predicted ATP-dependent RNA helicase (CG9253) we name Porthos (Pths) (Martin et al., 2021) (Figs. 3B, E), and two metabolic enzymes, <u>G</u>lyoxylate <u>R</u>eductase/<u>H</u>ydroxypyruvate <u>R</u>eductase (*dGR/HPR*, CG9331) (Figs. 3C and 3F) and <u>L</u>ysine α-<u>K</u>etoglutarate <u>R</u>eductase/<u>S</u>accharopine <u>D</u>e<u>h</u>ydrogenase (*dLKR/SDH*, CG7144) (Figs. 3D, G). Downregulation of *Glycerophosphate oxidase 2* (*Gpo2*, CG2137) (Fig. S3D) and *Golgi matrix protein 130 kD* (*GM130*, CG11061) (Fig. S3E) did not produce any invasion defect. GR/HPR is highly conserved from bacteria to mammals and the *Drosophila* form shows 48% identity to its human ortholog (NCBI BLAST). GR/HPR is the linchpin of the glyoxylate cycle, catalyzing the reduction of glyoxylate into glycolate and the conversion of hydroxypyruvate into D-glycerate (Fig. 3H) (Booth et al., 2006). This contributes to glucose and urea synthesis. The bifunctional enzyme dLKR/SDH is also highly conserved, with 71% identity to its human counterpart (identified by NCBI BLAST). It catalyzes the first two steps of lysine catabolism and can participate in the production of Acetyl CoA (Bhattacharjee, 1985) (Fig. 3I). We therefore conclude that Atos enhances macrophage tissue invasion by increasing the levels of the metabolic enzymes dLKR/SDH and dGR/HPR and the helicase ortholog Porthos.

**Figure 3.**
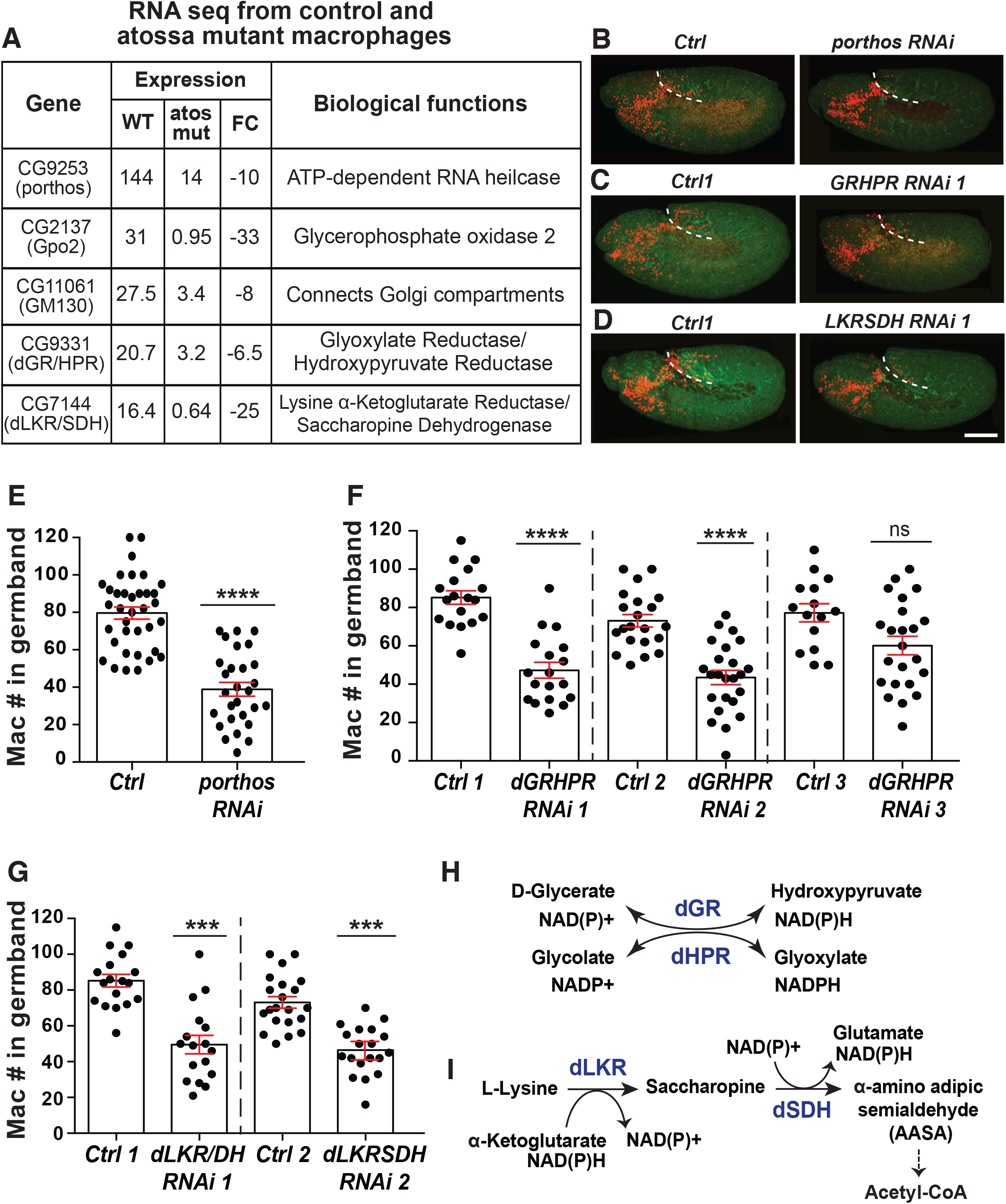
Atos leads to higher mRNA levels of an RNA helicase and metabolic enzymes required for germband invasion. **Fig 3A**. A selection of genes down-regulated in *atos*^*PBG*^ mutant macrophages compared to the control, chosen for having a >5 fold change in expression as well as an identified biological function. **Fig 3B-D**. Representative confocal images of early Stage 12 embryos from the control, and lines expressing an *RNAi* against (**B**) *porthos*, (**C**) *dGR/HPR* or (**D**) *dLKR/SDH* specifically in macrophages (red). *srpHemo-H2A::3XmCherry* labels macrophages. **Fig 3E**. Quantification of Stage 12 embryos reveals that expression of a *porthos RNAi* in macrophages decreases their number in the germband by 48%. Control n=36, *porthos RNAi* (BL36589) n=28, p<0.0001. **Fig 3F-G**. Quantification of Stage 12 embryos indicates that fewer macrophages have moved into the germband upon the expression in macrophages of any of (**F**) three different RNAis against *dGR/HPR* or (**G**) two different RNAis against *dLKR/SDH*, arguing that these metabolic enzymes are required in macrophages for tissue invasion. Control 1 n=18, *dGR/HPR RNAi 1* (VDRC 44653) n=18, p<0.0001, control 2 n=21, *dGR/HPR RNAi* 2 (VDRC 107680) n=24, p<0.0001, control 3 n=15, *dGR/HPR RNAi 3* (VDRC 64652) n=23, p=0.08. *dLKR/SDH RNAi 1* (VDRC 51346) n=17, control 2 n=21, *dLKR/SDH RNAi 2* (VDRC 109650) n=23, p<0.0001. **Fig 3H**. Schematic illustrates how the bifunctional enzyme dGR/HPR can catalyze the reduction of glyoxylate into glycolate and convert hydroxypyruvate into D-glycerate by oxidation of the cofactor NAD(P)H. **Fig 3I**. Schematic shows the metabolic pathway in which *Drosophila* Lysine α-Ketoglutarate Reductase/Saccharopine Dehydrogenase (dLKR/SDH) catalyzes the first two steps of the lysine catabolism pathway, resulting in the production of glutamate and acetyl-CoA, a TCA substrate, through several downstream enzymatic reactions. Glu: Glutamate, α-KG: α-Ketoglutarate, AASA: α-Aminoadipate δ-semialdehyde. Unpaired t test for (**E**-**G**). Scale bar: 50 µm in (**B**-**D**). See also **Figure S3** and **Data S1. Data S1:** Annotated primary and normalized RNA sequencing data from FACS sorted control and *atos* macrophages.

### The nuclear RNA helicase, Porthos, functions downstream of Atos in pioneer macrophages to allow their initiation of germband invasion

Atos’s target *porthos* (*CG9253*) displayed the strongest invasion defect upon RNAi knockdown (KD) (Fig 3E). Porthos is a conserved DEAD-box RNA helicase (Fig. S4A) sharing 71% identity and 84% similarity with its human ortholog, the helicase DDX47, including the conserved DEAD motif and helicase C terminal domain, with which DDX47 interacts with RNA structures. *porthos* is expressed in the embryo by *in situ* analysis in a pattern similar to *atos* but a few stages later, to *atos* in *Drosophila* embryos, being enriched in macrophages in the head region during Stages 9-12 (https://insitu.fruitfly.org/cgi-bin/ex/report.pl?ftype=1&ftext=FBgn0032919). In S2R+ cells, HA-tagged Porthos colocalized with markers for the nucleus (DAPI) and the nucleolus (Fibrillarin), where ribosome assembly and rRNA processing occur (Fig. 4SB). In embryonic macrophages HA-tagged Porthos also localized to the nucleus, detected by DAPI (Fig. 4A).

**Figure 4.**
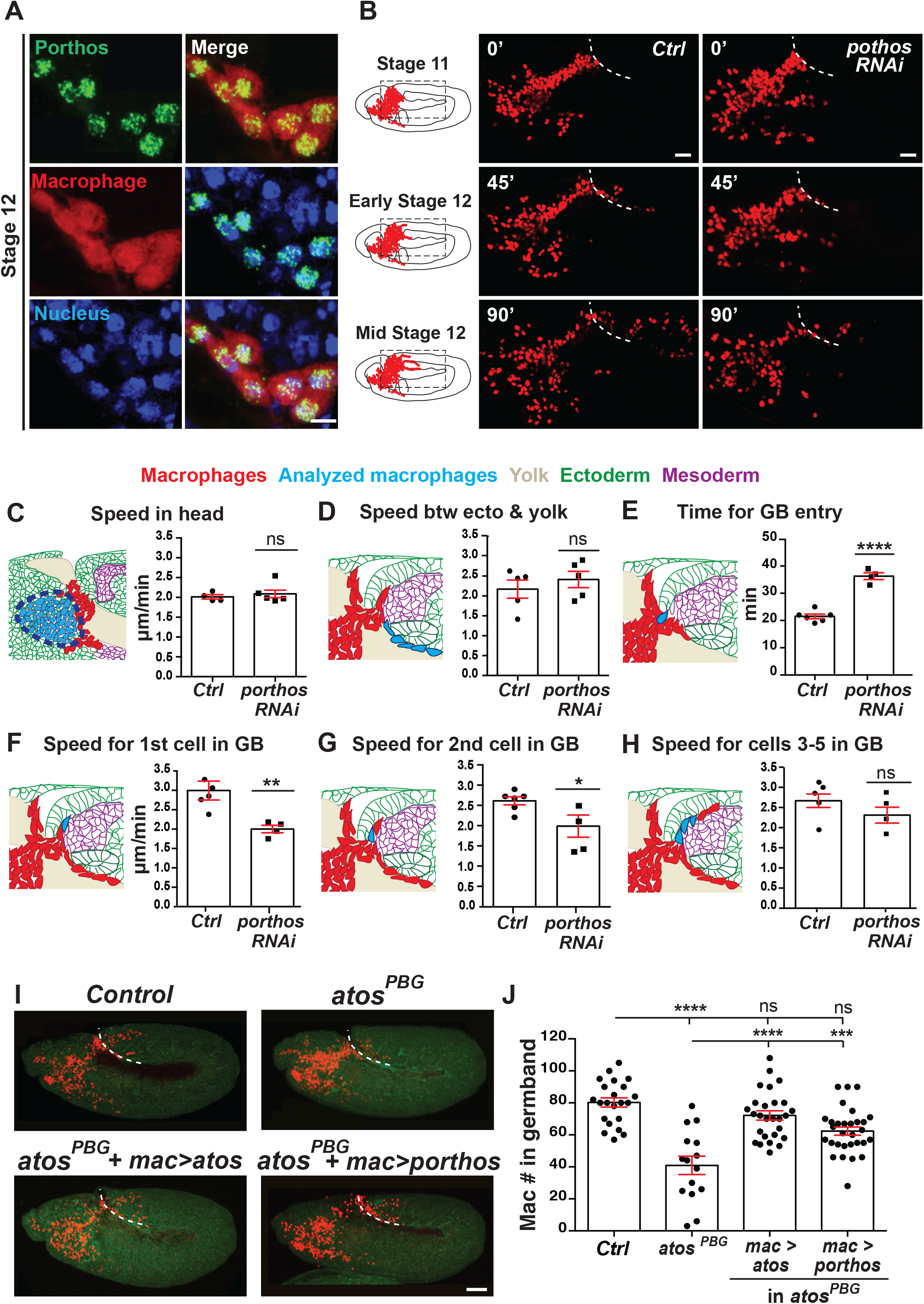
The nucleolar RNA helicase, Porthos, acts as a key downstream target of Atos to promote pioneer macrophage germband invasion. **Fig 4A**. Macrophages (red) near the germband in Stage 11/12 embryos show partial colocalization of the HA antibody labeling Porthos (green) with the nucleus stained by DAPI (blue). Embryo express *srpHemo-porthos::HA*. **Fig 4B**. Stills starting at Stage 11 from two-photon movies of control embryos and those expressing *porthos RNAi* in macrophages; stills show macrophage migration from the head mesoderm towards and into the germband at the indicated time points. White dotted line indicates the germband edge. Macrophage nuclei labeled by *srpHemo-H2A::3xmCherry. UAS-porthos RNAi* (BL36589) expressed by *srpHemo-GAL4*. **Fig 4C-H**. Quantification of macrophage migration parameters from two-photon movies. (**C-D)** Macrophages expressing *porthos RNAi* migrate with a similar speed in the head and between the yolk sac and the germband edge compared to the control. Speed in head: control=2.01 µm/min, *porthos RNAi*=2.09 µm/min; movie #: control=4, *porthos RNAi*=6; track #: control=507, *porthos RNAi*=859, p=0.56. Speed between yolk sac and germband mesoderm: control=2.17 µm/min, *porthos RNAi*=2.41 µm/min, p=0.45; movie #: control n=5, *porthos RNAi* n=5, track #: control n=40, *porthos RNAi* n=51. **Fig 4E**. The time required for the first macrophage nucleus to enter into the germband is significantly increased in embryos expressing *porthos RNAi* compared to the control. Control=21.5 min, n=6, *porthos RNAi*=36.2 min, n=4, p<0.0001. Blue arrow in schematic indicates route analyzed. **Fig 4F-G**. The speed of the first and second macrophage invading into the germband along the path between the mesoderm and ectoderm is significantly slower in embryos expressing *porthos RNAi* compared to the control. First macrophage speed: control=2.99 and *porthos RNAi*=2.0 µm/min; p=0.009; # movies: control n=4, *porthos RNAi* n=4. Second macrophage speed: control=2.61 and *porthos RNAi*=1.98 µm/min; p=0.037; # movies: control n=6, *porthos RNAi* n=4. **Fig 4H**. The speed of the third to fifth macrophages invading the germband is similar in macrophages downregulated for *porthos* and the control (speed: control=2.66 and *porthos RNAi*=2.31 µm/min; p=0.21; # movies: control n=5, *porthos RNAi* n=4). **Fig 4I**. Representative confocal images of early Stage 12 embryos from control, *atos*^*PBG*^, and *atos*^*PBG*^ expressing *atos::FLAG::HA* or *porthos::FLAG::HA* in macrophages (red) through *srpHemo-GAL4* control of *UAS* constructs. Embryo detected by phalloidin staining (green). **Fig 4J**. Quantification of macrophages in the germband shows that the *atos*^*PBG*^ mutant phenotype can be substantially rescued by expressing *porthos::FLAG::HA* in macrophages. Control (n=15), *atos*^*PBG*^ (n=22), *atos*^*PBG*^ with *srpHemo>atos::FLAG::HA* (n=27), *srpHemo>porthos::FLAG::HA* (n=30). For control vs *atos*^*PBG*^ p<0.0001, for control vs *atos* rescue of *atos*^*PBG*^ p<0.0001, for control vs *porthos* rescue of *atos*^*PBG*^ p=0.0007. Macrophages detected by cystoplasmic *srpHemo-3xmCherry* in (**A**) and nuclear *srpHemo-H2A::3xmCherry* in movies and in (**I**). Unpaired t test for (**C**-**H**), and one-way ANOVA with Tukey for (**J**). Scale bars: 50 µm in (**A**) and 30 µm in (**B**). See also **Figure S4** and **Videos 3** and **4**. Representative movies of macrophage migration into the germband in the control (**Video 3**) and the *porthos RNAi* embryos (**Video 4**). Macrophages (red) are labeled with *srpHemo-H2A::3xmCherry*. Arrow indicates first macrophage moving into the germband. The time interval between each acquisition is 40 s and the display rate is 15 frames/s. Scale bar: 20 μm.

To determine at which step in macrophage germband invasion Porthos is needed, we examined wild type embryos and those expressing *porthos* RNAi in macrophages. In fixed embryos we observed no change in migration along the non-invasive route of the vnc (Fig. S4C) or in the total number of macrophages compared to the control (Fig. S4D), arguing that Porthos is specifically required for migration into or within the tissues of the germband. We then utilized 2-photon imaging of live embryos and tracked macrophages as they moved from their initial position within the head towards the germband and then during their infiltration into this tissue (Videos 3-4, Fig. 4B and 4SE). We observed no significant change in speed or directionality in the head or on the yolk (Fig. 4C, Fig S4F-H) (speed: in head, 2 µm/min for control and *porthos RNAi*, p=0.56, and on yolk, control=2.1 and *porthos RNAi*=2.2 µm/min, p=0.35; directionality: in head, control=0.35 and *porthos* RNAi*=*0.37, p=0.27, and on yolk, control=0.42 and *porthos* RNAi=0.39, p=0.58). Moreover, we detected no significant change in the speed of macrophages moving on the yolk and beneath the germband beyond the entry point (control=2.2 and *porthos RNAi*=2.4 µm/min, p=0.45) (Fig. 4D). However, *porthos RNAi* macrophages waited 69% longer than the control to enter the germband tissue, (control=21.5 and *porthos RNAi*=36.3 min, p<0.0001) (Fig. 4E). Once within the germband, the first two macrophages invading between the mesoderm and ectoderm progressed significantly slower than the control (Fig. 4F-G) (1^st^ cell: control=3.0 and *porthos RNAi*=2.0 µm/min, p=0.009, 2^nd^ cell: control=2.6 and *porthos RNAi*=2.0 µm/min, p=0.037). In contrast, the speed of the subsequent macrophages was not significantly altered by *porthos RNAi* (Fig. 4H) (3^rd^-5^th^ cells: control=2.7 and *porthos RNAi*=2.3 µm/min p=0.21). Thus, *porthos RNAi* phenocopies *atos*’s migration defect. Finally, we expressed Porthos in *atos*^*PBG*^ to restore its higher levels in macrophages. This strongly improves the *atos* mutant phenotype (87% rescue) (Figs. 4I-J). Thus, we conclude that Porthos is a key player downstream of Atos, exerting an essential role in pioneer macrophages to specifically allow their initiation of germband invasion.

### Porthos alters translation of a subset of mRNAs

Given the helicase Porthos’s nucleolar localization we hypothesized that it might modulate translation. We purified ribosomes and polysomes by sucrose density gradient fractionation of the control and S2R+ cells treated with *porthos* dsRNA (Fig. 5A). We observed a reduction in polysomes, the 40S small subunit, and 80S ribosome fraction (Fig. 5B) along with an increase in the large 60S subunit peak in the *porthos* KD. This data suggests that Porthos is required for normal levels of 40S biogenesis, ribosome and polysome assembly, and supports the idea that the higher levels of Porthos triggered by Atos could affect mRNA translation.

**Figure 5.**
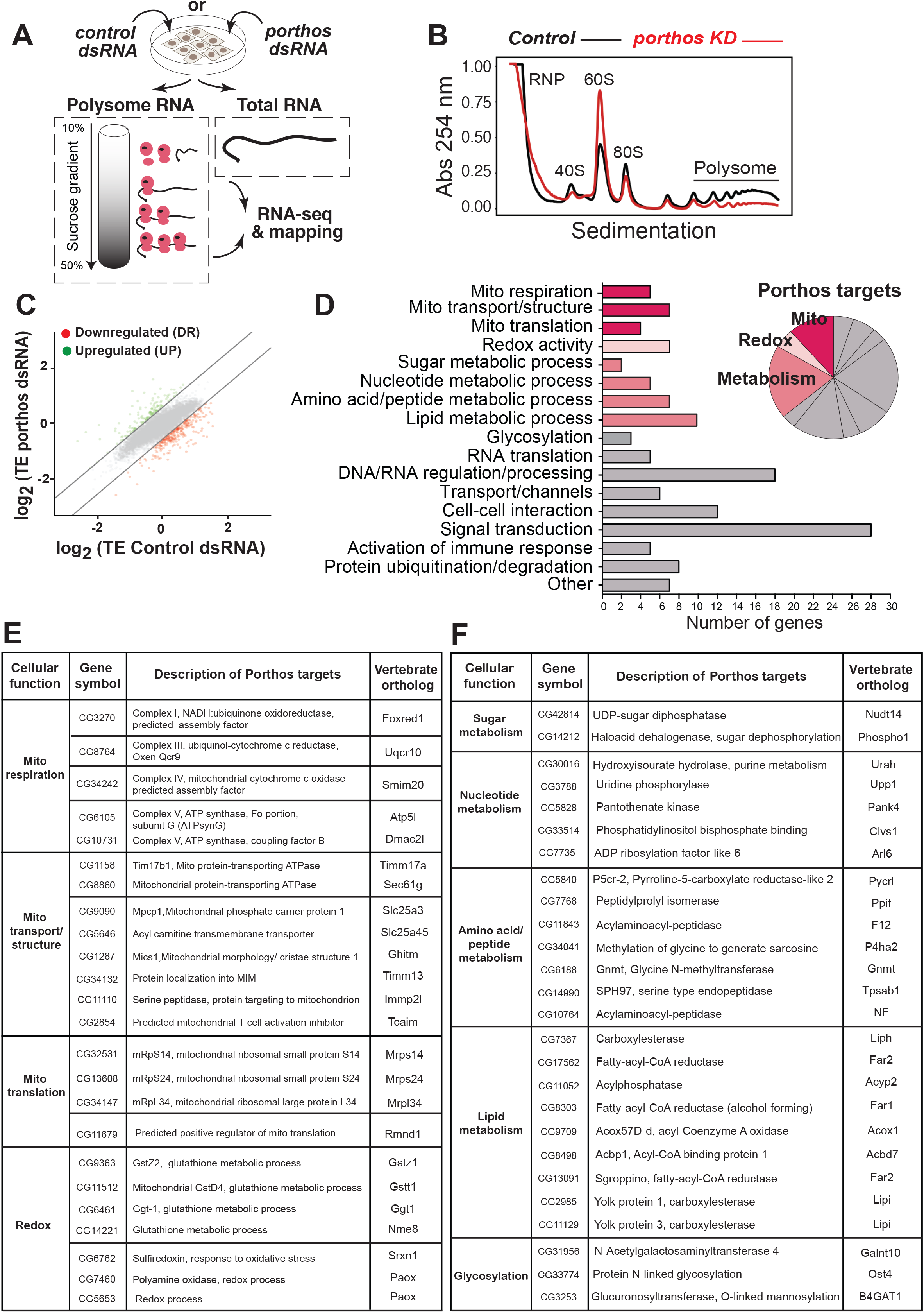
Porthos increases translation of an mRNA subset including many involved in mitochondrial OxPhos and metabolic processes. **Fig 5A**. Sucrose density gradient fractionation allowed purification of ribosome subunits and polysomes. Polysomal or total cellular mRNA fractions were isolated following dsRNA treatment and RNA sequencing libraries were prepared. **Fig 5B**. Sedimentation analysis showing the relative abundance of 40S, 60S, and 80S ribosomes indicates that *porthos* depletion by dsRNA markedly reduces the ratio of polysomes to monosomes. A non-targeting dsRNA was used as a control. Profiles were aligned on the basis of the 40S ribosome peak’s position and labeled with distinct colors, black for control and red for *porthos* KD, n=3 biological replicates. **Fig 5C**. Scatter plot of Translational efficiency (TE) values from *porthos dsRNA* S2R+ vs control *gfp dsRNA* cells. Red (down-regulated, DR) and green (up-regulated, UP) dots represent genes with log_2_ TE changes that meet the 2 standard deviation cutoff. **Fig 5D**. DR mRNAs in *porthos dsRNA* treated versus Control *dsRNA* treated S2R+ cells. 71% of the genes encoded proteins with predicted functions, the number corresponding to a functional category is shown. Proteins involved in mitochondrial-related functions, metabolic processes, and redox processes are highlighted. **Fig 5E**. Porthos modulates the translation of RNAs encoding components of mitochondrial OxPhos, including subunits of mitochondrial complexes III, and the ATP synthase complex V along with assembly factors for complexes I and IV. Porthos also enhances the TE of mitochondrial transporting channels, structural proteins as well as those involved in mitochondrial translation. **Fig 5F**. List of the proteins encoded by mRNAs that are downregulated in *porthos dsRNA* treated S2R+ cells that are involved in metabolic pathways. NF: Not Found. See also **Figure S5** and **Data S2. Data S2**: Complete set of TE values from polysome-sequencing data from control *gfp dsRNA* and *porthos dsRNA* treated S2R+cells.

To examine which mRNA transcripts depend on Porthos for their efficient translation, we performed polysome-profiling, sequencing transcripts associated with highly translationally active polysomes as well as all the transcripts in the S2R+ cells (Fig 5A, Supp. Data 2). We calculated translational efficiency (TE) as the ratio of the normalized reads present for each gene in the mRNAs from the polysome fraction to those in the total mRNA levels; this ratio was determined for the data from both the control *GFP RNAi* and *porthos KD* cells. We plotted the mean TE values for control (GFP KD) and *porthos* KD replicates and calculated the mean change in TE (ΔTE) for each gene as the ratio of TEs between control (GFP KD) and porthos KD replicates (Fig. 5C). Targets were defined as genes falling 2 standard deviations from the median ΔTE as previously described (Flora et al., 2018). We identified 204 annotated coding genes that were less efficiently translated and 102 that were more efficiently translated in *porthos KD* cells.

The mRNA targets whose TE Porthos enhances are involved in respiration, transport and translation in mitochondria, metabolic processes, transcription, translation, signal transduction, immune responses as well as redox processes (Fig. 5D-F, Fig. S5A-B). The targets include several components of mitochondrial OxPhos, namely ubiquinol cytochrome C reductase (complex III, UQCR-Q), ATP synthase subunit G and coupling factor F(o) (complex V), predicted assembly factors for complex I and IV, and proteins involved in mitochondrial translation and transport (Fig. 5E) as well as other metabolic pathways (Fig. 5F).

### ;Porthos is required for mitochondrial oxidative respiration and energy production

Mitochondria generate ATP through OxPhos mostly from the pyruvate formed by the glycolytic pathway (Pavlova and Thompson, 2016; Vander Heiden et al., 2009) (Fig. S6A) and thus can utilize metabolites downstream of the two enzymes we identified as Atos targets, LKR/SDH and GR/HPR. To directly investigate if Porthos regulates mitochondrial energy production, we generated S2R+ cells producing 56% of *porthos’s* normal mRNA levels with CRISPR/Cas9-mediated mutagenesis (which we call *porthos-KD* cells) (Fig. S6B). We then utilized a Seahorse assay in which the oxygen consumption rate (OCR) (Llufrio et al., 2018) is determined before and after sequential treatment with compounds affecting different steps in OxPhos (Figs. 6A, S6A). By comparing the OCR observed upon the different treatments we calculated OxPhos-dependent basal and maximum respiration and found that both were reduced 64% in *porthos-KD* (Figs. 6A-C) (see Methods). We also found significant decreases in OxPhos-dependent spare respiration capacity and as well as OxPhos-independent respiration (72% and 42% reduction, respectively) (Fig. 6C). S2R+ cells utilize primarily mitochondrial OxPhos rather than glycolysis for ATP production (Freijie et al., 2012); this remains the case even in the *porthos KD* cells (Fig. S6C) as we also observed a 60% reduction in the basal extracellular acidification rate (ECAR), a measure of lactate production through complete glycolysis (Fig. S6D). In totality, ATP production through OxPhos was reduced by 50% upon *porthos* depletion (Fig. 6C). Given that Porthos modulates the translation of subunits of mitochondrial complex III and the ATP synthase complex V, our data argues that Porthos induces a shift in metabolic capacity and flux that contributes to the upregulation of the OxPhos pathway and higher levels of energy production.

**Figure 6.**
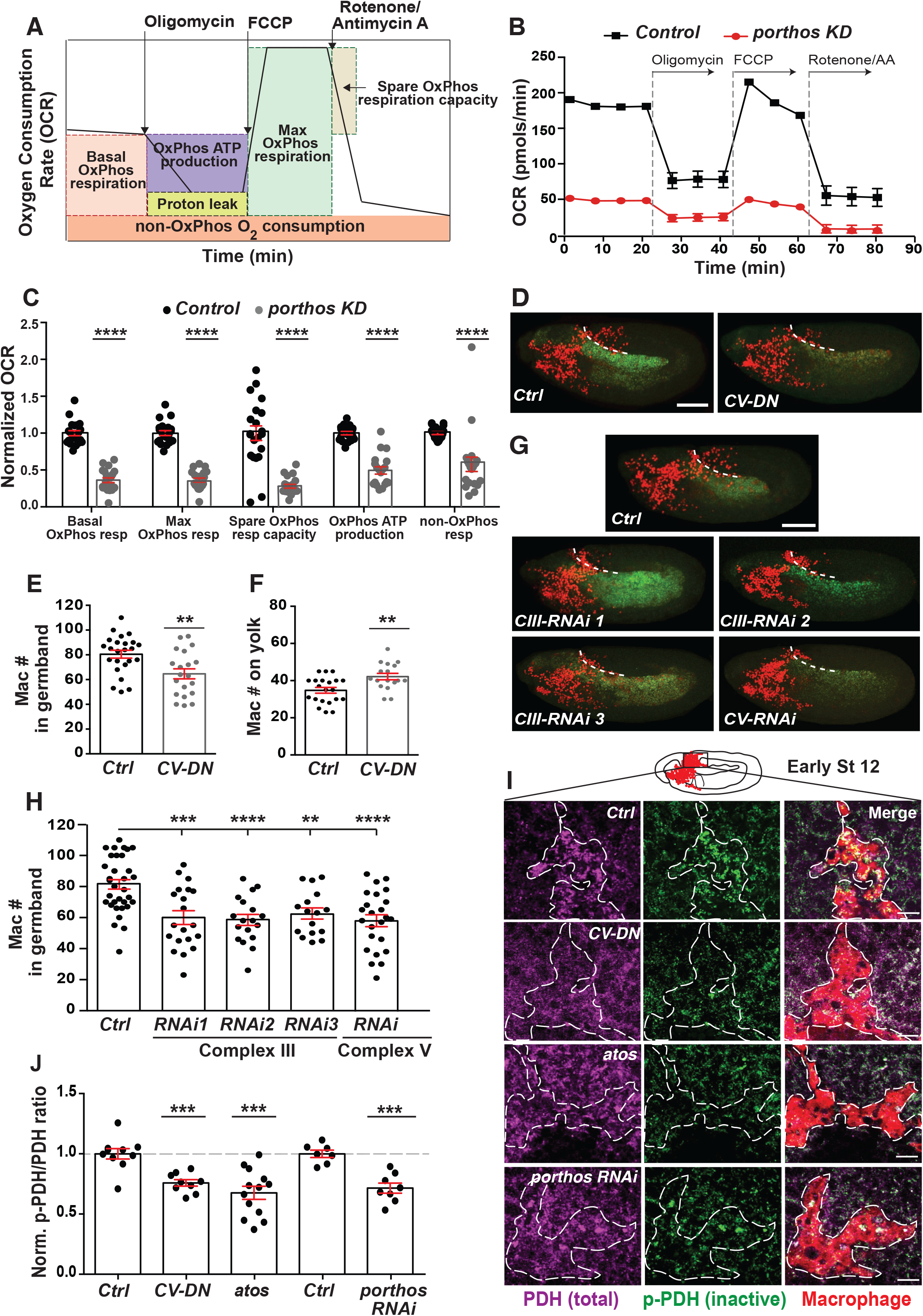
Higher levels of mitochondrial respiration are required in macrophages to power their germband tissue invasion. **Fig 6A**. Schematic of the procedure for mitochondrial energetic profiling in wild-type and *porthos KD* S2R+ cells with a Seahorse efflux assay. **Fig 6B**. The Oxygen Consumption Rate (OCR, pmols O_2_/min) was assessed as a representative parameter of OxPhos in control and *porthos KD* S2R+ cells by a Seahorse Bioscience XF96 Extracellular Flux Analyzer. The ATP synthase inhibitor oligomycin (2 µM), the uncoupler FCCP (2 µM), and the mitochondrial complex I inhibitor Rotenone (1 µM) with antimycin A (1 µM) were injected sequentially (see S6A). **Fig 6C**. Calculation from relative OCR values at different stages assesses basal and maximum OxPhos respiration, spare OxPhos respiration capacity, OxPhos ATP production, and non-OxPhos respiration rates. At least three independent biological experiments (n>6 technical replicates in each repeat). **Fig 6D-E. (D)** Representative confocal images and (**E**) quantification of Stage 12 embryos reveals that the number of macrophages (red) that penetrated into the germband in Stage 12 embryos is significantly decreased upon the expression of a dominant negative c-ring of the ATP synthase (CV-DN) compared to the control. Control n=24, *CV-DN* n=20, p=0.003. **Fig 6F**. Quantification of macrophages on the yolk in fixed early Stage 12 embryos shows a significant increase in the *CV-DN* embryos compared to the control. Control n=21, *CV-DN* n=17, p=0.003. **Fig 6G-H. (G)** Representative confocal images and (**H**) quantification of Stage 12 embryos indicates that fewer macrophages (red) move into the germband upon the expression in macrophages of any of three different *RNAis* against mitochondrial OxPhos *Complex III* (*Ubiquinol-cytochrome c reductase, UQCR*), or an *RNAi* against *Complex V* (*F1F0, CG3612*), arguing that these two components are required in macrophages for germband tissue invasion. Control n=34; *Complex III* (*Cyt-c1, CG4769*): *RNAi 1* (VDRC 109809) n=20, p=0.0001; *Complex III* (*UQCR-cp1, CG3731*): *RNAi 2* (VDRC 101350) n=18, p<0.0001; *Complex III* (*UQCR-cp2, CG4169*): *RNAi 3* (VDRC 100818) n=16, p=0.0027; *Complex V*: (*F1F0, CG3612*) *RNAi* (VDRC 34664) n=24, p<0.0001. **Fig 6I**. A single plane confocal microscope image during germband entry in early Stage 12 embryos from control (Ctrl) or *atos*^*PBG*^ embryos, or lines expressing *porthos* RNAi or *CV-DN* in macrophages. Antibodies used against the phosphorylated at S293 and thus inactivated Pyruvate Dehydrogenase (pPDH, green) or total PDH (magenta) in macrophages (red). Higher pPDH levels are usually found when ATP/ADP levels are high and input into the TCA cycle is being downregulated (Patel et al., 2014). **Fig 6J**. Quantification of normalized values for pPDH/PDH levels calculated from fluorescence intensities in macrophages from the genotypes in (**6I**) during initial germband invasion in early Stage 12. The pPDH/PDH ratio is significantly reduced in all compared to the control, arguing that decreasing the function of CV, *atos* or *porthos* in macrophages results in lower cellular ATP/ADP ratios compared to the control. Control n=10, *CV-DN* n=9, p=0.0002; *atos*^*PBG*^ n=13, level p=0.0002; control n=7, *macro>porthos RNAi* n=8, p=0.0002. Three independent experiments. Macrophages visualized in (**C**) and (**G**) with nuclear *srpHemo-H2A::3xmCherry* expression and (**I**) with cytoplasmic *srpHemo-3xmCherry*. Unpaired t test for (**B**-**C**), (**E**-**F**), and (**H**-**J**). Scale bars: 50 µm in (**D**) and (**G**), 10 µm in (**I**). See also **Figure S6**.

### Mitochondrial respiration is required for metabolism and energy production in macrophages to initiate invasion into the germband tissue

We sought to directly assess the importance of the OxPhos complexes whose TE is upregulated by Porthos for macrophage germband invasion in the embryo. Therefore, we tested the effect of a dominant negative form of *complex V*, the ATP synthase which converts the electron gradient produced during OxPhos into ATP (*CV-DN*) (Figs. 6D-F). We also expressed RNAis against different subunits of *complex III* and the α subunit of *complex V* in macrophages (Figs. 6G-H). Consistent with the polysome-profiling results from *porthos-KD* S2R+ cells, these treatments significantly reduced macrophage numbers within the germband (Figs. 6D-H) and increased them on the yolk at the germband entry site (Figs. 6F, S6E), phenocopying the germband invasion defect of *atos*^*PBG*^ or *porthos RNAi* in macrophages. We observed no significant difference in macrophage numbers on the vnc in late Stage 12 upon expression of *CV-DN* (Fig. S6F) or of the RNAis (Fig. S6G) compared to the control, indicating normal general migration. This data strongly supports the conclusion that higher levels of the OxPhos complexes III and V are required specifically for macrophage tissue invasion.

### Atos and its target Porthos increase macrophage bioenergetics for germband tissue invasion

To examine the bioenergetic state of embryonic macrophages *in vivo* in the absence of Porthos or Atos, we first assessed the Pyruvate dehydrogenase complex (PDH), which allows pyruvate formed by glycolysis to feed into the TCA cycle. PDH is a key point of metabolic regulation (Patel et al., 2014) (see Fig. 7A). Metabolites produced by the TCA cycle increase PDH’s phosphorylation thereby inhibiting it and thus the running of the cycle; metabolites utilized by the TCA cycle decrease PDH phosphorylation and activate it. Importantly, when energy levels fall and mitochondrial ADP levels rise, PDH is unphosphorylated and active, opening the gate to the TCA cycle and OxPhos (Patel et al., 2014). By antibody staining we determined the levels of phosphorylated inactive PDH (pPDH) and the total amounts of PDH (Lieber et al., 2019) in embryonic macrophages. We assessed the pPDH/PDH ratio; a smaller number indicates less inhibition and thus more activity of PDH. As a positive control we first examined macrophages expressing CV-DN, which blocks mitochondrial ATP synthase, and thus increases ADP levels. Indeed we observed a lower pPDH/PDH ratio than in the control (Fig. 6J). We also observed significantly lower pPDH/PDH ratios in macrophages invading the germband in *atos*^*PBG*^ embryos as well as those expressing *porthos* RNAi in macrophages compared to the control (Figs. 6I-J). Our results support the conclusion that in the absence of Atos or Porthos, macrophages *in vivo* have reduced ATP/ADP ratios, leading the cells to keep PDH in its active form to try to generate more ATP by converting pyruvate into acetyl CoA that can feed into the TCA cycle.

**Fig 7.**
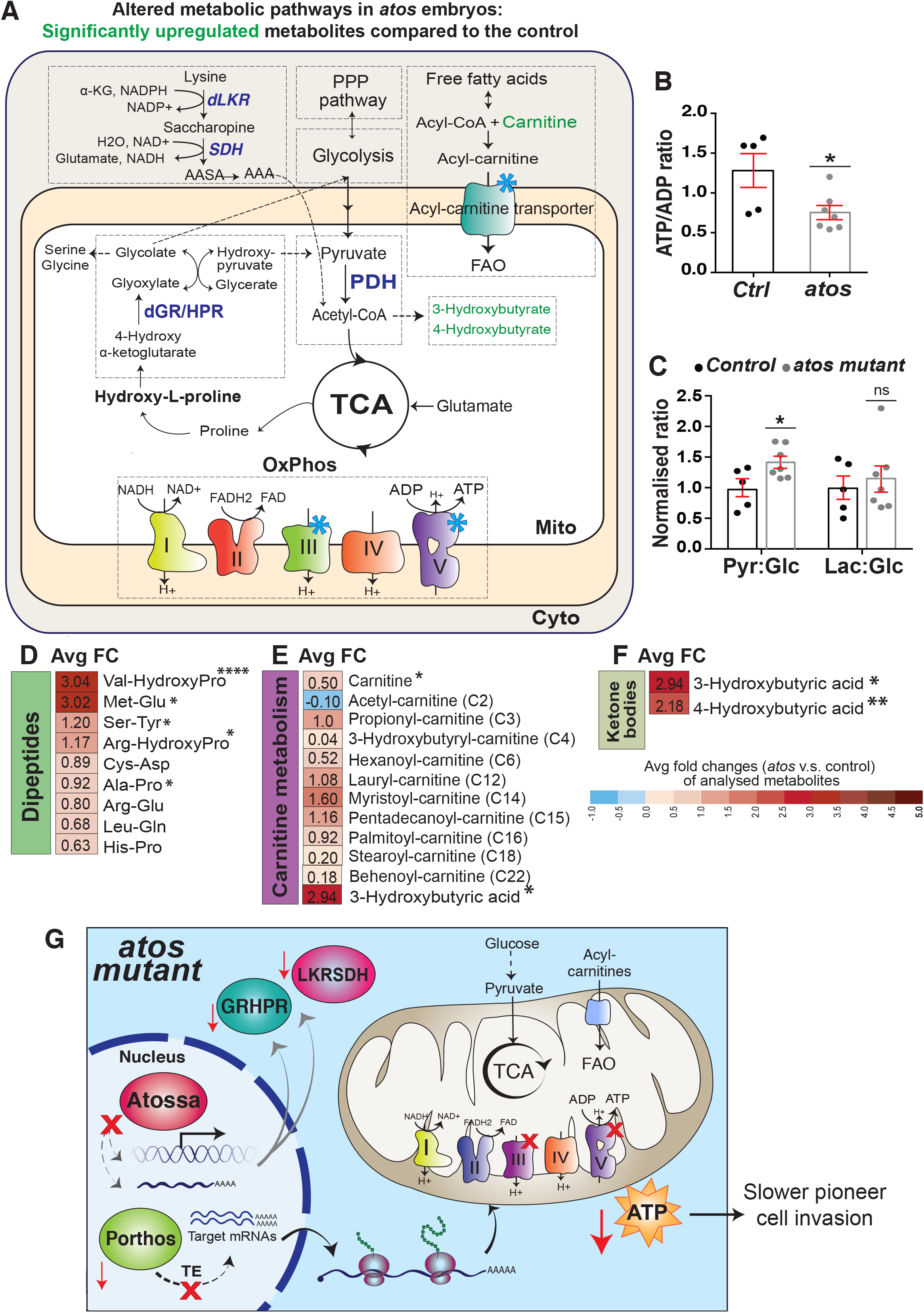
Mitochondrial metabolism is enhanced by Atos and Porthos. **Fig 7A**. Schematic depicting ATP-generating pathways in eukaryotic cells: glycolysis, the Pentose Phosphate Pathway (PPP), fatty acid oxidation (FAO), the TCA cycle, and the mitochondrial respiratory electron transport chain (ETC). Blue stars mark *porthos* targets. Green indicates individual metabolites with statistically significant upregulation in *atos*^*PBG*^ compared to the control. **Fig 7B-F**. Cellular metabolites were measured by LC–MS-based metabolomics from extracts of Stage 11 embryos (Control n=5, *atos*^*PBG*^ n=7). **Fig 7B**. Normalized ATP/ADP ratio values are decreased in *atos*^*PBG*^ compared to control embryos. (p-value=0.028). **Fig 7C**. Quantification in *atos*^*PBG*^ compared to wild-type embryos shows an increase in the pyruvate/glucose ratio (p-value=0.035), but none for the Lactate/Glucose ratio (p-value=0.65). **Fig 7D-F**. Heatmap of non-targeted metabolites in *atos*^*PBG*^ compared to wild-type embryos shown with average log_2_fold change (FC) reveals **(D)** a significant increase in some dipeptides including those containing hydroxyproline, **(E)** increases in intermediates of mitochondrial fatty acid β-oxidation (FAO), including different carnitine-conjugated lipids, and **(E-F)** a significant increase in ketone body substituents compared to the control. *p<0.05, **p<0.01, ***p<0.001, ****p<0.0001. Values are obtained from untargeted metabolomic analysis in (**B, D-F**). **Fig 7G**. Our model: Atos raises mRNA transcript levels of the helicase Porthos and the metabolic enzymes GR/HPR and LKR/SDH in macrophages. Metabolic pathways downstream of GR/HPR and LKR/SDH are known to produce metabolites that feed into glycolysis and the TCA cycle to produce ATP. Porthos enhances the translational efficiency of mRNAs, including those encoding mitochondrial OxPhos components and a mitochondrial carnitine transporter. Macrophages with elevated mitochondrial OxPhos can meet the demands for the energy needed to create a path for tissue invasion. However, the absence of Atos leads to reduced levels of GR/HPR, LKR/SDH and Porthos. This decreases generation of ATP through OxPhos leading to defective tissue infiltration of the pioneering macrophages. Unpaired t-test for (**B**-**F**). See also **Figure S7** and **Data S3. Data S3:** Primary metabolomics data from control and *atos* mutant embryos.

### Atos enhances cellular metabolism and ATP/ADP levels

To investigate the full complement of metabolic changes that Atos enables, we performed untargeted comparative metabolite profiling by capillary liquid chromatography-tandem mass spectrometry (LC-MS/MS) (Figs. S7A, 7A) characterizing extracts from control and *atos*^*PBG*^ embryos. Most importantly, consistent with the results we had observed in the Seahorse assay and the p-PDH/PDH ratio measurement, we observed a significantly decreased ATP/ADP ratio in the absence of Atos (Fig. 7B). Thus our metabolic data supports that Atos regulates a set of targets that shift metabolism to enhance ATP production. Consistent with Atos’s role in increasing GR/HPR levels, in *atos*^*PBG*^ we observed higher levels of this enzyme’s substrate, 4-hydroxy α-ketoglutarate (4-HαKG) (Figs. S7B-C) and of hydroxy-L-proline (HLP), the metabolite just upstream of 4-HαKG (Fig. 7A). We also observed significantly higher levels of dipeptides containing HLP (Fig. 7D). Atos also regulates LKR/SDH; we observed a reduction to 60% of control levels of its product alpha-amino adipic semialdehyde (ASAA), by targeted-metabolomics profiling (Fig. 7A).

As the metabolomics was conducted on embryos constitutively defective in Atos, we expected to see some compensatory changes as well. Matching our previous data, no indications of a metabolic shift away from mitochondrial OxPhos towards aerobic glycolysis in the absence of Atos were present (Figs. 7C, S7D-F). Instead we observed results consistent with a backup of some metabolites whose products would normally be fed into glycolysis and the TCA cycle. We found significantly higher levels of β-hydroxybutyrate, which can be broken down to acetyl-CoA (Puchalska and Crawford, 2017) along with increases in carnitine-conjugated fatty acids (Figs. 7E-F). There were strong increases in thymidine, which can be catabolized to a product that is fed into glycolysis (Tabata et al., 2017), and uridine which can be interconverted with thymidine, along with other purine and pyrimidine nucleotides (Figs. S7G-H). We observed a small increase in most amino acids in *atos*^*PBG*^ (Fig. S7I). Additionally strong reductions occurred in the glycine-related metabolite sarcosine (N-methylglycine) known to be a biomarker of highly metastatic prostate cancer (Fig. S7J) (Sreekumar et al., 2009; Zhang et al., 2012).

In sum, the metabolomics profiling data in combination with our other findings strongly supports the conclusion that Atos is a powerful regulatory protein, increasing the efficiency and amount of OxPhos by inducing a metabolic shift that affects the ETC and complex V as well as the TCA cycle (Fig. 7G).

## DISCUSSION

We identify a key regulator of energy levels in *Drosophila* macrophages as a highly conserved and previously uncharacterized nuclear protein, that we name Atos. Atos mRNA is deposited maternally and is ubiquitously expressed at low levels. However Atos mRNA is also developmentally upregulated in macrophages several hours prior to tissue invasion and down regulated after invasion is completed. Live imaging shows that the presence of Atos speeds the tissue entry and forward movement within the germband tissue of only the first two macrophages, the invasion pioneers. RNA sequencing indicates that Atos leads to the upregulation in macrophages of mRNAs encoding two metabolic enzymes, dGR/HPR by 6.5-fold and dLKR/SDH by 25-fold, as well as a 10-fold increase in the mRNA encoding an ATP-dependent RNA helicase, named Porthos. Each of these three proteins is required for normal amounts of invasion. We show in S2R+ cells that two-fold higher levels of Porthos mRNA correspond to two-fold higher OxPhos activity, a process that generates ATP by transferring electrons from NADH and FADH2 produced by the TCA cycle to oxygen (Martínez-Reyes and Chandel, 2020). We thus favor the hypothesis that these two metabolic enzymes act in pathways that ultimately feed into the TCA cycle and thus the ETC. We identify an increase in the active state of the PDH enzyme in *porthos* and *atos* mutant macrophages, an indirect indication that ATP could be lower without these proteins. Importantly, we directly detect two-fold lower ATP/ADP levels in *atos* mutant embryos. Given that Atos is much more highly expressed in macrophages at this stage than in the rest of the embryo, the effects within these immune cells will be even greater. In sum our data argues that the developmentally programmed upregulation of Atos triggers a metabolic shift by upregulating this triad of targets, ultimately significantly increasing ATP/ADP in all macrophages and thereby enabling pioneer macrophages to power the creation of a path for tissue infiltration against surrounding resistance. Our findings are consistent with previous work indicating that higher ATP levels are needed in the first cell to migrate through extracellular matrix (Kelley et al., 2019; Zhang et al., 2019). However, to our knowledge our work is the first to identify a concerted molecular pathway that can produce the higher energy levels needed to speed pioneer cell invasion.

The target of Atos that we have focused on in this study is a previously uncharacterized protein we call Porthos. Porthos belongs to a family of ATP-dependent DEAD-box RNA helicases that have essential roles in RNA metabolism (Martin et al., 2021, Bourgeois et al., 2016; Venema et al., 1997; Venema and Tollervey, 1995). We find Porthos localized to the nucleolus in macrophages, suggesting a function in ribosome production or assembly (Baßler and Hurt, 2019). Porthos’ vertebrate ortholog, DDX47, binds rRNA precursors (Sekiguchi et al., 2006); its *S. cerevisiae* ortholog, RRP3, can separate short RNA helices and is required for the RNA processing that produces the 18S rRNA component of the 40S ribosomal subunit (O’Day et al., 1996; Garcia et al., 2012). Consistent with this in S2R+ cells treated with *porthos* dsRNA we find a lower ratio of the 40S to the 60S ribosome subunits along with a strong decrease in multiple ribosomes sitting on an mRNA, called polysomes. Importantly, Porthos also enhances the translational efficiency (TE) of a subset of mRNAs. A significant subset of the mRNAs whose TE is enhanced by Porthos encode mitochondrial proteins. These are orthologs of proteins shown to affect many aspects of the organelle’s biology, from its specialized translation, its import of proteins and their insertion into the inner membrane where the ETC resides, to its import of fatty acids as fuel. Some of these targets are also components directly involved in OxPhos. We identify two orthologs of proteins that affect the assembly and function of OxPhos complexes I and IV (Formosa et al., 2015; Dennerlein et al., 2015), one of which causes mitochondrial disease if mutated (Calvo et al., 2010). The yeast ortholog of the complex III subunit we identify as a target, QCR9, is required to strongly increase reductase activity (Brandt et al., 2017). We identify a protein whose ortholog has been implicated in ATP synthase function (Belogrudov, 2002; Belogrudov, 2008). Another, complex V subunit G, fosters complex dimerization, thereby contributing to cristae formation (Davies et al., 2011; Hahn et al., 2016), as does another target, Mics1 (Oka et al., 2008). More cristae correlate with higher levels of OxPhos (Brandt et al., 2017), and have been proposed to foster ATP production (Mannella, 2020). Thus the increased OxPhos we see in cells with more Porthos could result from improved efficiency through multiple avenues; more translation of Porthos targets would be predicted to increase the amount, localization, and assembly of OxPhos components as well as the extent of the membrane folds in which they are localized. Co-regulation to increase this set of mitochondrial proteins could thus allow a concerted enhancement of OxPhos and mitochondrial energy production by avoiding that invidual steps become rate limiting.

Atos’s two mammalian orthologs, FAM214A and B, can fully substitute for Atos during macrophage invasion, arguing that they maintain Atos’s ability to increase ATP/ADP. All of Atos’s targets that we show act during invasion have highly conserved human orthologs whose mRNAs are broadly expressed along with FAM214A and B (Sekiguchi et al., 2006; Human Protein Atlas, BioGPS). Thus Atossa’s vertebrate orthologs could be utilized by particular mammalian cell types in energetically demanding circumstances. In the immune system FAM214A appears particularly enriched within plasmacytoid dendritic cells (pDCs) and B cells (Table 1); pDCs upregulate OxPhos in response to IFN-1s during anti-viral responses (Wu et al., 2016) and B cells upregulate OxPhos during differentiation for effective antibody secretion (Price et al., 2018). Furthermore, FAM214A and B are well expressed in the brain which utilizes large amounts of energy and produces almost all of its ATP though OxPhos (Raichle and Gusnard, 2002; Hall et al., 2012). A shift from aerobic glycolysis to OxPhos is required for neural stem cell differentiation and neural survival (Zheng et al., 2016); many neurodegenerative diseases are associated with defects in OxPhos (Koopman et al., 2013). Interestingly, four different single nucleotide polymorphisms (SNP) in FAM214A introns have been linked to more severe Alzheimer’s disease or neurofibrillary tangles in genome wide association studies while another SNP in a transcription factor-binding region was associated with increased general intelligence (p-value for all variants≤5×10^−6^; https://www.ebi.ac.uk/gwas/search?query=FAM214A), Sherva et al., 2020; Beecham et al., 2014; Wang et al., 2020; Davies et al., 2018). The importance of OxPhos enhancers for brain function is demonstrated by the many neurodegenerative diseases connected to defects in PGC-1 (Zheng et al., 2010; Cui et al., 2006; Weydt et al., 2006). PGC-1 activates OxPhos through transcription of mitochondrial genes and thus mitochondrial biogenesis (Lin et al, 2005). In contrast, Atossa increases translation from mitochondrial mRNAs that already exist by raising levels of the helicase Porthos. The closest *Drosophila* ortholog of PGC-1 also raises transcription of mitochondrial proteins and OxPhos (Tiefenbock et al., 2010) and is expressed in invading macrophages at levels comparable to Atossa in our RNAseq. We hypothesize that these two regulators of mitochondrial function could work in concert, with Atossa allowing faster and more easily reversible control of enhanced energy production. Thus investigating the mammalian version of the regulatory network that we identified in this work and strategies to modulate it in the brain and immune system is of wide interest.

Altogether, our work uncovers a surprising molecular genetic view into the physiology of the organism, revealing a heretofore unsuspected cross-regulatory mechanism that spans different levels of the biological organization of cellular metabolism, cell biology and the tissue invasiveness of the immune system.

## Supporting information

Key resource table

## Acknowledgements

We thank: The *Drosophila* Genomics Resource Center supported by NIH grant 2P40OD010949-10A1 for plasmids, the Bloomington *Drosophila* Stock Center supported by NIH grant P40OD018537 and the Vienna *Drosophila* Resource Center (Dietzl et al., 2007) for fly stocks, FlyBase (Thurmond et al., 2019) for essential genomic information, and the BDGP *in situ* database for data (Tomancak et al., 2002; Tomancak et al., 2007). We thank the Vienna BioCenter Core Facilities for RNA sequencing and analysis and the Life Scientific Service Units at IST Austria for technical support and assistance with microscopy and FACS analysis. We thank C. Guet, C. Navarro and Siekhaus group members for discussions and comments on the manuscript. D.E.S. was funded by Marie Curie CIG 334077/IRTIM and Austrian Science Fund (FWF) grant ASI_FWF01_P29638S, P.R. by NIH/NIGMS (R01GM111779-06 and RO1GM135628-01), and A.B. with support of the European Research Council (ERC) under the European Union’s Horizon 2020 research and innovation program, grant no. 677006, ‘‘CMIL’’. T.R.H. is supported by the Natural Sciences and Engineering Research Council of Canada (RGPIN-2019-06766).

## Author Contributions

S.E., E.T.M. A.G. J.B. J-W.G. and T.K. conducted experiments, T.R.H. and A.B. provided resources, S.E., T.R.H., J.B. J-W. A.G, T.K. P.R. and D.E.S. designed experiments, S.E., E.T.M., and D.E.S. wrote the paper, S.E., E.T.M. and J-W.G. conducted formal analysis, all authors reviewed and edited the paper, A.B., D.E.S. and P.R. conducted Supervision and Project Administration, S.E., D.E.S. and P.R carried out Conceptualization. D.E.S., P.R. A.B. and T.R.H. acquired funding.

## Declaration of Interests

The authors declare no competing interests.

## EXPERIMENTAL PROCEDURES

### Fly work

Flies were raised on food bought from IMBA (Vienna, Austria) which was prepared according to the standard recipe of agar, cornmeal, and molasses with the addition of 1.5% Nipagin. Adults were placed in cages in a Percival DR36VL incubator maintained at 29°C and 65% humidity or a Sanyo MIR-153 incubator at 29°C within the humidity controlled 25°C fly room; embryos were collected on standard plates prepared in house from apple juice, sugar, agar and Nipagin supplemented with yeast from Lesaffre (Marcq, France) on the plate surface. Embryo collections for fixation (7-8 hour collection) as well as live imaging (4-5 hour collection) were conducted at 29°C.

### Fly lines obtained used in this work

*srpHemo-GAL4* was provided by K. Brückner (Brückner et al., 2004). The RNA lines tested in this paper (Table S1) were obtained from the Bloomington *Drosophila* Stock Centre (Bloomington, USA) and the Vienna *Drosophila* Resource Center (VDRC, Vienna, Austria). Lines *w*^*-*^; *P{w[+mC] srpHemo-3xmCherry}, w*^*-*^; *P{w[+mC] srpHemo-H2A::3xmCherry}* were published previously (Gyoergy et al., 2018).

### Embryo fixation and immunohistochemistry

Embryos were collected on apple juice plates from between 6-8.5 hours at 29°C. Embryos were incubated in 50% Chlorox (DanClorix) for 5 min and washed. Embryos were fixed with 17% formaldehyde/heptane (ThermoFisher Scientific, Waltham, MA, USA) for 20 min followed by methanol or ethanol devitellinization. PDH and p-PDH staining utilized hand-devitellinized embryos. Fixed embryos were blocked in BBT (0.1 M PBS + 0.1% TritonX-100 + 0.1% BSA) for 2 hours at RT and then incubated overnight at 4°C. Antibodies were used at the following dilutions: Mouse anti α-GFP (Aves Labs Inc., Tigard, Oregon, 1:500), Rat anti-HA (Roche, Basel, Switzerland, 1:100), Mouse anti-PDH E1α (Abcam, Cambridge, UK, ab110334, 1:200) and Rabbit antiphoshpo-PDH E1α (S293) (Abcam, Cambridge, UK, ab92696, 1:200). Afterwards, embryos were washed in BBT for 2 hours, and incubated with secondary antibodies at RT for 2 hours, and washed again for 2 hours. Secondary antibodies and Phalloidin were used at the following dilutions: anti-rat 488 1:300, anti-chicken 488 1:500, anti-mouse 488 1:500 or anti-mouse 633 1:200, anti-rabbit 488 1:300, and Phalloidin 1:300 (all from ThermoFisher Scientific, Waltham, MA, USA) (Table S2). The embryos were mounted overnight at 4°C in Vectashield mounting medium (Vector Laboratories, Burlingame, USA), which contains DAPI. Embryos were placed on a slide and imaged with a Zeiss Inverted LSM800 Confocal Microscope using a Plain-Apochromat 20X/0.8 Air Objective or a Plain-Apochromat 63X/1.4 Oil Objective.

### S2R+ cell work and immunostaining

S2R+ cells (a gift from Frederico Mauri of the Knöblich laboratory at IMBA, Vienna) were grown in Schneider’s medium (Gibco) supplemented with 10% FBS (Gibco) and transfected with the *srpHemo-HA::CG9005*(*atos*), or *UAS-CG9005(atos)::FLAG::HA, UAS-<u>CG9253</u>*(*<u>porthos</u>*)*<u>::FLAG::HA</u>* and *srpHemo-GAL4* constructs using Effectene Tranfection Reagent (Qiagen, Hilden, Germany) following the manufacturer’s protocol (Table S3). Transfected S2R+ cells were grown on Poly-L-Lysine coated coverslips (ThermoFisher Scientific, Waltham, Massachusetts, USA) in complete Schneider’s medium (Gibco) supplemented with 10% FBS (Sigma-Aldrich, Saint Louis, Missouri, USA) to a confluency of 60%. For antibody staining, cells were fixed with 4% paraformaldehyde (Sigma-Aldrich, St Louis, MI, USA) in PBS for 15 minutes at room temperature (RT). Cells were washed three times with PBS followed by permeabilization with 0.5% Triton X-100 (Sigma-Aldrich) in PBS for 15 minutes and then blocked in BBT (see above) for at least 1 hour. Antibodies were diluted in blocking buffer and incubated for 2 hours at RT. Primary antibodies were used at the following working dilutions: Chicken anti-GFP (clone 5G4, Ogris lab, MFPL, 1:100), Rat anti-HA (Roche, Basel, Switzerland, 1:50), Mouse anti-Lamin (DSHB, lamin Dm0, ADL1010, 1:50), and Mouse anti-fibrillarin (gift from Rangan lab, 1:1). Cells were subsequently washed three times with PBS-Triton X-100 (0.05%) for 5 minutes each, followed by secondary antibody incubation in blocking/permeabilization buffer for 1 hour at RT. Secondary antibodies were used at the following working dilutions: anti-rat Alexa Flour 488 (1:50), anti-mouse Alexa Flour 488 (1:200), and anti-mouse Alexa Flour 633 (1:100) (all from ThermoFisher Scientific, Waltham, MA, USA). Cells were counterstained with DAPI (ThermoFisher Scientific) for 10 minutes in PBS-Triton X-100 (0.05%). After immunoblotting, cells were mounted with Vectashield (Sigma-Aldrich). Images were acquired using the Zeiss inverted LSM-800 confocal microscope. Pictures were processed with ImageJ.

### DNA isolation from single flies

Single male flies were frozen overnight before being grounded with a pellet homogenizer (VWR, Radnor, USA) and plastic pestles (VWR, Radnor, USA) in 50µl of homogenizing buffer (100 mM Tris-HCl, 100 mM EDTA, 100 mM NaCL, and 0.5% SDS). Lysates were incubated at 65°C for 30 minutes. Then 5 M KAc and 6 M LiCl were added at a ratio of 1:2.5 and lysates were incubated on ice for 10 min. Lysates were centrifuged for 15 minutes at 20,000xg, supernatant was isolated and mixed with Isopropanol. Lysates were centrifuged again for 15 minutes at 20,000xg, the supernatant was discarded and the DNA pellet was washed in 70% ethanol and subsequently dissolved in distilled water.

### FACS sorting of macrophages

For embryo collections, adult flies of either *w*^*+*^; *srpHemo-3xmCherry* or *w*^*+*^; *CG9005*^*BG02278*^; *srpHemo-3xmCherry* genotypes were placed into plastic cages topped with apple juice plates with yeast for egg laying. Collections were performed at 29°C at 8h-20h light-dark cycle. Macrophages were collected from Stage 11-early Stage 12, when macrophages initiate invasive migration into the extended germband. Briefly, adult flies laid eggs for 1 hour, then the isolated plates with embryos were kept at 29°C for an additional 4 hours 45 minutes to reach the desired age. Embryos were collected for 2 days with about 6-7 collections per day and stored meanwhile at +4°C to slow down development. Collected embryos were dissociated and the macrophages were sorted according to the procedure described in (Gyoergy et al., 2018). The cells were sorted using a FACS Aria III (BD) flow cytometer. Emission filters were 600LP, 610/20 and 502 LP, 510/50. Data was analyzed with FloJo software (Tree Star). The cells from the negative control embryos were sorted to set a baseline plotAbout. Approximately 1-1.5×10^5^ macrophages were sorted within 30 minutes.

### Sequencing of the macrophage transcriptome

Total RNA was isolated from the FACS-sorted macrophages using the Qiagen RNeasy Mini kit (Cat No. 74104). The quality and concentration of RNA was determined using the Agilent 6000 Pico kit (Cat No. 5067-1513) on the Agilent 2100 Bioanalyzer: about 100 ng of total RNA was extracted from 1.5 × 10^5^ macrophages. RNA sequencing was performed by the CSF facility of the Vienna Biocenter according to their standard procedures (https://www.vbcf.ac.at/facilities/next-generation-sequencing/). Briefly, a cDNA library was synthesized using the QuantSeq 3’ mRNA-seq Library Prep kit and 4 replicates of each of the genotypes (*w+; +; srpHemo::3xmCherry* or *w*^*+*^; *CG9005*^*BG02278*^; *srpHemo-3xmCherry)* were sequenced on the Illumina HiSeq 2500 platform.

The reads were mapped to the *Drosophila melanogaster* Ensembl BDGP6 reference genome with STAR (version 2.5.1b). The read counts for each gene were detected using HTSeq (version 0.5.4p3). The Flybase annotation (r6.19) was used in both mapping and read counting. The counts were normalised using the TMM normalization from the edgeR package in R (Anders and Huber, 2015; Dobin et al., 2013). (Prior to statistical testing the data was transformed and then the differential expression between the sample groups was calculated with the limma package in R. The functional analyses were done using the topGO and gage packages in R.

### Time-lapse imaging

Embryos were dechorionated in 50% bleach for 4 min, washed with water, and mounted in halocarbon oil 27 (Sigma) between a coverslip and an oxygen permeable membrane (YSI). The anterior dorsolateral region of the embryo was imaged on an inverted multiphoton microscope (TrimScope, LaVision) equipped with a W Plan-Apochromat 40X/1.4 oil immersion objective (Olympus). mCherry was imaged at an 820 nm excitation wavelength, using an optical parametric oscillator technology (Coherent Chameleon Compact OPO). Excitation intensity profiles were adjusted to tissue penetration depth and Z-sectioning for imaging was set at 1µm for tracking. For long-term imaging, movies were acquired for 180-200 minutes with a frame rate of 40 seconds. Embryos were imaged with a temperature control unit set to 29°C.

### Image Analysis

#### Macrophage cell counts

Autofluorescence of the embryo was used to measure the position of the germband to determine the stages for analysis of fixed samples. Embryos with germband retraction of between 29-31% were assigned to Stage 11. Embryos with 35-40% retraction (Stage 12) were analysed for the number of macrophages that had entered the germband. Embryos with above 50-75% retraction were used for the number along the ventral nerve cord (vnc) and in the whole embryo. Macrophages were visualized using confocal microscopy with a Z-resolution of 2 µm and the number of macrophages within the germband or the segments of the vnc was calculated in individual slices (and then aggregated) using the Cell Counter plugin in FIJI. Total macrophage numbers were obtained using Imaris (Bitplane) by detecting all the macrophage nuclei as spots.

#### Macrophage tracking, speed, directionality and time for macrophage entry analysis

Embryos in which the macrophage nuclei were labeled with *srpHemo-H2A::3XmCherry* were imaged and 250×130×36 µm^3^ 3D-stacks were typically acquired with a constant 0.5×0.5×1 µm^3^ voxel size at every 40-41 seconds for approximately 3 hours. Images acquired from multiphoton microscopy were initially processed with InSpector software (LaVision Bio Tec) to compile channels from the imaging data (Table 3). Afterwards, the exported files were further processed using Imaris software (Bitplane) to visualize the recorded channels in 3D and the movie from each imaged embryo was rotated and aligned along the AP axis for further tracking analysis.

To analyze the movies by Imaris, the following analyses were applied:

i. To calculate the migration parameters while macrophages migrate from the head mesoderm to the yolk zone, movies were cropped in time to that period (typically 60 minutes from the original movie was used for analysis).
ii. To calculate the migration parameters of the macrophage moving on the yolk zone into the edge of germband, movies were acquired from the time point of the first macrophage appearing in the yolk zone and recorded until the onset of germband retraction.
iii. Macrophage nuclei were extracted using the spot detection function and tracks generated in 3D over time. We could not detect all macrophages in the head mesoderm as spots because of limitations in our imaging parameters. Tracks of macrophages which migrate towards the dorsal vessel, ventral nerve cord (vnc) and to the anterior of the head were omitted. The edge of the germband was detected using autofluorescence from the yolk and the mean position of the tracks in X- and Y-axis was used to restrict analysis to before macrophages reach the edge of the germband.
iv. Nuclei positions in XYZ-dimensions were determined for each time point and used for further quantitative analysis.
v. The time point when the first macrophage nucleus reached the germband was defined as T0 and the time point when the macrophage nucleus was within the germband and moved forward along the route between the ectoderm and mesoderm was taken as T1 and T1-T0 was defined as time for macrophage entry. T0 and T1 were determined by precisely examining macrophage position in xy and z dimensions (examination of individual 2 micron slices) over time.
vi. To measure the speed along the route between the germband edge and the yolk, tracks generated from macrophages from the time when the first macrophages started to move along the mentioned path until germband retraction onset were utilized.
vii. To calculate the speed of migration of the first or second macrophages in the germband the track generated for the first or second macrophages alone was used to obtain the nuclei position in XYZ-dimensions. Moreover, the average speed of the third through fifth macrophages moving along the same route was also measured. Speed was calculated within the first 30-35 µm of the patrh between the germband ectoderm and mesoderm. The mean position of the tracks in X- and Y-axis was used to restrict analysis to either of the migratory zones (head, yolk, germband entry, route along the germband ectoderm and mesoderm, route along the germband mesoderm and the yolk).

Macrophage migratory parameters, including cell speed and directionality (persistence), were calculated in Matlab (The MathWorks Inc.) from single cell positions in 3D for each time frame measured in Imaris (Bitplane), as described elsewhere (Smutny et al., 2017). Briefly, instantaneous velocities from single cell trajectories were averaged to obtain a mean instantaneous velocity value over the course of the measurement. To calculate directionality values, single cell trajectories were split into segments of equal length (*l* ;*l* = 10 frames) and calculated via a sliding window as the ratio of the distance between the macrophage start-to-end distance (*D*) over the entire summed distance covered by the macrophage between each successive frame (*d*_*i*_) in a segment. Calculated directionality values were averaged over all segments in a single trajectory and all trajectories were averaged to obtain a directionality index (*I*) for the duration of measurement (with 0 being the lowest and 1 the maximum directionality) as follows:

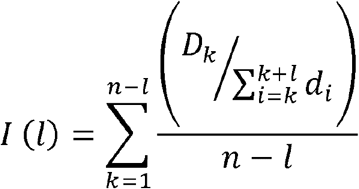

where *n* defines the total number of frames,*i* sum of frame-to-frame distances over one segment and *k* the sum over all segments of a trajectory.

Embryos from the control (*w*^*+*^; *+; srpHemo::3xmCherry)* and the CG9005 mutant *(w*^*+*^; *CG9005*^*BG02278*^; *srpHemo::3xmCherry)* were used for calculating the time for macrophage entry. Briefly, 100×130×34μm^3^ 3D-stacks were typically acquired with a constant 0.28×0.28×2μm^3^ voxel size at every 40-41 seconds for approximately 3 hours.

#### Cloning of constructs

Standard molecular biology methods were used and all constructs were sequenced by the Mycrosynth company (Vienna, Austria) before injection into flies. The enzymes *NotI*, T4 Polynucleotide Kinase (*PNK*) and *DpnI* were obtained from New England Biolabs, Ipswich, Massasuchetts, USA (Frankfurt, Germany). PCR amplifications were performed with GoTaq G2 DNA polymerase (Promega, Madison, USA) using a peqSTAR 2X PCR machine from PEQLAB, (Erlangen, Germany). All Infusion cloning was conducted using an Infusion HD Cloning kit (Clontech’s European distributer). The relevant oligo sequences were chosen using the Infusion primer Tool at the Clontech website (http://bioinfo.clontech.com/infusion/convertPcrsInit.do).

#### Construction of *srpHemo-*CG9005

A 3894 bp fragment containing the CG9005 ORF was amplified from the *UAS-CG9005::FLAG::HA* construct (Table S3) (*Drosophila* Genomics Resource Centre, DGRC) using relevant primers (Table S4). The fragment was cloned into the *srpHemo* plasmid (a gift from Katja Brückner (Brückner et al., 2004) after its linearization with *NotI*, using an Infusion HD cloning kit (Clontech’s European distributor).

#### Construction of *srpHemo-FAM214A* and *srpHemo-FAMB214B*

Fragments of 3225 bp and 1615 bp containing the FAM214A and FAMB214B ORFs, respectively, were amplified from cDNA prepared from dendritic cells (a gift from M. Sixt’s lab) with FAM214A Fwd and Rev primers, and with FAM214B Fwd and FAM214B Rev primers (Table S4). The fragments were cloned into the *srpHemo* plasmid using an Infusion HD cloning kit after its linearization with *NotI* (NEB).

#### Construction of mutant forms of *srpHemo-atossa*

Mutant forms of *atossa* (CG9005) were generated by removing the desired region from the CG9005 cDNA sequence by using inverse PCR followed by blunt end ligation and related primers (Table S4). Afterwards, *atossa* mutant constructs in the Bluescript vector were amplified and cloned into the *srpHemo* plasmid after its linearization with *NotI*, using an Infusion HD cloning kit.

#### Transgenic fly line production

The *srpHemo* and *UAS* constructs (Table S4) was injected into syncytial blastoderm stage embryos of M{3xP3-RFP.attP}ZH-86Fb (BL 24749) line (obtained from Peter Duchek of IMBA) to generate inserts on third chromosome by C31-mediated integration (Table S3) (Bischof et al., 2007; Gyoergy et al., 2018).

#### CRISPR sgRNA production and cloning

sgRNA target sequences for CRISPR-Cas9 based gene knock down for CG9253 (*porthos*) were designed as 20 nt sequences upstream of an NGG PAM motif in the *Drosophila* genome (https://www.flyrnai.org/crispr/) (Basset and Liu, 2014). The targeting oligonucleotides incorporated into *porthos* sgRNAs are given in (Table S4, The annealed oligo inserts were cloned into BspQ1-digested pAC-sgRNA-Cas9 vector (Addgene, plasmid # 49330) before transformation. Positive clones were confirmed by sequencing with pAC-sgRNA-Cas9-U6F primer (Table S4). All CRISPR-Cas9 constructs contain three distinct cassettes for expression of Cas9, an sgRNA against *porthos*, and a puromycin resistance marker.

#### Generation of *porthos* depleted S2R+ cells

To make the stable depleted cell lines, S2R+ Cells (2 × 10^5^) were seeded in Schneider medium plus 10% FCS (Gibco 21720024, Sigma F9665) in a 24-well plate. Plasmid sgRNA CRISPR *porthos* was co-transfected (1 µg of total DNA per well) with Effectene Tranfection Reagent (Qiagen, Hilden, Germany) following the manufacturer’s protocol. 4 hours after transfection the medium was changed and the cells were incubated for 72 hours at 25°C. Cells were then transferred to a 6-well plate before addition of 5µg/ml Puromycin. Selection with Puromycin took place for 7 days. Surviving cells were incubated without selection medium for 24 hours, after that they were added to 96-well cell culture plates in conditioned medium at a density of 1 cell/well. After 7 days we checked the wells for growing colonies to rule out that more than 1 colony was present per well. When cells were dense enough we first transferred them to a 24-, then a 12- and finally a 6-well plate. Once the cells reached confluency, we extracted the genomic DNA to perform a PCR-based prescreening of *porthos*-depleted cells to detect effective CRISPR (Table S4).

#### Quantitative Real Time-PCR (qRT-PCR) analysis

To verify the effective knockdown of genes, we first isolated RNA from S2R+ cells (1×10^7^ for the control and KD cells) according to the manufacturer’s protocol (Qiagen RNeas Mini Kit Cat No./ID: 74104). We used 500 ng of isolated RNA for cDNA synthesis, according to the manufacturer’s protocol (Qiagen Omniscript RT, Cat No./ID: 205111). Afterwards we performed qPCR to assess the mRNA expression of *atossa* and *porthos*, using *RpS20* as an internal control. Primer sequences for *Drosophila atossa* (CG9005) and *porthos* (CG9253) transcripts were designed using NCBI’s primer design tool (https://www.ncbi.nlm.nih.gov/tools/primer-blast/) and primer sequences for RpS20 gene, as an internal control gene, were obtained from the FlyPrimerBank (http://www.flyrnai.org/FlyPrimerBank) (Table S5). We amplified 4 µL cDNA (50 ng) using 10 µL of Takyon™ No Rox SYBR MasterMix Blue dTTP (Eurogentec, Liege, Belgium), 2 µL of each reverse and forward primers (10 mM). The thermal cycling conditions were as follows: 40 cycles of amplification each consisting of 10 s at 95°C, 15 s at 60°C and 10 s at 72°C, and cooling at 4°C. The experiments were carried out in technical triplicates and three biological replicates for each data point. The qPCR experiment was run on a LightCycler 480 (Roche, Basel, Switzerland) and data were analyzed in the LightCycler 480 Software and Prism (GraphPad Software). To calculate the fold change in *atossa* and *porthos* mRNA levels compared to the house-keeping gene mRNA levels, we averaged the Ct values of the technical replicates of each trial. We measured Δct by subtracting the housekeeping gene Ct average from the Ct average of *atossa* or *porthos*. Afterwards, the 2^^^- Δct was calculated for each trial.

### Polysome profiling in *porthos*-KD S2 cells

#### RNAi treatment of S2 cells

dsRNA for *porthos* (CG9253) was prepared as described by the SnapDragon manual (https://www.flyrnai.org/snapdragon). Briefly, template was prepared from S2 cell cDNA using the following primers designed using SnapDragon 5’-TAATACGACTCACTATAGGATAAG GAAGGGGACAGCGAG-3’ and the reverse primer: 5’-TAATACGACTCACTATAGGTTTGAAATGCCAGTTCCCTC-3’ both of which contain a T7 polymerase promoter. As a negative control, we made non-targeting dsRNA against GFP using the following primers: 5’-TAATACGACTCACTATAGGGGAGCGCACCATCTTCTTCAA-3’ and 5’-TAATACGACTCACTATAGGGCTGCTTGTCGGCCATGATATAG-3’. We performed *in vitro* transcription overnight at 37°C using the T7 Megascript kit (AM1334) following manufacturer’s instructions (Table S4). The RNA was treated with DNAse and purified using acid-phenol chloroform extraction and ethanol precipitated. The resulting RNA was annealed by heating at 65°C for 5 minutes and slow cooling to 37°C for an hour. Knocking down in S2 cells was performed using 1 µg of dsRNA as previously described (https://www.ncbi.nlm.nih.gov/pmc/articles/PMC4465107/). 0.5-1.0 × 10^6^ cells were seeded 30 minutes prior to transfection to adhere. Prior to transfection, the media was changed for 500 µl of fresh media. The seeded cells were treated with 500 µl of transfection complexes per well of a 6-well plate. 48 hours post transfection, cells were passaged to 10 cm dishes. After 3 more days cells were harvested for further analysis.

#### Polysome profiling and polysome sequencing

Polysome sequencing was performed as described by (Flora et al., 2018) with minor modifications. Cells were incubated with fresh medium 2-4 hours before harvesting. Cycloheximide (100 μg/ml) was first added to the medium for 3 min at RT, and the cells were subsequently centrifuged at 800 xg for 3 min. The cell pellet was afterwards washed two times with ice-cold phosphate-buffered saline (1X PBS, pH 7.4). The supernatant was discarded and the pellet was gently resuspended in 300 µl of lysis buffer A (300 mM NaCl, 15 mM Tris-HCl, pH 7.5, 15 mM EDTA, 1 mg/ml heparin, 1% Triton-X100, and 100 μg/ml cycloheximide) and lysed for 15 min on ice. The lysate was clarified by centrifugation at 8500 xg for 5 min at 4°C. 20% of the lysate was kept aside as an input. The clarified lysate was loaded onto a 10%-50% sucrose gradient in Buffer B (300 mM NaCl, 15 mM Tris-HCl, pH 7.5, 15 mM MgCl_2_, supplemented with 100 μg/ml cycloheximide) and centrifuged for 3 hours at 35,000 rpm in an SW41 rotor in a Beckman L7 ultracentrifuge (Beckman Coulter, Krefeld, Germany). The gradients were simultaneously fractionated on a Density Gradient Fractionation System (#621140007) at 0.75 ml/min. We added 20 μl of 20% SDS, 8 μl of 0.5 M pH 8 EDTA, and 16 μl of proteinase K (#P8107S) to each polysome fraction and incubated them for 30 min at 37°C. The RNA from each fraction was extracted by standard acid phenol: chloroform purification followed by 80% ethanol precipitation. The polysome fractions were then measured for RNA content and RNAseq libraries were prepared.

#### Polysome-seq library preparation and mRNA sequencing

The RNA was first treated with Turbo DNAse (TURBO DNA-free Kit, Life Technologies, AM1907) and then purified using DNAse Inactivation buffer. The RNA was then centrifuged for 1.5 min at 1000 xg and the supernatant was collected and centrifuged once more at the same condition. The RNA quantity was determined by measuring the absorbance at 260 nm (NanoDrop 2000 spectrophotometer; Peqlab).

Poly-A selection was performed according to manufacturer’s instructions (Bioo Scientific Corp., 710 NOVA-512991). Following Poly-A selection mRNA libraries were prepared according to manufacturer’s instructions (Bioo Scientific Corp., NOVA-5138-08), except that the RNA was incubated at 95°C for 13 min to generate optimal fragment sizes. The sequencing library quantity was determined using Qubit (Thermo Fisher Scientific). The library integrity was assessed with a Bioanalyzer 2100 system (RNA 6000 Pico kit, Agilent Technologies). The libraries on biological duplicates from each genotype were subjected to 75 base-pair single-end sequencing on Illumina NextSeq500 at the Center for Functional Genomics (CFG).

#### Data analysis of S2 cell polysome sequencing

First the reads were assessed for their quality using FastQC. Mapping of the reads was performed against *Drosophila* Genome (dm6.01, www.fruitfly.org) using Hisat version 2.1.0. Mapped reads were then assigned to feature using featureCount version v1.6.4. To calculate Translation efficiency (TE), TPMs (transcripts per million) values for polysome-libraries were calculated (Flora et al., 2018). All transcripts with zero reads were discarded from libraries for further analysis. The log2 ratio of TPMs between the polysome fraction and total mRNA was measured. This ratio represents TE. The TE value of each replicate was averaged and delta TE (ΔTE) was calculated as (*porthos* dsRNA TE)/(GFP dsRNA TE). Targets were defined as transcripts falling greater or less than two standard deviations (SD) from the median of ΔTE (Table S5).

#### Extracellular flux measurements for bioenergetic profiling

Cellular respiration was assessed using a Seahorse XF96 extracellular flux analyzer (Agilent Technologies, Santa Clara, CA USA). The oxygen consumption rate (OCR) as a measure of oxygen utilization of cells is an important indicator of mitochondrial function. The extracellular acidification rate (ECAR) is a measure of glycolytic activity measured via extracellular acidification due to lactate release, formed during the conversion of glucose to lactate during anaerobic glycolysis. Prior to measurement, wild-type and *porthos* KD cells were seeded at 10 × 10^5^ cells per well in Seahorse XF96 polystyrene tissue culture plates (Agilent) and incubated in unbuffered Seahorse RPMI assay medium (Agilent) supplemented with glucose (25 mM; Sigma-Aldrich), sodium pyruvate (1 mM; Gibco), and glutamine (2 mM; Gibco) in a non-CO2 incubator at 25°C and pH 7.4 for 1 h before the experiment. Cellular oxygen consumption was assessed in basal condition (prior to any addition) and after addition of oligomycin (2 μM; Agilent) Carbonyl cyanide-4 (trifluoromethoxy) phenylhydrazone (FCCP, 2 μM; Sigma-Aldrich), antimycin A and rotenone (both at 1 μM; Agilent). The three drugs were injected into the XF96 plate sequentially. This allowed for calculation of OCR linked to ATP production, maximal respiration capacity and spare respiratory capacity. Basal respiration was measured prior to injection of oligomycin A. Both OCR and ECAR were measured every 4 min with a mixing of 2 min in each cycle, with 4 cycles in total for the first step and 3 cycles thereafter.

Different parameters from the OCR graph were measured as follows. ATP turnover was calculated by subtracting the “last rate measurement before oligomycin” from the “minimum rate measurement after oligomycin injection”. Maximal respiration was defined as (maximum rate measurement after FCCP) - (non-mitochondrial respiration). Spare respiratory capacity (SRC) was measured by subtracting basal respiration from maximal respiration (Mookerjee et al., 2017).

#### Metabolomics profiling analysis

Samples for metabolomics were assessed by the VBCF metabolomics facility according to Rao et al. with slight modifications (https://www.viennabiocenter.org/facilities/metabolomics/) (Rao et al., 2019). 1 gr of wild-type or *atos* embryos were extracted using an ice-cold MeOH:ACN:H2O (2:2:1, v/v) solvent mixture. A volume of 1mL of cold solvent was added to each pellet, vortexed for 30 s, and incubated in liquid nitrogen for 1 min. The samples were thawed at room temperature and sonicated for 10 min. This cycle of cell lysis in liquid nitrogen combined with sonication was repeated three times. To precipitate proteins, the samples were incubated for 1 h at −20°C, followed by centrifugation at 13,000 rpm for 15 min at 4°C. The supernatant was removed and evaporated. The dry extracts were reconstituted in 100 μL of ACN:H2O (1:1, v/v), sonicated for 10 min, and centrifuged at 13,000 rpm for 15 min at 4°C to remove insoluble debris. The supernatants were transferred to Eppendorf tubes, shock frozen and stored at -80°C prior to LC/MS analysis. A volume of 1 μL of the metabolite extract was injected on a ZIC-pHILIC HPLC column operated at a flow rate of 100 μL/min, directly coupled to a TSQ Quantiva mass spectrometer (Thermo Fisher Scientific).

We used the following transitions for quantitation in the negative ion mode: AMP 346 m/z to 79 m/z, ADP 426 m/z to 134 m/z, ATP 506 m/z to 159 m/z, IMP 347 m/z to 79 m/z, GMP 362 m/z to 211 m/z, GDP 442 m/z to 344 m/z, GTP 522 m/z to 424 m/z, taurine 124 m/z to 80 m/z, malate 133 m/z to 115 m/z, citrate 191 m/z to 111 m/z, pyruvate 87 m/z to 43 m/z, lactate 89 m/z to 43 m/z, NADH 664 m/z to 408 m/z, NAD 662 m/z to 540 m/z, hexose phosphates 259 m/z to 97 m/z, Acetyl CoA 808 m/z to 408 m/z, CoA 766 m/z to 408 m/z, succinate 117 m/z to 73 m/z. Glutamine 147 m/z to 130 m/z, glutamate 148 m/z to 84 m/z, serine 106 m/z to 60 m/z were calculated in the positive ion mode. For all transitions, the optimal collision energy was defined by analyzing pure metabolite standards. Chromatograms were manually interpreted using trace finder (Thermo Fisher Scientific), validating experimental retention times with the respective quality controls. All measurements were within the linear range of detection.

For the metabolomics analysis, the metabolite concentration was normalized using a Z-score normalization method with the formula of y = (x−α)/λ, in which x refers to the real concentration, α indicates the mean value of all samples, and λ is the variance of all samples. The normalized concentrations of metabolites were applied to generate a heatmap, which showed the concentration difference of all metabolites. For KEGG (http://www.kegg.jp, Tokyo, Japan) pathway analysis, the clusterProfiler R package was employed.

#### Statistics and repeatability

Statistical tests as well as the number of embryos/ cells assessed are listed in the figure legends. All statistical analyses were performed using GraphPad Prism and significance was determined using a 95% confidence interval. Data points from individual experiments/ embryos were pooled to estimate mean and SEM. No statistical method was used to predetermine sample size and the experiments were not randomized. Unpaired t-test or Mann-Whitney was used to calculate the significance in differences between two groups and One-way Anova followed by Tukey post-test followed by Conover or Dunn’s post-test for multiple comparisons. All measurements were performed in 3-50 embryos. Representative images illustrated in Figures 1A-C, Figures 2B-C,E, Figures S2A-B, Figures 3B-D, Figure 4A,I, Figure S4B, and Figure 6D,G,I were from separate experiments that were repeated at least 3 and up to 7 times. Stills shown in Figure 1F, Figure S1I, Figure 4B, and Figure S4E are representative images from two-photon movies, which were repeated at least 3 times. Raw data from embryo scoring and analyzed tracking output from each movie is in Data S4.

## Exact genotype of *Drosophila* lines used in Figures

**Figure 1 and Figure S1**

**Figs. 1A-C:** Control: *w-; +; srpHemo-H2A::3xmCherry*, CG9005 mutant: *w-; P{EP}CG9005*^*BG02278*^; *srpHemo-H2A::3xmCherry*, CG9005 rescue: *w-; P{EP}CG9005*^*BG02278*^; *srpHemo-CG9005, srpHemo-H2A::3xmCherry*. **Fig. 1D:** Control: *w-; +; srpHemo-H2A::3xmCherry*, CG9005 mutant: *w-; P{EP}CG9005*^*BG02278*^; *srpHemo-H2A::3xmCherry*, Df1: *w-; P{EP}CG9005*^*BG02278*^*/ Df(2R)ED2222; srpHemo-H2A::3xmCherry*, Df2: *w-; P{EP}CG9005*^*BG02278*^*/Df(2R)BSC259; srpHemo-H2A::3xmCherry*, CG9005 rescue: *w-; P{EP}CG9005*^*BG02278*^; *srpHemo-CG9005, srpHemo-H2A::3xmCherry*. **Fig. 1E:** Control 1: *w*^*-*^ *P(w+)UAS-dicer/w-; P{attP,y[+],w[3’]/+; srpHemo-Gal4 UAS-GFP*, CG9005 RNAi 1: *UAS-Dicer2/w-; CG9005 RNAi* (*v106589*)*/+; srpHemo-Gal4 UAS-GFP, UAS-H2A::RFP/+*, Control 2: *w*^*-*^ *P(w+)UAS-dicer/w-; +; srpHemo-Gal4 UAS-GFP*, CG9005 RNAi 2: *UAS-Dicer2/ w-; CG9005 RNAi* (*v36080*)*/+; srpHemo-Gal4 UAS-GFP, UAS-H2A::RFP/+*, Control 3: *w*^*-*^ *P(w+)UAS-dicer****/****w-; P{attP,y[+],w[3’]/+; srpHemo-Gal4 UAS-GFP*, CG9005 RNAi 3: *UAS-Dicer2/w-; CG9005 RNAi* (*v33362*)*/+; srpHemo-Gal4 UAS-GFP, UAS-H2A::RFP/+*. **Figs. 1F-L:** Control: *w-; +; srpHemo-H2A::3xmCherry*, CG9005 mutant: *w-; P{EP}CG9005*^*BG02278*^; *srpHemo-H2A::3xmCherry*.

**Fig. S1A:** Control: *w-; +; srpHemo-H2A::3xmCherry*, mutant: *w-; P{EP}CG9005*^*BG02278*^; *srpHemo-H2A::3xmCherry*, Df1 cross: *w-; P{EP}CG9005*^*BG02278*^*/Df(2R)ED2222; srpHemo-H2A::3xmCherry*, Df2 cross: *w-; P{EP}CG9005*^*BG02278*^*/Df(2R)BSC259; srpHemo H2A::3xmCherry*, CG9005 rescue: *w-; P{EP}CG9005*^*BG02278*^; *srpHemo-CG9005, srpHemo-H2A::3xmCherry*. **Figs. S1B**,**H:** Control 1: *w*^*-*^ *P(w+)UAS-dicer/w-; P{attP,y[+],w[3’]/+; srpHemo-Gal4 UAS-GFP*, CG9005 RNAi 1: *UAS-Dicer2/w-; CG9005 RNAi* (*v106589*)*/+; srpHemo-Gal4 UAS-GFP, UAS-H2A::RFP/+*. Control 2: *w*^*-*^ *P(w+)UAS-dicer/w-; +; srpHemo-Gal4 UAS-GFP*, CG9005 RNAi 2: *UAS-Dicer2/w-; CG9005 RNAi* (*v36080*)*/+; srpHemo-Gal4 UAS-GFP, UAS-H2A::RFP/+*. Conrol 3: *w*^*-*^ *P(w+)UAS-dicer/w-; P{attP,y[+],w[3’]/+; srpHemo-Gal4 UAS-GFP*, CG9005 RNAi 3: *UAS-Dicer2/ w-; CG9005 RNAi* (*v33362*)*/+; srpHemo-Gal4 UAS-GFP, UAS-H2A::RFP/+*. **Figs. S1C**,**G:** Control: *w-; +; srpHemo-H2A::3xmCherry*, mutant: *w-; P{EP}CG9005*^*BG02278*^; *srpHemo-H2A::3xmCherry*. **Fig. S1D:** Control 1: *w*^*-*^ *P(w+)UAS-dicer/w-; P{attP,y[+],w[3’]/+; srpHemo-Gal4 UAS-GFP*, CG9005 RNAi 1: *UAS-Dicer2/w-; CG9005 RNAi* (*v106589*)*/+; srpHemo-Gal4 UAS-GFP, UAS-H2A::RFP/+*. **Fig. S1E:** Control 2: *w*^*-*^ *P(w+)UAS-dicer/w-; +; srpHemo-Gal4 UAS-GFP*, CG9005 RNAi 2: *UAS-Dicer2/w-; CG9005 RNAi* (*v36080*)*/+; srpHemo-Gal4 UAS-GFP, UAS-H2A::RFP/+*. **Fig. S1F:** Conrol 3: *w*^*-*^ *P(w+)UAS-dicer/w-; P{attP,y[+],w[3’]/+; srpHemo-Gal4 UAS-GFP*, CG9005 RNAi 3: *UAS-Dicer2/w-; CG9005 RNAi* (*v33362*)*/+; srpHemo-Gal4 UAS-GFP, UAS-H2A::RFP/+*. **Figs. S1I-L:** Control: *w-; +; srpHemo-H2A::3xmCherry*, CG9005 mutant: *w-; P{EP}CG9005*^*BG02278*^; *srpHemo-H2A::3xmCherry*.

**Figure 2 and Figure S2**

**Fig. 2B:** *w-;+; UAS-atossa::FLAG::HA, srpHemo-Gal4, srpHemo-H2A::3xmCherry*. **Figs. 2C**,**D:** Control: *w-; +; srpHemo-H2A::3xmCherry, atos* mutant: *w-; atossa* ^*BG02278*^; *srpHemo-H2A::3xmCherry*, Atossa rescue: *w-; atossa* ^*BG02278*^; *srpHemo-atossa, srpHemo-H2A::3xmCherry*, rescue: *w-; atossa* ^*BG02278*^; *srpHemo-atossa* ^*DUF4210-*^, *srpHemo-H2A::3xmCherry*, rescue: *w-; atossa* ^*BG02278*^; *srpHemo-atossa* ^*CherSeg-*^, *srpHemo-H2A::3xmCherry*, rescue: *w-; atossa* ^*BG02278*^; *srpHemo-atossa* ^*DU4210-/CherSeg-*^, *srpHemo-H2A::3xmCherry*, rescue: *w-; atossa* ^*BG02278*^; *srpHemo-atossa* ^*TAD1-/TAD2-*^, *srpHemo-H2A::3xmCherry*. **Figs. 2E**,**F:** Control: *w-; +; srpHemo-H2A::3xmCherry*, mutant: *w-; atossa*^*BG02278*^; *srpHemo-H2A::3xmCherry*, rescue: *w-; atossa*^*BG02278*^; *srpHemo-FAM214A, srpHemo-H2A::3xmCherry*, rescue: *w-; atossa*^*BG02278*^; *srpHemo-FAM214B, srpHemo-H2A::3xmCherry*.

**Figs. S2B:** Rescue: *w-; atossa*^*BG02278*^; *srpHemo-atossa* ^*TAD1-*^, *srpHemo-H2A::3xmCherry*, rescue: *w-; atossa*^*BG02278*^; *srpHemo-atossa* ^*TAD2-*^, *srpHemo-H2A::3xmCherry*. **Fig. S2C:** Control: *w-; +; srpHemo-H2A::3xmCherry, atos* mutant: *w-; atossa*^*BG02278*^; *srpHemo-H2A::3xmCherry*, Atossa rescue: *w-; atossa*^*BG02278*^; *srpHemo-atossa, srpHemo-H2A::3xmCherry*, rescue: *w-; atossa*^*BG02278*^; *srpHemo-atossa* ^*TAD1-*^, *srpHemo-H2A::3xmCherry*, rescue: *w-; atossa*^*BG02278*^; *srpHemo-atossa* ^*TAD2-*^, *srpHemo-H2A::3xmCherry*. **Fig. S2D:** Control: *w-; +; srpHemo-H2A::3xmCherry*, mutant: *w-; atossa*^*BG02278*^; *srpHemo-H2A::3xmCherry*, rescue: *w-; atossa*^*BG02278*^; *srpHemo-atossa, srpHemo-H2A::3xmCherry*, rescue: *w-; atossa*^*BG02278*^; *srpHemo-atossa* ^*DUF4210-*^, *srpHemo-H2A::3xmCherry*, rescue: *w-; atoss* ^*BG02278*^; *srpHemo-atossa* ^*CherSeg-*^, *srpHemo-H2A::3xmCherry*, rescue: *w-; atossa*^*BG02278*^; *srpHemo-atossa*^*DUF4210-/CherSeg-*^, *2srpHemo-H2A::3xmCherry*, rescue: *w-; atossa*^*BG02278*^; *srpHemo-atossa*^*TAD1-*^, *srpHemo-H2A::3xmCherry*, rescue: *w-; atossa*^*BG02278*^; *srpHemo-atossa*^*TAD2-*^, *srpHemo-H2A::3xmCherry*, rescue: *w-; atossa*^*BG02278*^; *srpHemo-atossa*^*TAD1-/2-*^, *srpHemo-H2A::3xmCherry*. **Fig. S2E:** Control: *w-; +; srpHemo-H2A::3xmCherry, atos* mutant: *w-; atossa*^*BG02278*^; *srpHemo-H2A::3xmCherry*, Atossa rescue: *w-; atossa*^*BG02278*^; *srpHemo-atossa, srpHemo-H2A::3xmCherry*, rescue: *w-; atossa*^*BG02278*^; *srpHemo-FAM214A, srpHemo-H2A::3xmCherry*, rescue: *w-; atossa*^*BG02278*^; *srpHemo-FAM214B, srpHemo-H2A::3xmCherry*.

**Figure 3 and Figure 3S**

**Figs. 3B**,**F:** Control (for *porthos* or *CG9253*): *w/y,w[1118]; P{attP,y[+],w[3’]}; srpHemo-Gal4, srpHemo-H2A::3xmCherry/+*, CG9253 RNAi (*porthos*): *w-; porthos RNAi* (*v36589*)*/+; srpHemo-Gal4, srpHemo-H2A::3xmCherry/+*. **Fig. 3C:** Control 1 (for CG9331 or *GR/HPR*): *w/y,w[1118]; P{attP,y[+],w[3’]}; srpHemo-Gal4, srpHemo-H2A::3xmCherry/+*, CG9331 RNAi 1 (GR/HPR): *UAS-Dicer2/ w-; GR/HPR RNAi* (*v44653*)*/+; srpHemo-Gal4, srpHemo-H2A::3xmCherry/+*. **Fig. 3D:** Control 1 (for CG7144 or *LKR/SDH*): *w/y,w[1118]; P{attP,y[+],w[3’]}; srpHemo-Gal4, srpHemo-H2A::3xmCherry/+*, CG7144 RNAi 1 (*LKR/SDH*): *UAS-Dicer2/ w-; LKR/SDH RNAi* (*v51346*)*/+; srpHemo-Gal4, srpHemo-H2A::3xmCherry/+*. **Fig. 3F:** Control 1: *w/y,w[1118]; P{attP,y[+],w[3’]}; srpHemo-Gal4, srpHemo-H2A::3xmCherry/+*, CG9331 RNAi 1 *(GR/HPR*): *UAS-Dicer2/ w-; GR/HPR RNAi* (*v44653*)*/+; srpHemo-Gal4, srpHemo-H2A::3xmCherry/+, Control* 2: *w/y,w[1118]; P{attP,y[+],w[3’]}; srpHemo-Gal4, srpHemo-H2A::3xm-Cherry/+*, CG9331 RNAi 2 (*GR/HPR*): *UAS-Dicer2/ w-; GR/HPR RNAi* (*v10780*)*/+; srpHemo-Gal4, srpHemo-H2A::3xmCherry/+*, Control 3: *w/y,w[1118]; P{attP,y[+],w[3’]}; srpHemo-Gal4, srpHemo-H2A::3xmCherry/+*, CG9331 RNAi 3 *(GR/HPR): UAS-Dicer2/ w-; GR/HPR RNAi* (*64652*)*/+; srpHemo-Gal4, srpHemo-H2A::3xmCherry/+*. **Fig. 3G:** Control 1: *w/y,w[1118]; P{attP,y[+],w[3’]}; srpHemo-Gal4, srpHemo-H2A::3xmCherry/+*, CG7144 RNAi 1 (*LKR/SDH*): *UAS-Dicer2/ w-; LKR/SDH RNAi (v51346)/+; srpHemo-Gal4, srpHemo-H2A::3xmCherry/+*, Control 2: *w/y,w[1118]; P{attP,y[+],w[3’]}; srpHemo-Gal4, srpHemo-H2A::3xmCherry/+*, CG7144 RNAi 2 *(LKR/SDH): UAS-Dicer2/ w-; LKR/SDH RNAi* (*v109650*)*/+; srpHemo-Gal4, srpHemo-H2A::3xmCherry/+*.

**Figs. S3A-B:** Control: *w-; +; srpHemo-H2A::3xmCherry*, mutant: *w-; P{EP}CG9005*^*BG02278*^; *srpHemo-H2A::3xmCherry*. **Fig. S3D:** Control 1: *w/y,w[1118]; P{attP,y[+],w[3’]}; srpHemo-Gal4, srpHemo-H2A::3xmCherry/+*, CG2137 RNAi 1 (*Gpo2*): *w-/y,w[1118]; Gpo2 RNAi* (*v41234*)*/+; srpHemo-Gal4, srpHemo-H2A::3xmCherry/+*, Control 2: *w/y,w[1118]; P{attP,y[+],w[3’]}; srpHemo-Gal4, srpHemo-H2A::3xmCherry/+*, CG2137 RNAi 2 (*Gpo2*): *w-/y,w[1118]; Gpo2 RNAi* (*68145*)*/+; srpHemo-Gal4, srpHemo-H2A::3xmCherry/+*. **Fig. S3E:** Control 1: *w/y,w[1118]; P{attP,y[+],w[3’]}; srpHemo-Gal4, srpHemo-H2A::3xmCherry/+*, CG11061 RNAi 1 (*GM130*): *w-/y,w[1118]; GM130 RNAi* (*v330284*)*/+; srpHemo-Gal4 UAS-GFP, UAS-H2A::RFP/+*, Control 2: *w/y,w[1118]; P{attP,y[+],w[3’]}; srpHemo-Gal4, srpHemo-H2A::3xmCherry/+*, CG11061 RNAi 2 (*GM130*): *w-/y,w[1118]; GM130 RNAi* (*64920*)*/+; srpHemo-Gal4, srpHemo-H2A::3xmCherry/+*. **Fig. S3F:** Control (for CG9253 or *porthos*): *w/y,w[1118]; P{attP,y[+],w[3’]}; srpHemo-Gal4, srpHemo-H2A::3xmCherry/+*, CG9253 RNAi (*porthos*): *w-; porthos RNAi* (*v36589)/+; srpHemo-Gal4, srpHemo-H2A::3xmCherry/+*. **Fig. S3G:** Control 1: *w/y,w[1118]; P{attP,y[+],w[3’]}; srpHemo-Gal4, srpHemo-H2A::3xmCherry/+, CG9331* RNAi 1 (*GR/HPR*): *UAS-Dicer2/ w-; GR/HPR RNAi* (*v44653*)*/+; srpHemo-Gal4, srpHemo-H2A::3xmCherry/+, Control* 2: *w/y,w[1118]; P{attP,y[+],w[3’]}; srpHemo-Gal4, srpHemo-H2A::3xmCherry/+*, CG9331 RNAi 2 (*GR/HPR*): *UAS-Dicer2/ w-; GR/HPR RNAi* (*v10780*)*/+; srpHemo-Gal4, srpHemo-H2A::3xmCherry/+, Control 3*: *w/y,w[1118]; P{attP,y[+],w[3’]}; srpHemo-Gal4, srpHemo-H2A::3xmCherry/+*, CG9331 RNAi 3 (*GR/HPR*): *UAS-Dicer2/ w-; GR/HPR RNAi (64652)/+; srpHemo-Gal4, srpHemo-H2A::3xmCherry/+*. **Fig. S3H:** Control 1: *w/y,w[1118]; P{attP,y[+],w[3’]}; srpHemo-Gal4, srpHemo-H2A::3xmCherry/+*, CG7144 RNAi 1 (*LKR/SDH*): *UAS-Dicer2/ w-; LKR/SDH RNAi* (*v51346*)*/+; srpHemo-Gal4, srpHemo-H2A::3xmCherry/+*, Control 2: *w/y,w[1118]; P{attP,y[+],w[3’]}; srpHemo-Gal4, srpHemo-H2A::3xmCherry/+*, CG7144 RNAi 2 (*LKR/SDH*): *UAS-Dicer2/w-; LKR/SDH RNAi* (*v109650*)*/+; srpHemo-Gal4, srpHemo-H2A::3xmCherry/+*.

**Figure 4 and Figure 4S**

**Fig. 4A:** *w-;+; UAS-porthos::FLAG::HA, srpHemo-Gal4, srpHemo::3xmCherry*. **Figs. 4B-H:** Control: *w/y,w[1118]; P{attP,y[+],w[3’]}; srpHemo-Gal4, srpHemo-H2A::3xmCherry/+*, CG9253 RNAi (*porthos*): *w-; porthos RNAi* (*v36589*)*/+; srpHemo-Gal4, srpHemo-H2A::3xmCherry/+*. **Figs. 4I-J:** Control: *w-; +; srpHemo-H2A::3xmCherry, atos* mutant: *w-; atossa*^*BG02278*^; *srpHemo-H2A::3xmCherry*, Atos rescue: *w-; atossa*^*BG02278*^; *UAS-atossa::FLAG::HA, srpHemo-Gal4, srpHemo-H2A::3xmCherry*, rescue: *w-; atossa*^*BG02278*^; *UAS-porthos::FLAG::HA, srpHemo-Gal4, srpHemo-H2A::3xmCherry*.

**Figs. 4SC-H:** Control: *w/y,w[1118]; P{attP,y[+],w[3’]}/+; srpHemo-Gal4, srpHemo-H2A::3xmCherry/+*, CG9253 RNAi (*porthos*): *w-; porthos RNAi* (*v36589*)*/+; srpHemo-Gal4, srpHemo-H2A::3xmCherry/+*.

**Figure 6 and Figure 6S**

**Fig. 6D-F:** Control: *w-; +; srpHemo-Gal4, srpHemo-H2A::3xmCherry*, dominant negative inhibitor of Complex V (CV-DN): *w-;UAS-CVDN; srpHemo-Gal4, srpHemo-H2A::3xmCherry*. **Figs. 6G-H:** Control: *w-; P{attP,y[+],w[3’]}/+; srpHemo-Gal4, srpHemo-H2A::3xmCherry*, Complex III (Cyt-c1, CG4769) RNAi 1: *w-; cyt-c1 RNAi* (*v109809*)*/+; srpHemo-Gal4, srpHemo-H2A::3xmCherry*, Complex III (UQCR-cp1, CG3731) RNAi 2: *w-; UQCR-cp1 RNAi (v101350)/+; srpHemo-Gal4, srpHemo-H2A::3xmCherry*, Complex III (UQCR-cp2, CG4169) RNAi 3: *w-; UQCR-cp2 RNAi* (*v100818*)*/+; srpHemo-Gal4, srpHemo -H2A::3xmCherry*, Complex V (ATP synthase F1F0, CG3612) RNAi: *w-; RNAi* (*v34664*)*/+; srpHemo-Gal4, srpHemo-H2A::3xmCherry*. **Fig. 6J:** Control: *w-;* +; *srpHemo-Gal4, srpHemo-3xmCherry, atos* mutant: *w-; atossa*^*BG02278*^; *srpHemo-Gal4, srpHemo-3xmCherry*, Control: *w/y,w[1118]; P{attP,y[+],w[3’]};srpHemo-Gal4, srpHemo-3xmCherry/+*, CG9253 RNAi (*porthos*): *w-; porthos RNAi* (*v36589*)*/+; srpHemo-Gal4, srpHemo-3xmCherry/+*, Control: *w-; +; srpHemo-Gal4, srpHemo-3xmCherry, CV-DN*: *w-;UAS-CV DN; srpHemo-Gal4, srpHemo-3xmCherry*.

**Fig. 6SF:** Control: *w-; P{attP,y[+],w[3’]/+; srpHemo-Gal4, srpHemo-H2A::3xmCherry*, Complex III (Cyt-c1, CG4769) RNAi 1: *w-; cyt-c1 RNAi* (*v109809*)*/+; srpHemo-Gal4, srpHemo-H2A::3xmCherry*, Complex III (UQCR-cp1, CG3731) RNAi 2: *w-; UQCR-cp1 RNAi (v101350)/+; srpHemo-Gal4, srpHemo-H2A::3xmCherry*, Complex III (UQCR-cp2, CG4169) RNAi 3: *w-; UQCR-cp2 RNAi* (*v100818*)*/+; srpHemo-Gal4, srpHemo-H2A::3xmCherry*, Complex V (ATP synthase F1F0, CG3612) RNAi: *w-; CG3612 RNAi* (*v34664*)*/+; srpHemo-Gal4, srpHemo-H2A::3xmCherry*. **Figs. 6SG-H:** *w-;* +; *srpHemo-Gal4, srpHemo-3xmCherry, atos* mutant: *w-; atossa*^*BG02278*^; *srpHemo-Gal4, srpHemo-3xmCherry*, Control: *w/y,w[1118]; P{attP,y[+],w[3’]};srpHemo-Gal4, srpHemo-3xmCherry/+*, CG9253 RNAi (*porthos*): *w-; porthos RNAi* (*v36589*)*/+; srpHemo-Gal4, srpHemo-H2A::3xmCherry/+*, Control: *w-; +; srpHemo-Gal4, srpHemo-3xmCherry, CV-DN*: *w-;UAS-CV DN; srpHemo-Gal4, srpHemo-3xmCherry*

**Figures 7 and S7:**

**Figs. 7B-H, SB-I:** Control: *w-; +; srpHemo-3xmCherry*, mutant: *w-; atossa*^*BG02278*^; *srpHemo-3xmCherry*.

## Resource Availability

Fly lines, plasmids and other reagents utilized are available upon request from the Lead contact

Original reads from RNA sequencing and Polysome profiling has been deposited at: (will be done once paper is in revision).

## Notes

### Competing Interest Statement

The authors have declared no competing interest.

