## Supplemental Figures and Legends for "A genetic program boosts mitochondrial function to power macrophage tissue invasion"

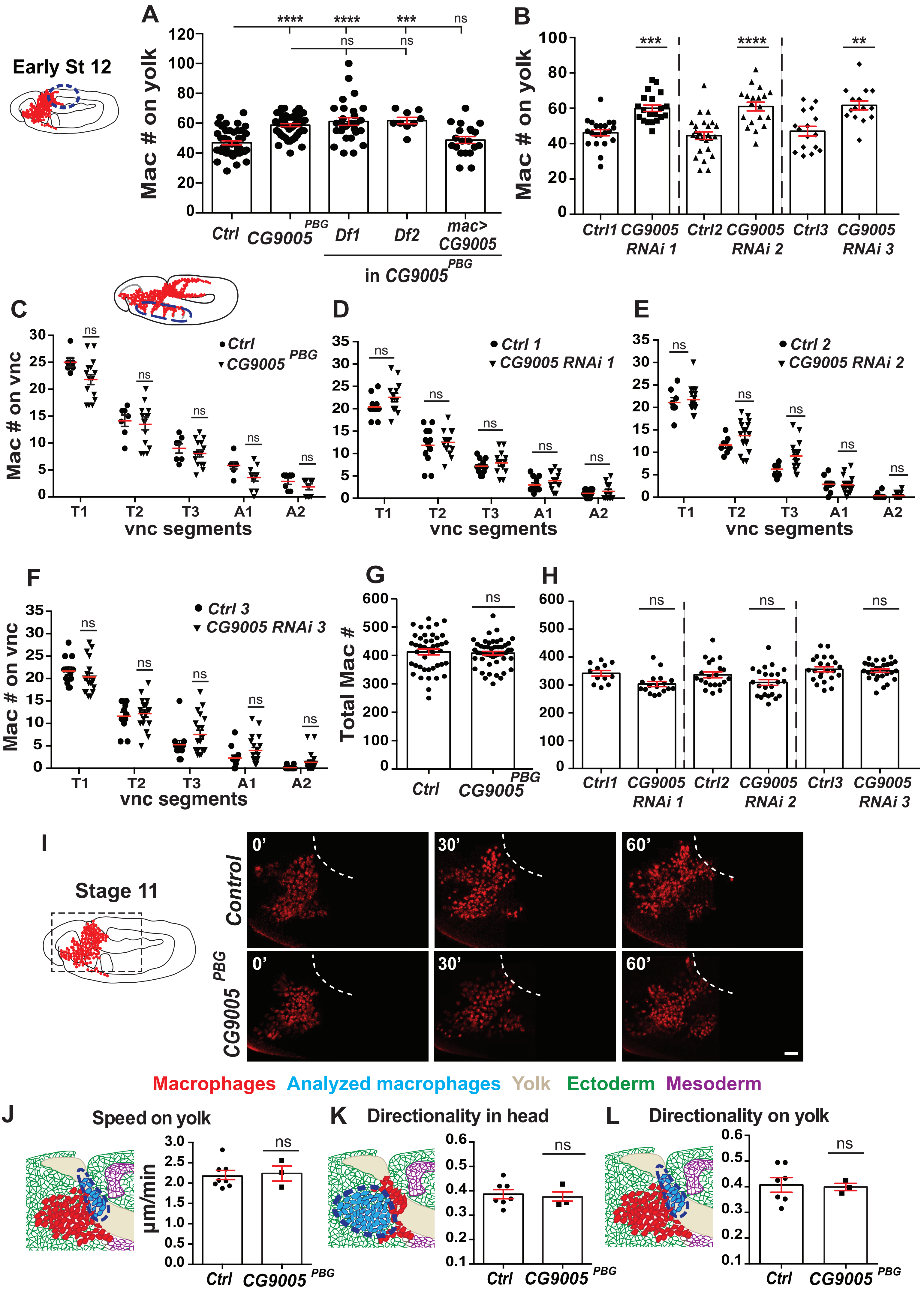
A
Early St 12
Mac # on yolk
Ctrl
CG9005 PBG
Df1
Df2
mac> CG9005
in CG9005 PBG
****
***
ns
B
Ctrl1
CG9005 RNAi 1
Ctrl2
CG9005 RNAi 2
Ctrl3
CG9005 RNAi 3
**
C
Mac # on vnc
vnc segments
T1
T2
T3
A1
A2
D
Ctrl 1
CG9005 RNAi 1
E
Ctrl 2
CG9005 RNAi 2
F
Ctrl 3
CG9005 RNAi 3
G
Total Mac #
H
I
Stage 11
Control
0'
30'
60'
Macrophages
Analyzed macrophages
Yolk
Ectoderm
Mesoderm
J
Speed on yolk
μm/min
K
Directionality in head
L
Directionality on yolk

**Figure S1 related to Figure 1: *CG9005$^{PBG}$* mutant macrophages migrate normally within the head and along the vnc. Fig S1A-B.** Quantification of macrophages on the yolk in fixed early Stage 12 embryos shows a significant increase in (**A**) the *P{GT1}CG9005$^{BG02278}$* P element mutant (*CG9005$^{PBG}$*) and in (**B**) lines expressing each of the CG9005 RNAis in macrophages compared to the control. (**A**): control n=43, mutant n=50, mutant/*Df1* n=28, mutant/*Df2* n=9, rescue=20; p<0.0001 for control vs mutant, p=0.99 for control vs rescue, p=0.001 for mutant vs rescue. (**B**): control 1 n=21, CG9005 *RNAi 1* n=20, p=0.0002; control 2 n=25, CG9005 *RNAi 2* n=19, p<0.0001; control 3 n=16, CG9005 *RNAi 3* n=15, p=0.001). **Fig S1C-F.** Macrophage quantification in ventral nerve cord (vnc) segments reveals no significant difference in macrophage migration along the vnc between *CG9005$^{PBG}$* mutant (n=15) and control embryos (n=7, p>0.05) or *srpHemo>CG9005 RNAi* embryos compared to the controls (control 1 n=8, CG9005 *RNAi 1* n=13, p=0.25; control 2 n=8, CG9005 *RNAi 2* n=16, p=0.5; control 3 n=8, CG9005 *RNAi 3* n=16, p>0.99). **Fig S1G-H.** Quantification of the total macrophage number reveals no significant difference between the control (n=43) and *CG9005$^{PBG}$* mutant embryos (n=50, p=0.69), or the control and *srpHemo>CG9005 RNAi* embryos (control 1 n=12, *CG9005 RNAi 1* n=17, p=0.9; control 2 n=27, *CG9005 RNAi 2* n=19, p=0.84; control 3 n=23, *CG9005 RNAi 3* n=27, p=0.16). **Fig S1I.** Stills from two-photon movies of control and *CG9005$^{PBG}$* mutant embryos, showing macrophages migrating starting at Stage 10 from the head towards the germband. Elapsed time indicated in minutes. The germband edge (white dotted line) was detected by yolk autofluorescence. **Fig S1J-L.** Quantification of migration parameters from two-photon live imaging of macrophages. **Fig S1J.** Macrophages on the yolk sac in the *CG9005$^{PBG}$* mutant reach the germband with a similar speed to control macrophages. Speed: control and mutant=2.2 µm/min; movie #: control=8, mutant=3; track #: control=373, mutant=124, p=0.78. **Fig S1K-L.** Macrophage directionality (**K**) in the head or (**L**) on the yolk sac shows no change in the *CG9005$^{PBG}$* mutant compared to the control. Head directionality: control=0.39, mutant=0.37, p=0.74; yolk sac directionality: control=0.40, mutant=0.39, p=0.86. Macrophages analyzed in **A-L** were labeled with *srpHemo-H2A::3xmCherry* to visualize nuclei. In schematics, macrophages are shown in red and analyzed macrophages in light blue, the ectoderm in green, the mesoderm in purple, and the yolk in beige. Throughout this work embryos were staged for imaging and quantification based on germband retraction away from the anterior of less than 29% for stage 10, 29%–31% for stage 11, and 35%–40% for stage 12. In all figures histograms show mean±SEM, ns=p>0.05, *p<0.05, **p<0.01, ***p<0.001, ****p<0.0001. One-way ANOVA with Tukey for (**A**) and unpaired t test for (**B-H**) and (**J-L**).

Figure. S2

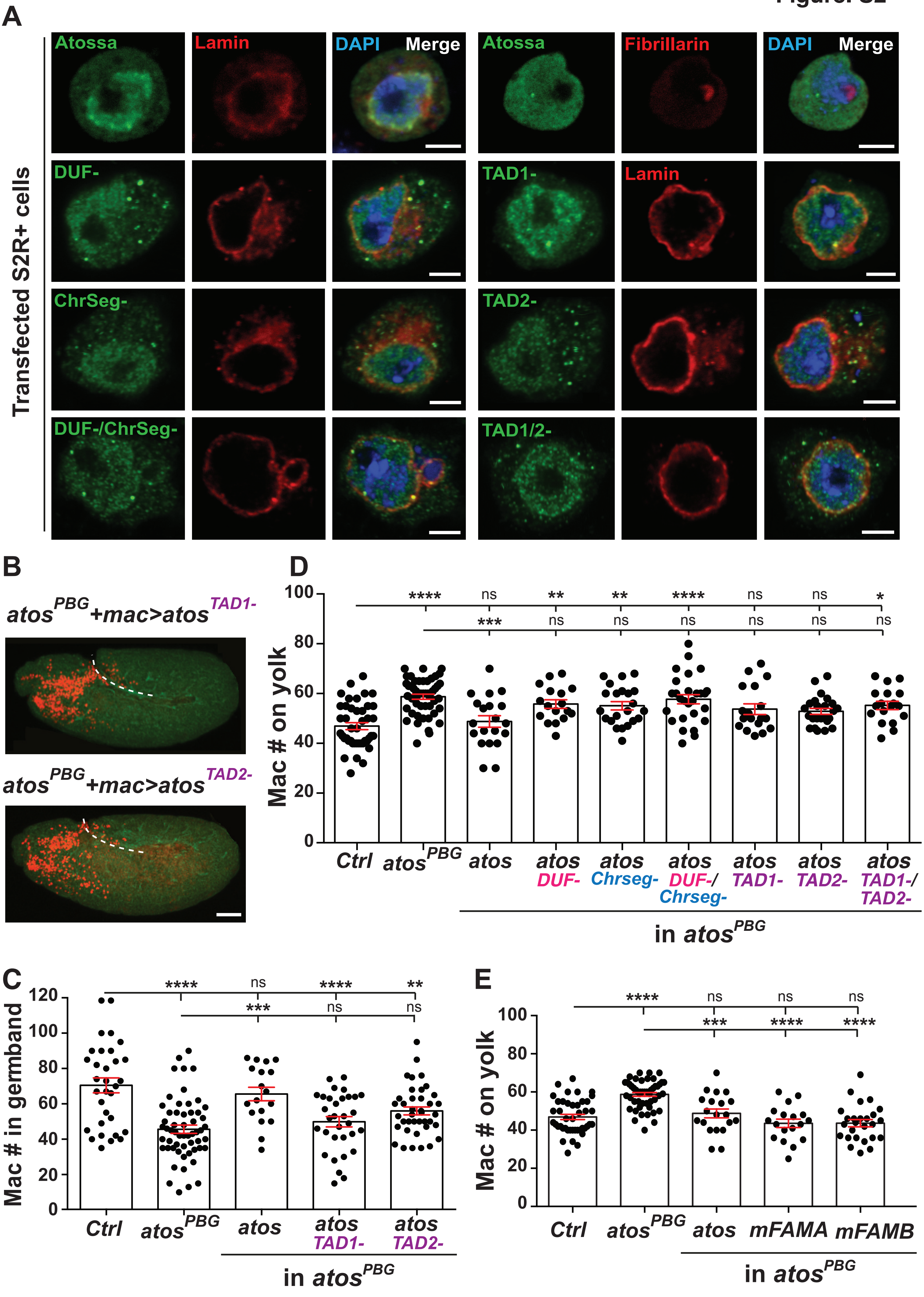

A
Atossa
Lamin
DAPI
Merge
Atossa
Fibrillarin
DAPI
Merge
Transfected S2R+ cells
DUF-
TAD1-
Lamin
ChrSeg-
TAD2-
DUF-/ChrSeg-
TAD1/2-
B
atos PBG +mac>atos TAD1-
atos PBG +mac>atos TAD2-
C
Mac # in germband
****
ns
****
**
***
ns
ns
120
100
80
60
40
20
0
Ctrl
atos PBG
atos
atos TAD1-
atos TAD2-
in atos PBG
D
Mac # on yolk
****
ns
**
**
****
ns
ns
*
***
ns
ns
ns
ns
ns
ns
100
80
60
40
20
0
Ctrl
atos PBG
atos
atos DUF-
atos Chrseg-
atos DUF-/ Chrseg-
atos TAD1-
atos TAD2-
atos TAD1-/ TAD2-
in atos PBG
E
Mac # on yolk
****
ns
ns
ns
***
****
****
100
80
60
40
20
0
Ctrl
atos PBG
atos
mFAMA
mFAMB
in atos PBG

**Figure S2 related to Figure 2. Atos's TAD domains are essential in macrophages for their tissue infiltration. Fig. S2A.** S2R+ cells were transfected with wild type Atos or forms lacking the indicated domains. HA tagged Atos (green), the nuclear membrane marker Lamin (red) and the nucleolar marker Fibrillarin (red) were visualized with antibodies, and nuclear DNA with DAPI (blue). All forms of Atos are expressed under direct control of the *srpHemo* promoter. **Fig S2B.** Representative confocal images of Stage 12 embryos from $atos^{PBG}$ mutants expressing Atos lacking either TAD1 or 2 in macrophages from the *srpHemo* promoter. Macrophages (red) were visualized with *srpHemo-H2A::3xmCherry* expression and the embryo outlines with phalloidin staining to detect actin (green). **Fig S2C.** Quantification shows that deletion of TAD1 or 2 blocks Atos's ability to rescue the germband migration defect of Stage 12 $atos^{PBG}$ mutant embryos upon expression in macrophages. Control n=32, mutant n=56, WT rescue n=18, $TAD1^-$ n=32, $TAD2^-$ n=39. For control vs WT rescue $p>0.99$, for control vs $TAD1^-$ rescue $p<0.0001$, and for control $TAD2^-$ rescue $p=0.003$. **Fig S2D.** Quantification in fixed early Stage 12 embryos shows a significant increase in the number of macrophages on the yolk in the $atos^{PBG}$ mutant and $atos^{PBG}$ expressing forms of *atos* lacking either DUF4210, ChrSeg, both DUF4210 and ChrSeg, or TAD1 and TAD2 compared to control embryos and $atos^{PBG}$ embryos expressing WT Atos. Control n=43, *atos* mutant n=50, WT *atos* rescue n=20, $DUF4210^-$ rescue n=17, $ChrSeg^-$ rescue n=22, $DUF4210^-/ChrSeg^-$ rescue n=27, $TAD1^-$ rescue n=18, $TAD2^-$ rescue n=24, $TAD1^-/TAD2^-$ rescue n=18. For control vs *atos* mutant $p<0.0001$, for control vs WT rescue $p>0.99$, for control vs. other rescues expressing *atos* lacking conserved motifs $p>0.1$. **Fig S2E.** Quantification shows a similar number of macrophages on the yolk in fixed early Stage 12 $atos^{PBG}$ mutant embryos which express *mFAM214A* or *mFAM214B* in macrophages compared to the control. Control n=43, mutant n=50, WT rescue n=20, *mFAM214A* rescue n=18, *mFAM214B* rescue n=26. For control vs $atos^{PBG}$ $p=0.93$, for control vs *mFAM214A* rescue $p=0.65$, for control vs *mFAM214B* rescue $p=0.56$, for $atos^{PBG}$ mutant vs $atos^{PBG}$, *mFAM214A* and *mFAM214B* rescues $p<0.0001$. One-way ANOVA with Tukey for (**C**-**E**). Scale bars: 3 µm in (**A**), 50 µm in (**B**).

Figure. S3

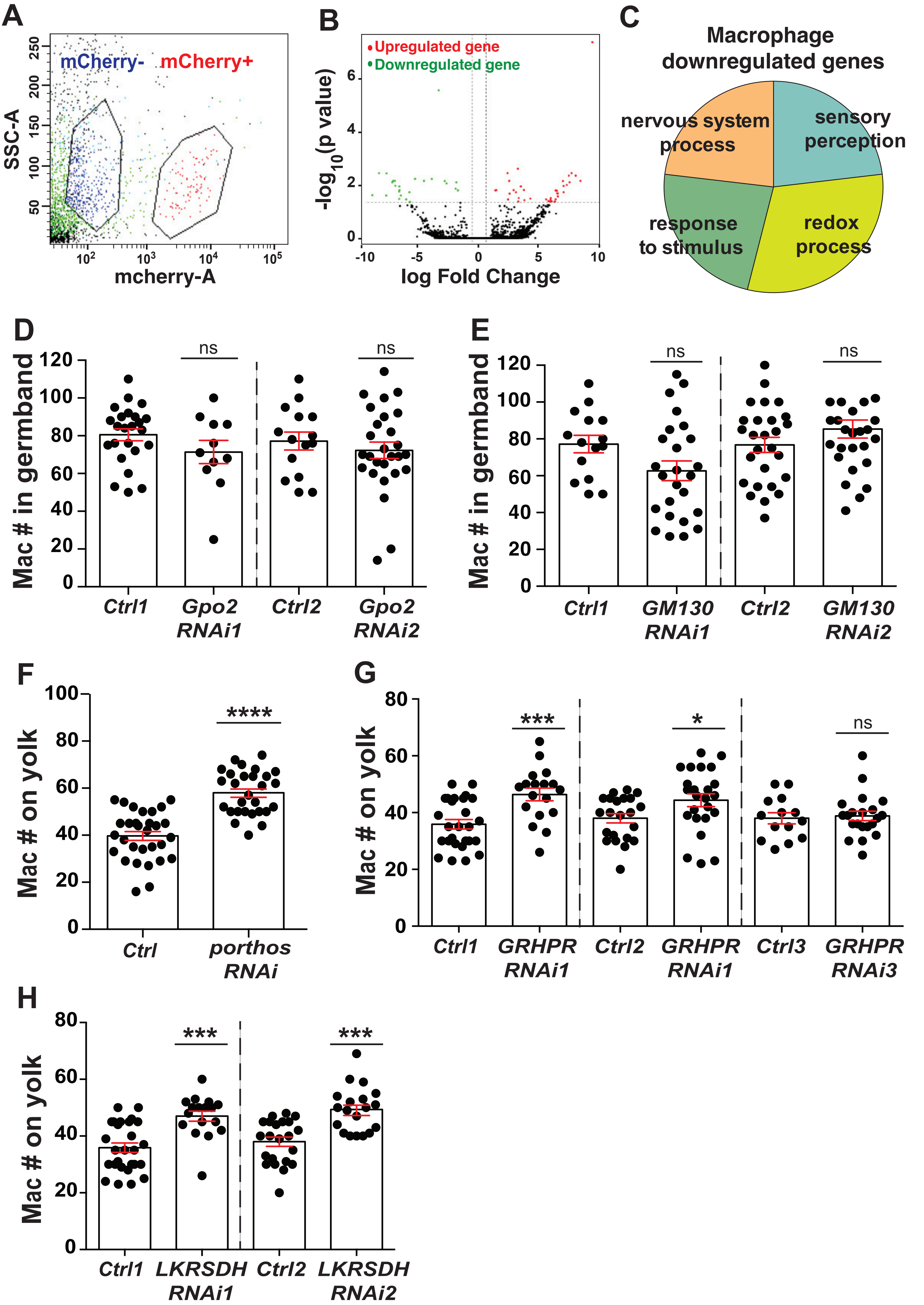

A
mCherry-
mCherry+
SSC-A
mcherry-A
B
Upregulated gene
Downregulated gene
-log10(p value)
log Fold Change
C
Macrophage downregulated genes
nervous system process
sensory perception
response to stimulus
redox process
D
Mac # in germband
ns
ns
Ctrl1
Gpo2 RNAi1
Ctrl2
Gpo2 RNAi2
E
Mac # in germband
ns
ns
Ctrl1
GM130 RNAi1
Ctrl2
GM130 RNAi2
F
Mac # on yolk
****
Ctrl
porthos RNAi
G
Mac # on yolk
***
*
ns
Ctrl1
GRHPR RNAi1
Ctrl2
GRHPR RNAi1
Ctrl3
GRHPR RNAi3
H
Mac # on yolk
***
***
Ctrl1
LKRSDH RNAi1
Ctrl2
LKRSDH RNAi2

**Figure S3 related to Figure 3. Macrophage transcriptome analysis reveals that Atos targets participate in signaling, cell communication and ion transport.**

**Fig S3A.** FACS plot of Side Scatter (SSC) vs. mCherry fluorescence signal in macrophages obtained from embryos expressing *srpHemo-3xmCherry*. The two populations are sorted as mCherry marker + (red) and – (blue) cells. **Fig S3B.** Genes expressed differentially in analysis of RNA sequencing data from macrophages from the $atos^{PBG}$ mutant compared to the control are shown in a volcano plot graphing the $\log_{10}$ of the P value against the log fold change (FC) of the mean normalized expression levels. Each point represents the average value of one gene's expression from four replicate experiments. Dotted vertical lines indicate a $\log_{10}$ fold change $\geq 1$ and the dotted horizontal line a *P* value of $\leq 0.05$. Statistically significant up- and down-regulated genes are reported as red and green dots, respectively. **Fig S3C.** Gene ontology (GO) analysis of downregulated genes from $atos^{PBG}$ mutant macrophages compared to the control shows that theses genes are involved in oxidation-reduction processes, stress responses as well as the nervous system. **Fig S3D-E.** Quantification in fixed early Stage 12 embryos reveals that knockdown by two different RNAis of (**D**) *Glycerophosphate oxidase 2* (*Gpo2*, *CG2137* or (**E**) *Golgi matrix protein 130 kD* (*GM130*, CG11061) did not change the macrophage number within the germband compared to their controls. For (**D**) control 1 n=24, *Gpo2 RNAi 1* (VDRC 41234) n=11, p=0.26; control 2 n=15, *Gpo2 RNAi 2* (VDRC 68145) n=27, p=0.38. For (**E**) control 1 n=15, *GM130 RNAi 1* (VDRC 330284) n=25, p=0.14; control 2 n=27, *GM130 RNAi 2* (VDRC 64920) n=20, p=0.34. **Fig S3F-H.** Quantification reveals that expression of RNAis against *porthos, GR/HPR,* and *LKR/SDH* in macrophages leads to a significant increase in macrophage numbers on the yolk in fixed early Stage 12 embryos compared to their controls. For (**F**) control n=30, *porthos* RNAi n=28, p<0.0001. For (**G**) control 1 n=27, *dGR/HPR RNAi 1* (VDRC 44653) n=18, p=0.0003; control 2 n=22, *dGR/HPR RNAi* 2 (VDRC 107680) n=24, p=0.04; control 3 n=14, *dGR/HPR RNAi 3* (VDRC 64652) n=21, p=0.7. For (**H**) control 1 n=27, *dLKR/SDH RNAi 1* (VDRC 51346) n=17, p=0.0002; control 2 n=22, *dLKR/SDH RNAi 2* (VDRC 109650) n=19, p=0.0004. Unpaired t test for (**D**-**H**).

Figure. S4

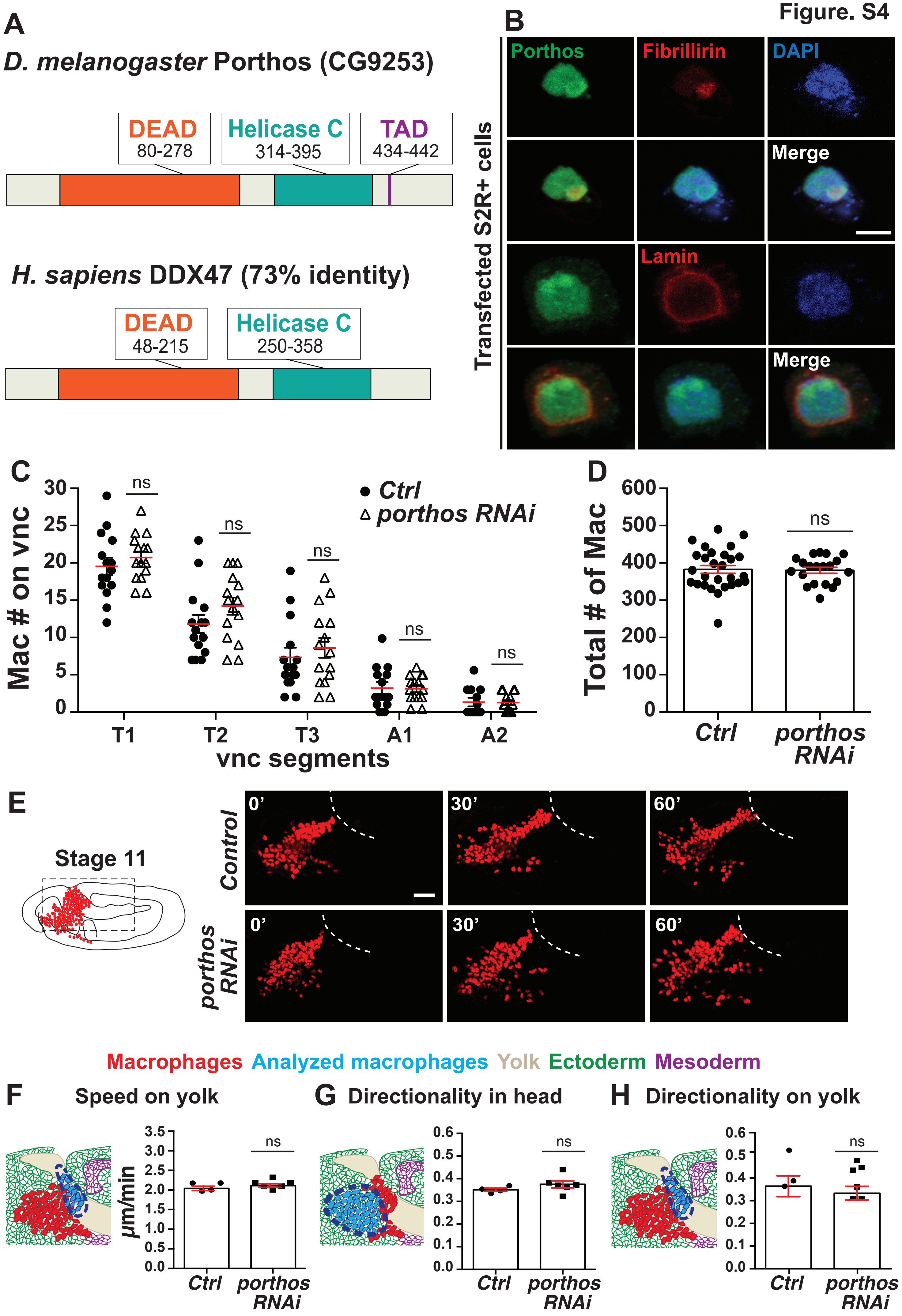
A
D. melanogaster Porthos (CG9253)
DEAD
80-278
Helicase C
314-395
TAD
434-442
H. sapiens DDX47 (73% identity)
DEAD
48-215
Helicase C
250-358
B
Transfected S2R+ cells
Porthos
Fibrillirin
DAPI
Merge
Lamin
Merge
C
Mac # on vnc
30
25
20
15
10
5
0
ns
Ctrl
porthos RNAi
T1
T2
T3
A1
A2
vnc segments
D
Total # of Mac
600
500
400
300
200
100
0
ns
Ctrl
porthos RNAi
E
Stage 11
Control
porthos RNAi
0'
30'
60'
Macrophages Analyzed macrophages Yolk Ectoderm Mesoderm
F
Speed on yolk
μm/min
3.5
3.0
2.5
2.0
1.5
1.0
0.5
0.0
ns
Ctrl
porthos RNAi
G
Directionality in head
0.6
0.5
0.4
0.3
0.2
0.1
0.0
ns
Ctrl
porthos RNAi
H
Directionality on yolk
0.6
0.5
0.4
0.3
0.2
0.1
0.0
ns
Ctrl
porthos RNAi

**Figure S4 related to Figure 4. Downregulation of *porthos* recapitulates the *atos* mutant phenotype.**

**Fig S4A.** Deduced protein structure of Porthos (CG9253). Porthos contains two conserved motifs, a DEAD motif (Asp-Glu-Ala-Asp) and a Helicase C domain, as well as a predicted transactivation domain (TAD). *Drosophila* Porthos shows 71% identity and 84% similarity to its human ortholog, DDX47. **Fig S4B.** Porthos (green) in S2R+ cells transfected with *UAS-porthos::HA* and *srpHemo-Gal4*, and stained for the nuclear membrane marker Lamin (red), colocalizes with the staining for the nucleolar marker Fibrillarin (red), and DAPI (blue). **Fig S4C-D.** Quantification of macrophage numbers in fixed Stage 12 embryos. (**C**) Expression of *porthos RNAi* in macrophages has no effect in their numbers on (**C**) the vnc or (**D**) in the whole embryo compared to the control. For (**C**) control n=15, *porthos RNAi* n=15, p>0.35. For (**D**) control n=28, *porthos RNAi* n=20, p=0.85. **Fig S4E.** Stills from two-photon movies of the migration of macrophages labeled with *srpHemo-H2A::3xmCherry* in control embryos and in those expressing *porthos RNAi* in macrophages. Macrophages from both genotypes have a similar (**F**) directionality in the head, and (**G**) speed and (**H**) directionality on the yolk sac, to control macrophages. Speed on yolk sac: control=2.10 µm/min, *porthos RNAi*=2.15 µm/min; p=0.35; movie #: control n=4, *porthos RNAi* n=6; track #: control n=104, *porthos RNAi* n=168. Directionality in head: control n=0.35, *porthos RNAi* n=0.37; p=0.27; movie #: control n=4, *porthos RNAi* n=6. Directionality on yolk: control=0.42, *porthos RNAi*=0.39; p=0.58; movie #: control n=3, *porthos RNAi* n=6. Unpaired t test for (**C**), (**D**), and (**F-H**). Scale bar is 5 µm in (**B**) and 30 µm in (**E**).

**Figure. S5**

## A

| Biological function | Gene symbol | Description of Porthos targets | Vertebrate ortholog |
|---|---|---|---|
| DNA regulation, Transcription | CG11403 | DNA DEAD/H box helicase 11 | Ddx11 |
| | CG11335 | Lysyl oxidase-like 1 (Loxl1), euchromatinization | Loxl2 |
| | CG10694 | nucleotide-excision repair | Rad23a |
| | CG12659 | Chromatin remodeling | Ino80c |
| | CG5441 | taxi, transcription factor | Atoh1 |
| | CG13005 | Zinc finger protein 839, transcription factor | Zfp839 |
| | CG7963 | Zinc finger C2H2 transcription factor | Gm14322 |
| | CG8021 | SLIRP2, mRNA processing | Slirp |
| | CG8159 | Regulation of transcription | Plag1 |
| | CG11456 | Regulation of transcription by RNA polymerase II | Plagl2 |
| | CG10654 | Regulation of transcription by RNA polymerase II | J23Rik |
| | CG31626 | Regulation of transcription by RNA polymerase II | Pou2af1 |
| | CG12442 | wuc, regulation of transcription by RNA polymerase II | Lin52 |
| | CG12320 | A1 cistron-splicing factor, AAR2 | Aar2 |
| | CG12938 | U7 snRNA-associated Sm-like protein LSm10 | Lsm10 |
| | CG7637 | snRNA/rRNA pseudouridine synthesis | Nop10 |
| RNA translation | CG15693 | RpS20, ribosomal small protein S20 | Rps20 |
| | CG3997 | RpL39, ribosomal large protein L39 | Rpl39l |
| | CG30425 | RpL41, ribosomal large protein L41 | NF |
| | CG4061 | Rtca, RNA 3'-terminal phosphate cyclase | Rtca |
| | CG18643 | Dtd, D-aminoacyl-tRNA deacylase, tRNA metabolic process | Dtd1 |
| Protein degradation | CG8272 | SCF-dependent proteasomal ubiquitin-dependent proteolysis | Lrrc29 |
| | CG14260 | Proteasomal ubiquitin-dependent proteolysis | NF |
| | CG31807 | Ubiquitin-protein transferase | Rfwd3 |
| | CG8419 | Ubiquitin-protein transferase | Trim45 |
| | CG32847 | Ubiquitin-protein ligase | Rnf185 |
| | CG5001 | Chaperone/unfolded protein binding | Dnajb5 |
| | CG2046 | Proteasome assembly chaperone 1 | Psmg1 |
| | CG6972 | Desumoylating isopeptidase 1 | Desi1 |
| Immune cell response | CG2723 | ImpE3, Ecdysone-inducible gene E3 | NF |
| | CG1367 | Cecropin A2, activity against Gram-negative bacteria | NF |
| | CG10794 | Diptericin B, activity against Gram-negative bacteria | NF |
| | CG16712 | IM33 peptide against systemic microbial infection | Eppin |
| | CG33493 | Antibacterial humoral response | Ndufa5 |

## B

| Biological function | Gene symbol | Description of Porthos targets | Vertebrate ortholog |
|---|---|---|---|
| Signal transduction | CG1279 | reticulon 2, ER organization and function | Rtn1 |
| | CG5417 | Srp14, protein targeting to ER | Srp14 |
| | CG12843 | Tetraspanin 42Ei, Integrin signaling | Cd63 |
| | CG5657 | Sarcoglycan β, negative regulator of EGFR pathway | Sgcb |
| | CG3302 | Corazonin, a G-protein-coupled receptor | NF |
| | CG42366 | Mitogen-activated protein kinase | NF |
| | CG8767 | Mos oncogeneactivates the MAPK cascade | Mos |
| | CG9336 | positive regulation of voltage-gated K+ channel | NF |
| | CG3504 | inaD, fast light-induced signaling | Lnx1 |
| | CG7916 | Haemolymph juvenile hormone binding | NF |
| | CG18188 | Damm, caspase family of cysteine proteases | Casp6 |
| | CG9470 | Metallothionein A, metal ion homeostasis | Mt1 |
| | CG3227 | insensitive, corepressor for the product of Su(H) | NF |
| | CG17479 | Sphingosine kinase 1, regulates cell division/traffiking | NF |
| | CG17962 | Z600, a mitotic inhibitor | NF |
| | CG10861 | Autophagy-related 12 | Atg12 |
| | CG14937 | G2/M transition of mitotic cell cycle | NF |
| | CG32812 | negative regulation of phosphatase activity | Chp1 |
| | CG31391 | negative regulation of phosphatase activity | Ppp1r36 |
| Transport | CG17137 | Porin2, voltage-dependent anion channel 1 | Vdac1 |
| | CG7912 | Sulfate transport and transmembrane transport | Slc26a11 |
| | CG18345 | Trpl, transient receptor potential-like | Trpc5 |
| | CG32069 | ER to Golgi vesicle-mediated transpor | Ier3ip1 |
| | CG11703 | Sodium:potassium-exchanging ATPase | Atp1b1 |
| | CG5421 | H(+)-transporting two-sector ATPase | Atp6ap1l |
| Cell-cell interaction | CG13664 | Cadherin 96Cb, control of cell adhesion | Cdh6 |
| | CG16719 | Regulation of cytoskeleton organization | Spef1 |
| | CG5987 | TTLL6B, microtubule cytoskeleton organization | Ttll6 |
| | CG4537 | Cytoplasmic microtubule organization | Cript |
| | CG7802 | Neyo, regulation of cell shape/apical constriction | NF |
| | CG12408 | Troponin C isoform 4, control of muscle contraction | Calm4 |
| | CG8121 | Pasiflora 2 (pasi2), endothelial barrier function | NF |
| | CG5458 | Radial spoke head protein 1, axoneme assembly | Rsph1 |
| | CG31020 | Sanpodo, cell division/cell fate determination | NF |
| | CG31801 | Mst36Fa, spermatogenesis | NF |

**Figure S5 related to Figure 5. Porthos increases the translation of a subset of mRNAs.**

**Fig 5SA-B.** Other target mRNAs downregulated in *porthos KD* cells are involved in gene regulation and RNA processing, mRNA translation, cellular transport, cell signaling, cell-cell interactions, immune responses, and protein degradation. NF: Not Found.

Figure. S6

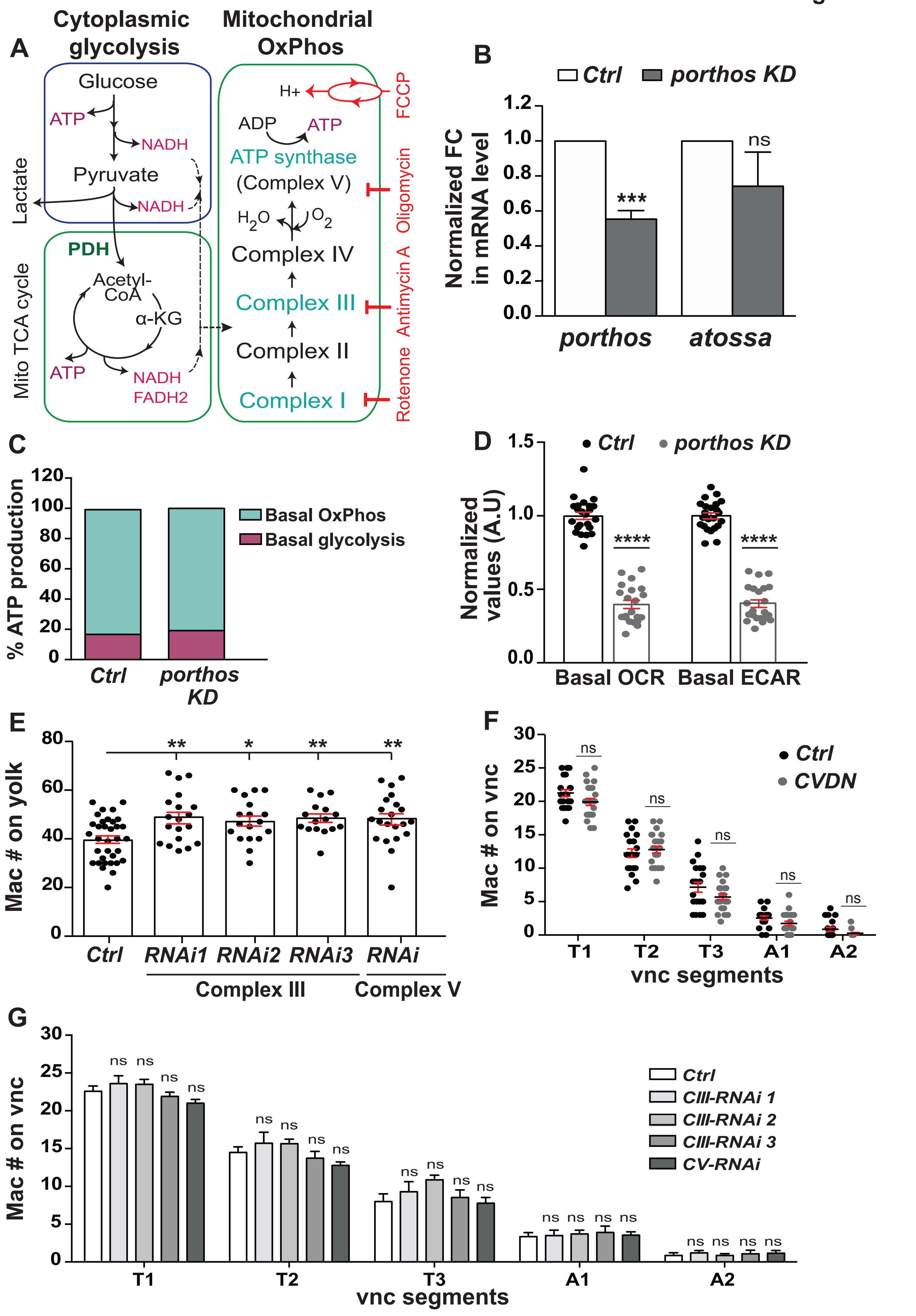
A
Cytoplasmic glycolysis
Mitochondrial OxPhos
Glucose
ATP
NADH
Pyruvate
NADH
Lactate
PDH
Acetyl-CoA
α-KG
ATP
NADH
FADH2
Mito TCA cycle
H+
ADP
ATP
ATP synthase
(Complex V)
H2O
O2
Complex IV
Complex III
Complex II
Complex I
FCCP
Oligomycin
Antimycin A
Rotenone
B
Ctrl
porthos KD
Normalized FC in mRNA level
***
ns
porthos
atossa
C
% ATP production
Basal OxPhos
Basal glycolysis
Ctrl
porthos KD
D
Ctrl
porthos KD
Normalized values (A.U)
****
****
Basal OCR
Basal ECAR
E
Mac # on yolk
**
*
**
**
Ctrl
RNAi1
RNAi2
RNAi3
RNAi
Complex III
Complex V
F
Ctrl
CVDN
Mac # on vnc
ns
T1
T2
T3
A1
A2
vnc segments
G
Mac # on vnc
ns
Ctrl
CIII-RNAi 1
CIII-RNAi 2
CIII-RNAi 3
CV-RNAi
T1
T2
T3
A1
A2
vnc segments

**Figure S6 related to Figure 6. Depletion of *atos* or *porthos* causes impairment in mitochondrial metabolic activity, reduced ATP production, and a deficiency in macrophage tissue invasion.**

**Fig S6A.** Schematic indicating the specific inhibitors (in red at right) used to block the function of mitochondrial OxPhos components. The glycolysis, TCA cycle, and mitochondrial respiratory chain in eukaryotic cells are shown. **Fig S6B.** Graph shows relative *porthos* and *atos* mRNA levels (± SEM) in *porthos KD* S2R+ cells measured by qPCR from at least three independent experiments. The data are normalized to results for the internal control gene RpS20. *Porthos KD* S2R+ cells contain 56% of normal *porthos* mRNA levels and display a slight statistically insignificant decrease in *atos* mRNA levels. t-test was used followed by Sidak's correction. Control n=6, *porthos* n=6, p= 0.0002, *atos* n=3, p=0.09. **Fig S6C.** The contribution of basal OxPhos ATP production rate and glycolytic ATP production rate were calculated. The plot shows that both wild-type and *porthos KD* S2R+ cells utilize OxPhos respiration as the predominant bioenergetic pathway to produce ATP in these cells. Porthos depletion produced no increase in the relative utilization of glycolysis. **Fig S6D.** The relative basal values of the Oxygen Consumption rate (OCR), as a marker of OxPhos, and Extracellular Acidification Rate (ECAR), as an indication of glycolysis, in control and *porthos KD* S2R+ cells are plotted. Basal respiration rate is calculated before the addition of antimycin A. **Fig S6E.** Quantification in fixed early Stage 12 embryos shows a significant increase of macrophages on the yolk upon the expression in macrophages of any of three different *RNAis* against mitochondrial OxPhos *Complex III* (*UQCR)* or an *RNAi* against *Complex V* (*F1F0,* CG3612). Control n=34, *Complex III* (*Cyt-c1,* CG4769): *RNAi 1* (VDRC 109809) n=19, p=0.0049; *Complex III* (*UQCR-cp1,* CG3731): *RNAi 2* (VDRC 101350) n=18, p=0.024; *Complex III* (*UQCR-cp2,* CG4169): *RNAi 3* (VDRC 100818) n=16, p=0.009; *Complex V* (*F1F0, CG3612*): *RNAi* (VDRC 34664) n=21, p=0.0068. **Fig S6F-G.** Quantification of the number of macrophages in vnc segments does not show a significant change in general migration along the vnc in embryos whose macrophages express (**F**) *CV-DN* or (**G**) *RNAis* against mitochondrial OxPhos complex components compared to the control. (**F**): Control n=20, *CV-DN* n=23, p>0.05. **(G):** Control n=14, *Complex III (Cyt-c1,* CG4769): *RNAi 1* (VDRC 109809) n=10, p>0.8; *Complex III* (*UQCR-cp1,* CG3731): *RNAi 2* (VDRC 101350) n=14, p>0.05; *Complex III* (*UQCR-cp2,* CG4169): *RNAi 3* (VDRC 100818) n=11, p>0.9; *Complex V* (*F1F0,* CG3612): *RNAi* (VDRC 34664) n=18, p>0.2. Unpaired t test for (**B**) and (**D-G**).

**Figure. S7**

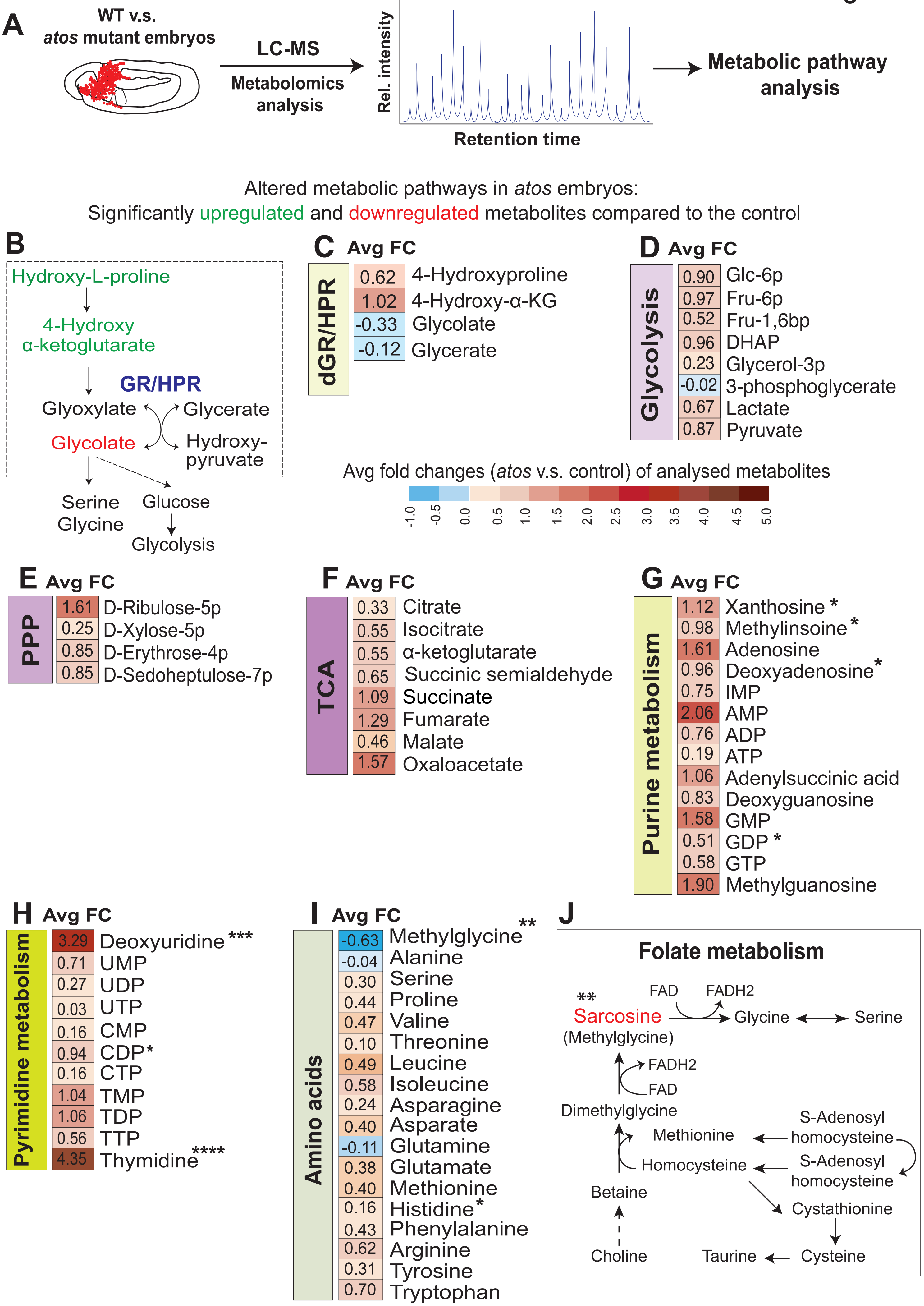
A
WT v.s.
atos mutant embryos
LC-MS
Metabolomics
analysis
Rel. intensity
Retention time
Metabolic pathway
analysis
Altered metabolic pathways in atos embryos:
Significantly upregulated and downregulated metabolites compared to the control
B
Hydroxy-L-proline
4-Hydroxy
α-ketoglutarate
GR/HPR
Glyoxylate
Glycerate
Glycolate
Hydroxy-
pyruvate
Serine
Glycine
Glucose
Glycolysis
C Avg FC
dGR/HPR
0.62 4-Hydroxyproline
1.02 4-Hydroxy-α-KG
-0.33 Glycolate
-0.12 Glycerate
D Avg FC
Glycolysis
0.90 Glc-6p
0.97 Fru-6p
0.52 Fru-1,6bp
0.96 DHAP
0.23 Glycerol-3p
-0.02 3-phosphoglycerate
0.67 Lactate
0.87 Pyruvate
Avg fold changes (atos v.s. control) of analysed metabolites
-1.0 -0.5 0.0 0.5 1.0 1.5 2.0 2.5 3.0 3.5 4.0 4.5 5.0
E Avg FC
PPP
1.61 D-Ribulose-5p
0.25 D-Xylose-5p
0.85 D-Erythrose-4p
0.85 D-Sedoheptulose-7p
F Avg FC
TCA
0.33 Citrate
0.55 Isocitrate
0.55 α-ketoglutarate
0.65 Succinic semialdehyde
1.09 Succinate
1.29 Fumarate
0.46 Malate
1.57 Oxaloacetate
G Avg FC
Purine metabolism
1.12 Xanthosine *
0.98 Methylinsoine *
1.61 Adenosine
0.96 Deoxyadenosine *
0.75 IMP
2.06 AMP
0.76 ADP
0.19 ATP
1.06 Adenylsuccinic acid
0.83 Deoxyguanosine
1.58 GMP
0.51 GDP *
0.58 GTP
1.90 Methylguanosine
H Avg FC
Pyrimidine metabolism
3.29 Deoxyuridine ***
0.71 UMP
0.27 UDP
0.03 UTP
0.16 CMP
0.94 CDP*
0.16 CTP
1.04 TMP
1.06 TDP
0.56 TTP
4.35 Thymidine ****
I Avg FC
Amino acids
-0.63 Methylglycine **
-0.04 Alanine
0.30 Serine
0.44 Proline
0.47 Valine
0.10 Threonine
0.49 Leucine
0.58 Isoleucine
0.24 Asparagine
0.40 Asparate
-0.11 Glutamine
0.38 Glutamate
0.40 Methionine
0.16 Histidine *
0.43 Phenylalanine
0.62 Arginine
0.31 Tyrosine
0.70 Tryptophan
J
Folate metabolism
**
Sarcosine
(Methylglycine)
FAD
FADH2
Glycine
Serine
FADH2
FAD
Dimethylglycine
Methionine
S-Adenosyl
homocysteine
Homocysteine
S-Adenosyl
homocysteine
Betaine
Cystathionine
Choline
Taurine
Cysteine

**Figure S7 related to Figure 7. Atos and Porthos enhance ATP production by programming mitochondrial oxidative phosphorylation metabolism. Fig S7A.** Schematic illustrates the metabolic profiling procedure in wild-type and *atos* mutant embryos at Stage 12. **Fig S7B-C.** Heatmap of non-targeted metabolites in *atos* mutant embryos reveals an increase in substrates of the dGR/HPR enzyme, including 4-hydroxy α-ketoglutarate and hydroxyproline (HLP) and a smaller decrease in its products, glycolate and glycerate. **Fig S7D-F.** Global metabolite screening reveals less than 1 fold increases for most (**D**) glycolytic intermediates and up to 3 fold increases for metabolites from (**E**) the Pentose Pathway (PPP), and (**F**) the TCA cycle in the *atos* mutant compared to the control. **Fig S7G-H.** Analysis reveals strong increases in thymidine, which can be catabolized to products that feed into the TCA cycle, as well as uridine along with increases in some cellular nucleotide precursors and purine and pyrimidine metabolites. **Fig S7I.** Heatmap of non-targeted metabolites in *atos* mutant embryos reveals a small increase in most amino acids in the *atos* mutant a significant increase in some dipeptides including those containing hydroxyproline. **Fig S7J.** Schematic shows a link between Folate metabolism and glycine/serine metabolism, in which the glycine-related metabolite sarcosine (N-methylglycine) was significantly reduced in the *atos* mutant. Metabolites with statistical significant change are shown as: *$p<0.05$, **$p<0.01$, ***$p<0.001$, ****$p<0.0001$. Unpaired t test for (**C-I**).
