## Supplementary material for "A genetic program boosts mitochondrial function to power macrophage tissue invasion": Key resource table

**Table S1.** Fly lines utilized in this paper.

| Experimental models: Organisms/Strains | | | |
|---|---|---|---|
| **Designation** | **Source of reference** | **Identifiers** | **Additional information** |
| *srpHemo-Gal4* | PMID: 15239955 | Brückner et al., 2004 | *D. melanogaster* |
| *srpHemo-3xmCherry* | PMID: 29321168 | RRID:BDSC_78358 and 78359 | *D. melanogaster* (Gyoergy et al., 2018) |
| *srpHemo-H2A::3xmCherry* | PMID: 29321168 | RRID:BDSC_78360 and 78361 | *D. melanogaster* (Gyoergy et al., 2018) |
| *CG9005$^{BG02278}$* | Bloomington *Drosophila* Stock Center (BDSC) | RRID:BDSC_12768 | |
| *Df(2R)ED2222 (Df1)* | BDSC 8911 | | |
| *Df(2R)BSC259 (Df2)* | BDSC 23159 | | |
| *UAS-CG9005 RNAi 1* | VDRC | VDRC: v106589 | |
| *UAS-CG9005 RNAi 2* | VDRC | VDRC: v36080 | |
| *UAS-CG9005 RNAi 3* | BDSC | BDSC: 33362 | |
| *srpHemo-HA::CG9005 (srpHemo-HA::atossa)* | this paper | | CG9005 amplified from genome cloned into DSPL172 (PMID: 29321168) |
| *srpHemo-HA::atossa$^{nls1-}$* | this paper | | CG9005 amplified from genome cloned into DSPL172 |
| *srpHemo-HA::atossa$^{DUF4210-}$* | this paper | | CG9005 amplified from genome cloned into DSPL172 |
| *srpHemo-HA::atossa $^{ChrSeg-}$* | this paper | | CG9005 amplified from genome cloned into DSPL172 |
| *srpHemo-HA::atossa $^{DUF4210-/ChrSeg-}$* | this paper | | CG9005 amplified from genome cloned into DSPL172 |
| *srpHemo-HA::atossa$^{TAD1-}$* | this paper | | CG9005 amplified from genome cloned into DSPL172 |
| *srpHemo-HA::atossa$^{TAD2-}$* | this paper | | CG9005 amplified from genome cloned into DSPL172 (PMID: 29321168) |
| *srpHemo-HA::atossa$^{TAD1-/TAD2-}$* | this paper | | CG9005 amplified from genome cloned into DSPL172 |
| *srpHemo-FAM214A* | this paper | | FAM214A amplified from dendritic cell cDNA library cloned into *srpHemo* plasmid (DSPL172) |
| *srpHemo-FAM214B* | this paper | | FAM214B amplified from dendritic cell cDNA library |

| | | | |
|---|---|---|---|
| | | | cloned into *srpHemo* plasmid (DSPL172) |
| *UAS-HA::EGFP* | this paper | | |
| *UAS-CG9253 RNAi* (*porthos*) | VDRC, RRID: | VDRC: v36589 | |
| *UAS-CG9331 RNAi 1* (*GRHPR*) | (VDRC), RRID: | VDRC: v44653 | |
| *UAS-CG9331 RNAi 2* (GRHPR) | BDSC, RRID: | BDSC: 64652 | |
| *UAS-CG9331 RNAi 3* (*GRHPR*) | (VDRC), RRID: | VDRC: v107680 | |
| *UAS-CG7144 RNAi 1* (*LKRSDH*) | (VDRC), RRID: | VDRC: v51346 | |
| *UAS-CG7144 RNAi 2* (*LKRSDH*) | (VDRC), RRID: | VDRC: v109650 | |
| *UAS-CG2137 RNAi 1* (*Gpo2*) | (VDRC), RRID: | VDRC: v1234 | |
| *UAS-CG2137 RNAi 2* (*Gpo2*) | BDSC, RRID: | BDSC: 68145 | |
| *UAS-CG11061 RNAi* (*GM130*) | BDSC, RRID: | BDSC: 64920 | |
| *UAS-CG11061 RNAi* (*GM130*) | (VDRC), RRID: | VDRC: v330284 | |
| *y[-] v[-];attP40-pVALIUM22-UAS-ATPsyn Subunit C* (*CG1746*) *E121Q* | (VDRC), RRID: | Thomas Hurd, et al., 2016 | |
| *UAS-CG4769 RNAi 1 (Cyt-c1)* | (VDRC), RRID: | VDRC: *v109809* | |
| *UAS-CG4169 RNAi 2 (UQCR-cp2)* | (VDRC), RRID: | VDRC: *v100818* | |
| *UAS-CG3731 RNAi 3 (UQCR-cp1)* | (VDRC), RRID: | VDRC: *v101350* | |
| *UAS-CG3612 RNAi (ATP synthase F1F0)* | (VDRC), RRID: | VDRC: *v34664* | |

**Table S2:** List of key resources used in this paper.

| Antibodies | | | |
|---|---|---|---|
| **Designation** | **Source of reference** | **Identifiers** | **Additional information** |
| Chicken polyclonal anti-GFP | Aves Labs | Cat# GFP-1020, RRID:AB_10000240 | |
| Rat monoclonal anti-HA | Roche | Roche Cat# 3F10, RRID: AB_2314622 | |
| Mouse Lamin (lamin Dm0) | *Drosophila* Studies Hybridoma Bank (DSHB) | Cat# ADL1010 | |
| Mouse Fibrillarin | Rangan lab | N/A | |
| Mouse anti-Pyruvate Dehydrogenase E1-alpha subunit antibody (PDH E1α) [8D10E6 ] | Abcam | Cat# ab110334, RRID:AB_10866116 | |
| Rabbit antiphoshpo-Pyruvate Dehydrogenase E1-alpha subunit (PDH E1α, S293) | Abcam | Cat# ab92696, RRID:AB_10711672 | |
| Goat anti-Chicken IgY (H+L) Secondary Antibody, Alexa Fluor 488 | Thermo Fisher Scientific | Cat# A-11039, RRID: AB_2534096 | |
| Alexa Fluor 488 goat anti-rat | Thermo Fisher Scientific | Cat# A21212, RRID: AB_11180047 | |
| Goat anti-Mouse IgG1 Secondary Antibody, Alexa Fluor 488 conjugate | Thermo Fisher Scientific | Cat# A-21121, RRID: AB_2535764 | |
| Goat anti-Mouse IgG2b Secondary Antibody, Alexa Fluor 633 conjugate | Thermo Fisher Scientific | Cat# A-21146, RRID:AB_2535782 | |
| Goat anti-Rabbit IgG (H+L) Secondary Antibody, Alexa Fluor 488 conjugate | Thermo Fisher Scientific | Cat# R37116, RRID: AB_2556544 | |
| Phalloidin 488 | Thermo Fisher Scientific | Cat# A12379, RRID:AB_2315147 | |

| | | |
|---|---|---|
| Phalloidin 633 | Thermo Fisher Scientific | Cat# 50-6559-05, RRID:AB_2574272 |
| **Chemicals** | | |
| Vectashield mounting medium | Vector Laboratories, RRID:SCR_000821 | VectorLabs: H-1000 |
| Vectashield Mounting medium with DAPI | Vector Laboratories, RRID:SCR_000821 | VectorLabs: H-1200 |
| Beckman Coulter 9/16x3.5 PA tubes | | Cat. #331372 |

| | | |
|---|---|---|
| **Critical Commercial Assays** | | |
| Infusion cloning kit | Clontech's European distributor | Cat# A14606 |
| MEGAscript_ T7 Transcription Kit | Thermo Fisher Scientific | Cat# AM1334 |
| MEGAscript_ T3 Transcription Kit | Thermo Fisher Scientific | Cat# AM1338 |
| Effectene Tranfection Reagent kit | Qiagen, Hilden, Germany | |
| DNeasy Blood & Tissue Kit | Qiagen, Hilden, Germany | |
| QIAGEN Rneas Mini Kit | Qiagen, Hilden, Germany | Cat#74104 |
| Takyon™ No Rox SYBR MasterMix blue dTTP | Eurogentec, Liege, Belgium | |
| TURBO DNA-free Kit | Life Technologies | Cat# AM1907 |
| Agilent Seahorse XF Cell Mito Stress Test kit | Agilent Technologies, Inc., Santa Clara, CA, USA | Cat# 103015-100 |
| Agilent 6000 Pico kit | Agilent Technologies, Waldbronn, Germany | Cat# 5067-1513 |

**Table S3.** The DNA plasmid constructs utilized in gene construction.

| Recombinant DNA | | | |
|---|---|---|---|
| **Designation** | **Source of reference** | **Identifiers** | **Additional information** |
| *UAS-CG9005::FLAG::HA* | *Drosophila* Genomics Resource Center | DGRC: UFO03339 Flybase: FBgn0033638 | *atossa* |
| *UAS-CG9253::FLAG::HA* | *Drosophila* Genomics Resource Center | DGRC: UFO12394 Flybase: FBgn0032919 | *porthos* |
| *UAS-CG9331::FLAG::HA* | *Drosophila* Genomics Resource Center | DGRC: UFO02643 Flybase: FBgn0032889 | Glyoxylate reductase (NADP(+)) Hydroxypyruvate reductase (GR/HPR) |
| *UAS-CG7144::FLAG::HA* | *Drosophila* Genomics Resource Center | DGRC: UFO05689 Flybase: FBgn0286198 | Lysine ketoglutarate reductase/saccharopine dehydrogenase (LKRSDH) |
| *pAC-sgRNA-Cas9* | Addgene | Addgene: 49330 | 49330 (DSPL 232) |

**Table S4.** Oligonucleotides utilized in gene construction.

| No. | Name | Sequence |
|---|---|---|
| 1 | FP-CG9005 | TAGAAGCTTCTGCAAATGATACCGACAAGCGTCACC |
| 2 | RP-CG9005 | GTGCCTAGGCGCGCCCTAAATCCTGCCGGCGCT |
| 3 | FP-HACG9005 | TAGAAGCTTCTGCAAATGTACCCATACGATGTTCCAGATTAC GCTGCCGCCGCCATGATACCGACAAGCGTCACC |
| 4 | RP-HACG9005 | GTGCCTAGGCGCGCCAGCGTAATCTGGAACATCGTATGGGT AGGCGGCGGCAATCCTGCCGGCGCTCTC |
| 5 | infFPCG9005_NotIBluS | ACCGCGGTGGCGGCCATGTACCCATACGATGTTCCAG |
| 6 | infRPCG9005_NotIBluS | CGAAGTTATGCGGCCCTAAATCCTGCCGGCGCTC |
| 7 | FP-DUF4210$^-$CG9005 | TTGTGCGAGATTCGTTTGCCG |
| 8 | RP-DUF4210$^-$CG9005 | AACGGACGTCCTCCAAATTGAG |
| 9 | FP-ChrSeg$^-$CG9005 | AGTGCGCGACAGGAGAGC |
| 10 | RP- ChrSeg -CG9005 | AGTCGCTTCATCTGCTCGG |
| 13 | FP-FAM214A-V13 | ATGAAGCCAGACCGAGATGC |
| 14 | RP-FAM214A | TCAACATCTTGGTGAAAACTGAG |
| 15 | infFP-FAM214A-V13 | TAGAAGCTTCTGCAAATGAAGCCAGACCGAGATGC |
| 16 | infRP-FAM214A | GTGCCTAGGCGCGCCTCAACATCTTGGTGAAAACTG |
| 17 | PF-FAM214B | GGCTTCATGCGCCACGTG |
| 18 | RP-FAM214B | CGATCAGGGCAAAGGTGAATAACG |
| 19 | infFP-FAM214B | TAGAAGCTTCTGCAAGGCTTCATGCGCCACGTG |
| 20 | infRP-FAM214B | GTGCCTAGGCGCGCCCGATCAGGGCAAAGGTGA |
| 21 | Insitu-CG9005 FP1 | CCTCCTTGGGCTCGGCTACTGC |
| 22 | Insitu-CG9005 RP1 | GATAATACGACTCACTATAGGGTTGACGTTGGGAAAATT |
| 23 | Insitu-CG9005 RP2 | GATAATACGACTCACTATAGGGTTTGCAAAGTTGTGCT |
| 24 | Insitu-CG9253 FP1 | GGAAAGATCTCGGTCTCAATGAG |
| 25 | Insitu-CG9253 RP1 | GATAATACGACTCACTATAGGGCCATCACCTCATCCTCC |
| 26 | sgRNA-F1-CG9005 | TTCG GCAGTCGGATGTCCGTATGCAGG |
| 27 | sgRNA-R1-CG9005 | AACGCATACGGACATCCGACTGC C |
| 28 | sgRNA-F2-CG9005 | TTCGCAGTTCGTAGAAGTAAGAGACGG |

| | | |
|---|---|---|
| 29 | sgRNA-R2-CG9005 | AACTCTCTTACTTCTACGAACTG C |
| 30 | sgRNA-F3-CG9005 | TTCGCGGCGGATTCTGTCCCACCCAGG |
| 31 | sgRNA-R3-CG9005 | AACGGGTGGGACAGAATCCGCCG C |
| 32 | sgRNA-F1-CG9253 | TTCGGATCCAACGTGAGGCCATTCCGG |
| 33 | sgRNA-R1-CG9253 | AACGAATGGCCTCACGTTGGATC C |
| 34 | sgRNA-F2-CG9253 | TTCGGGCCATTCCGGTCGCCTTACAGG |
| 35 | sgRNA-R2-CG9253 | AACGTAAGGCGACCGGAATGGCC C |
| 36 | sgRNA-F3-CG9253 | TTCGCCCTCGTGGGGGTTAGCACGAGG |
| 37 | sgRNA-R3-CG9253 | AACCGTGCTAACCCCCACGAGGG C |
| 38 | infNotI-TCHA-EGFPHA-FP | AACAGATCTGCGGCCGCATGTGTTGCCCGGGCTGCTGT |
| 39 | infNotI-TCHA-EGFP-RP | CCTCGAGCCGCGGCCGCTTAAGCGTAATCTGGCACATC |
| 40 | CG9005qPCR-FP1 | TGTTCAGATTCTCGCCACCA |
| 41 | CG9005qPCR-RP1 | TGAGGATTTGCCCAGCTGTT |
| 42 | CG9005qPCR-FP2 | GCACGCCTTATTTGTGCGAG |
| 43 | CG9005qPCR-RP2 | CCCGCATGTCGTAGGGTATC |
| 44 | CG9005qPCR-FP3 | TATGCGGCAGGGAGAAAGTT |
| 45 | CG9005qPCR-RP3 | GTGGTCTCTTCTGTCCACCG |
| 46 | CG9253qPCR-FP1 | GCCTTACAGGGCAAGGATGT |
| 47 | CG9253qPCR-RP1 | ATGCCAATCCCGCTACCAAG |
| 48 | CG9253qPCR-FP2 | TCTAGGTAGCGAGGAGGAGC |
| 49 | CG9253qPCR-RP2 | TGGCCTCACGTTGGATCTTC |
| 50 | CG9253qPCR-FP3 | TTCGACCACGTGCTGCTATT |
| 51 | CG9253qPCR-RP3 | TTGTAGCTGCGTCTGTTCGT |
| 52 | RpL32 qPCR-FP | AGCATACAGGCCCAAGATCG |
| 53 | RpL32 qPCR-RP | TGTTGTCGATACCCTTGGGC |
| 54 | RpS20 qPCR-FP | ACGGTGCAAAGAACCAGAACT |
| 55 | RpS20 qPCR-RP | GGAGTCTTACGGGTGGTGATG |
| 56 | pAC-sgRNA-Cas9-U6F | TTTGATTCTAAAGGAAATTTTGAAAA |

**Table S5**. List of software tools, analytical packages, and laboratory devices utilized in this paper.

| Software and Algorithms | | |
|---|---|---|
| **Designation** | **Source of reference** | **Identifiers** |
| ImageJ/FIJI | | http://fiji.sc/ RRID:SCR_002285) |
| Imaris | Bitplane | http://www.bitplane. com/imaris/imaris, RRID:SCR_007370 |
| Matlab | Mathworks | https://www.math works.com/products /matlab.html, RRID:SCR_001622 |
| FlowJo | | https://www.flowjo .com/RRID:SCR_008520 |
| LaVision ImSpector | LaVision BioTec | http://www.lavision biotec.com/, RRID:SCR_015249 |
| Proteome Discoverer 1.4 | | https://www.thermo fisher.com/order/ catalog/product/ OPTON-30795, RRID:SCR_014477 |
| LightCycler 480 software (v. 1.5) | Roche Diagnostics | https://lifescience.roche .com/en_at/products/ lightcycler14301-480 -software-version-15.html |
| 9 aaTAD Prediction Tool | | https://www.med.muni.cz/9aaTAD/analysis.php#matches |
| Conserved Domain Architecture Retrieval Tool (CDART) program | | https://www.ncbi.nlm.nih.gov/Structure/8exington/8exington.cg i |
| Conserved Domain Database (CDD) | | https://www.ncbi.nlm.nih.gov/Structure/cdd/wrpsb.cgi |
| Prism | GraphPad | https://www.graphpad.com/ scientific-software/prism/ RRID:SCR_002798 |
| Flyrnai | sgRNA design | https://www.flyrnai.org/crispr/ http://tools.flycrispr.molbio.wisc.edu/targetFinder/ |
| NCBI primer design tool | Primer design | https://www.ncbi.nlm.nih.gov/tools/primer-blast/ |
| Infusion primer tool | Clontech website | http://bioinfo.clontech.com/infusion/convertPcrPrimersInit.do |
| HISAT2 | | https://ccb.jhu.edu/software/hisat2/index.shtml Kim et al., 2015 |
| MEME Suite | | http://meme-suite.org/doc/overview.html Bailey et al., 2009 |
| Homer (v4.10.4) | | http://homer.ucsd.edu/homer/ |

| Others | | |
|---|---|---|
| **Designation** | **Source of reference** | **Identifiers** |
| Nikon Eclipse Ti Inverted widefield Microscope | Nikon | https://www.nikoninstruments.com/en_EU/Products/Inverted-Microscopes/Eclipse-Ti-E |
| Zeiss LSM 800 Confocal Microscope | Zeiss | https://www.zeiss.com/microscopy/us/products/confocal-microscopes.html |
| LaVision 2-Photon Inverted Microscope | LaVision BioTec | http://www.lavisionbiotec.com/products/trim-scope-ii-1.html |
| YSI Stretch membranes | YSI | https://www.ysi.com/Accessory/id-066155/Membranes-10-Pack-Standard |
| LightCycler 480 | Roche Diagnostics | Idaho Technology Inc., Salt Lake City, UT, USA. |
| FACS Aria III (BD) flow cytometer | | |
| Leica SP8 FALCON inverted confocal | WLL, FALCON, Leica | https://www.leica-microsystems.com/products/confocal-microscopes/p/dive/ |
| Beckman L7 ultracentrifuge | Beckman Coulter, Krefeld, Germany | |
