## Supplementary material for "A genetic program boosts mitochondrial function to power macrophage tissue invasion": Table 1

**Table 1.** Expression of FAM214A and FAM214B genes, the vertebrate orthologs of *Drosophila* Atossa, in vertebrate human immune cells.

| Gene | Immune Cells w/highest level | Description | Expression data | Source |
|---|---|---|---|---|
| **FAM214A** | Plasmacytoid dendritic cells (DCs) | Human | RNA Seq | **The Human Protein Atlas** https://www.proteinatlas.org/ENSG00000047346-FAM214A/blood |
| | Dendritic cells (DC.DC6.123+.Bl) | Human population Medium level expression | Population RNA Seq | **Immune Cell Atlas** http://immunecellatlas.net/ICA_Skyline.php?gene=FAM214A&celltype=all&organ=Blood&datatype=rnaseq&scale=Local |
| | Plasma B cells (B.PC) | High expression | RNA Seq | **Immgen** http://rstats.immgen.org/Skyline/skyline.html |
| | Regulatory T cells (Cd4+, Cd25+) | Human | Microarray | **BioGPS** http://biogps.org/#goto=genereport&id=56204 |
| **FAM214B** | Neutrophils | Human | RNA Seq | **The Human Protein Atlas** https://www.proteinatlas.org/ENSG00000005238-FAM214B/blood |
| | Blood monocytes (Mo.16+.Bl, CD16+) | Human population Medium level expression | Population RNA Seq | http://immunecellatlas.net/ICA_Skyline.php?gene=FAM214B&celltype=all&organ=Blood&datatype=rnaseq&scale=Local |
| | Neutrophils Thio-induced pertonial neutrophils (GN.Thio.PC) | High expression | RNA Seq | **Immgen** http://rstats.immgen.org/Skyline/skyline.html |
| | Neutrophils | Human | Microarray | **BioGPS** http://biogps.org/#goto=genereport&id=80256 |
